# Genetic and Environmental Contributions to Epigenetic Aging Across Adolescence and Young Adulthood

**DOI:** 10.1101/2024.06.10.598273

**Authors:** Dmitry V. Kuznetsov, Yixuan Liu, Alicia M. Schowe, Darina Czamara, Jana Instinske, Charlotte K. L. Pahnke, Markus M. Nöthen, Frank M. Spinath, Elisabeth B. Binder, Martin Diewald, Andreas J. Forstner, Christian Kandler, Bastian Mönkediek

## Abstract

**Background:** Epigenetic aging estimators commonly track chronological and biological aging, quantifying its accumulation (i.e., epigenetic age acceleration) or speed (i.e., epigenetic aging pace). Their scores reflect a combination of inherent biological programming and the impact of environmental factors, which are suggested to vary at different life stages. The transition from adolescence to adulthood is an important period in this regard, marked by an increasing and, then, stabilizing epigenetic aging variance. Whether this pattern arises from environmental influences or genetic factors is still uncertain.

This study delves into understanding the genetic and environmental contributions to variance in epigenetic aging across these developmental stages. Using twin modeling, we analyzed four estimators of epigenetic aging, namely Horvath Acceleration, PedBE Acceleration, GrimAge Acceleration, and DunedinPACE, based on saliva samples collected at two timepoints approximately 2.5 years apart from 976 twins of four birth cohorts (aged about 9.5, 15.5, 21.5, and 27.5 years at first and 12, 18, 24, and 30 years at second measurement occasion).

**Results:** Half to two-thirds (50–68%) of the differences in epigenetic aging were due to unique environmental factors, indicating the role of life experiences and epigenetic drift, besides measurement error. The remaining variance was explained by genetic (Horvath Acceleration: 24%; GrimAge Acceleration: 32%; DunedinPACE: 47%) and shared environmental factors (Horvath Acceleration: 26%; PedBE Acceleration: 47%). The genetic and shared environmental factors represented the primary sources of stable differences in corresponding epigenetic aging estimators over 2.5 years.

Age moderation analyses revealed that the variance due to individually-unique environmental sources was smaller in younger than in older cohorts in epigenetic aging estimators trained on chronological age (Horvath Acceleration: 47% to 49%; PedBE Acceleration: 33% to 68%). The variance due to genetic contributions, in turn, potentially increased across age groups for epigenetic aging estimators trained in adult samples (Horvath Acceleration: 18% to 39%; GrimAge Acceleration: 24% to 43%; DunedinPACE: 42% to 57%).

**Conclusions:** Transition to adulthood is a period of the increasing variance in epigenetic aging. Both environmental and genetic factors contribute to this trend. The degree of environmental and genetic contributions can be partially explained by the design of epigenetic aging estimators.

## Background

Estimators of epigenetic aging offer insights into how individuals age over time and across age groups [28]. Their increased estimates are associated with age-related diseases and mortality in adults [43, 57], and with worse cognitive functioning [58] and more socioeconomic disadvantage in adolescents and children [59]. By measuring DNA methylation (DNAm) levels at specific DNA loci, epigenetic aging estimators address, along universal, specific aspects of aging processes.

Understanding epigenetic aging measures and the aging processes underlying them requires considering their training models with a variable number of CpGs, training markers, and age ranges. The Horvath clock, for example, was trained to predict chronological age in samples with a large age range. It considers the discrepancy between chronological and epigenetic age to indicate accelerated or decelerated aging across the whole life span [27]. Similarly, in buccal samples of children and adolescents, the pediatric-buccal-epigenetic (PedBE) clock illuminates the specific aging processes in younger age groups [47]. Alternatively, epigenetic aging estimators have been developed on the basis of links between aging and biological health indicators. Two examples of such measures are the biomarker of human mortality risk GrimAge [40, 41] and the biomarkers of the pace of aging, DunedinPoAm and DunedinPACE [4, 5].

These measures account for DNAm at locus-specific sites associated, for example, with plasma proteins and smoking (GrimAge) or biomarkers of organ systems integrity, such as blood pressure or serum leptin levels (Dunedin). In addition, Dunedin measures are trained on longitudinal data and represent, in contrast to the degree of epigenetic aging, the pace of epigenetic aging [4, 5].

The strength of the prediction by epigenetic aging measures stems from their ability to represent both genetically driven aging processes and the cumulative effects of the environment on DNAm [35]. The contributions of genetic factors to the variance in epigenetic aging can be reflected in the narrow– and broad-sense *heritability,* the proportion of the phenotypic variance attributable to genetic differences at the time and age of measurement in a certain population.

Narrow-sense heritability quantifies only the contribution of additive genetic factors, whereas broad-sense heritability also includes the interactions between genetic factors such as allelic dominance (i.e., interactions within specific gene loci of chromosomes) and epistasis (i.e., interactions across genetic variants between gene loci).

Figure 1 summarizes previously reported estimates of SNP-^1^, pedigree-^2^, and twin-based^3^ heritability for the aforementioned epigenetic aging measures (see also Additional file 1: Table S1). As shown in Figure 1, the narrow-sense heritability of the Horvath clock measured in blood is not constant across particular ages. According to previous studies, heritability estimates vary from 39% to 61% in older adults, which is on average lower than the estimates obtained for young adults, ranging from 60% to 77% [26, 27, 28, 33, 48, 66]. Further, prior research revealed a lower heritability of the Horvath clock in adolescent than in young adult samples, with a range from 37% to 43% [43, 67]. However, the estimates in the adolescent samples were derived from pedigree– and SNP-based studies, whereas the estimates in the young adult samples were derived from twin studies, which typically result in higher heritability estimates. Given that the twin-based heritability of the Horvath clock appears to peak at birth [27], the lower heritability estimates for adolescence may potentially represent missing heritability besides the age-associated pattern [78].

**Figure 1.**
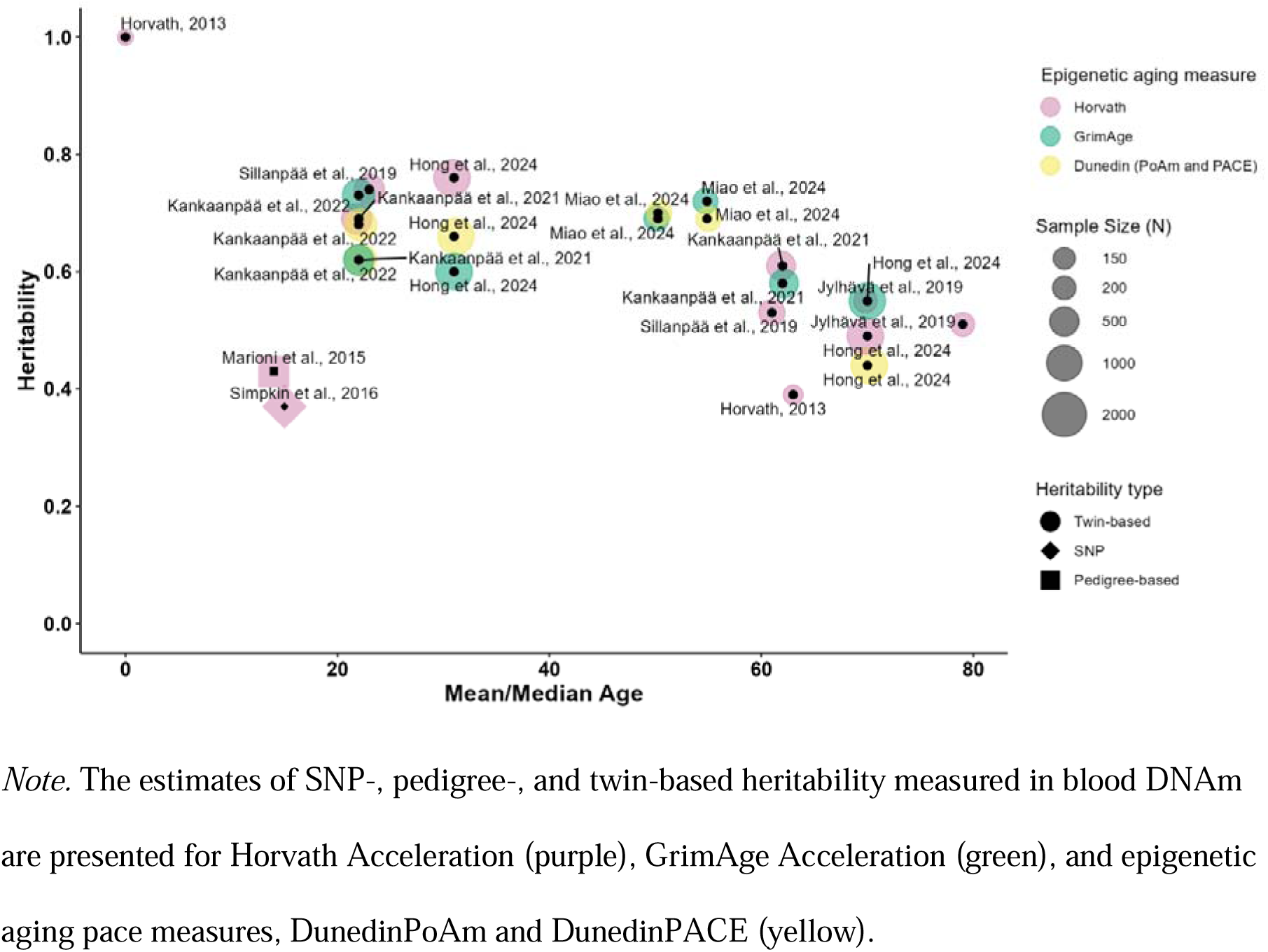
The SNP-, pedigree-, and twin-based heritability of epigenetic aging estimated in previous studies.

The possibility of extrapolating the potential age-associated trend in the heritability of the Horvath clock to other epigenetic aging measures has just started to be explored. For example, heritability comparisons between epigenetic aging estimators are confined mainly to emerging adults [34] or older individuals [26]. In the sample of emerging adults, the Horvath clock, GrimAge, and pace of aging demonstrated similar heritability estimates of 62–73% [34]. Later in life, the Horvath clock, in comparison to GrimAge and DunedinPACE, demonstrated higher estimates until age 66 [26]. Knowledge on the possible comparability of heritability estimates between tissues and resources of DNAm is mainly limited to the comparison of blood tissue and postmortem brain, with lower estimates for the latter (Additional file 1: Table S1). Studying the age-associated patterns of heritability across a broader age range and spectrum of epigenetic aging measures that vary in training ages and phenotypes, therefore, would allow to elucidate trends that are common and specific to certain epigenetic aging measures. Furthermore, DNAm based on saliva can be assumed to correlate with DNAm based on both blood and buccal cells [9]. The examination of the age-associated patterns of heritability based on DNAm derived from noninvasive saliva samples, therefore, may also be indicative of cross-tissue accuracy and tissue-specific differences in heritability.

The trend of epigenetic aging heritability across particular ages may arise from environmental sources and their interaction with genetic factors. A design of twins reared together allows to disentangle environmental factors shared by twins, which increase twin siblings’ similarity, from non-shared or individually-unique environmental factors, which increase twin siblings’ dissimilarity in DNAm. Particularly in younger age, when twins live together in one household, shared environments may be more important. These environments can encompass, for instance, prenatal factors, social backgrounds, common parental and peer influences, and shared neighborhoods. Research on complex traits suggests that shared environmental and genetic factors may interact, particularly in younger individuals [6, 32]. With respect to the variance in epigenetic aging, the role of shared environmental factors was found to increase before adulthood when twins lived together [37]. Individually-unique environmental factors are assumed to be important across the whole life span and can obtain greater importance when twins live in different households and have more life experiences of their own. These environments can include non-shared perinatal factors, stochastic changes in life circumstances, and individual-specific experiences, such as different peer groups or differences in experiences of objectively shared environments. Individually-unique environmental factors have been shown to moderate the influences of genetic factors on DNAm [22] and possibly have the same effect on DNAm-based epigenetic aging during adulthood [30]. These shifts in genetic and environmental contributions to the variance in epigenetic aging over the life course suggest a possible link between life stage processes and genelenvironment interplay in epigenetic aging.

Adolescence and the transition to adulthood, so-called emerging adulthood, are potential periods of dynamic genetic and environmental contributions to the variance of epigenetic aging. The time before transition is a period of growth and development with a faster change rate in DNAm and epigenetic age [1, 28]. Studies have shown, for instance, that this change rate peaks in infancy, decreases nonlinearly until approximately age 20, and then stabilizes [27, 28].

Variance in epigenetic aging can also undergo changes. Starting small at birth, the variance in the epigenetic age of the Horvath clock increases with chronological age until adulthood onset, and then stabilizes [37]. It remains unclear whether these trends of later stabilization in change rates correspond to trends in the relative contributions of genetic and non-genetic factors and which sources, genetic or environmental, are responsible for the growing and stabilizing variance of epigenetic aging during adolescence and the transition to adulthood.

To address these gaps in the epigenetic literature, this study aimed to investigate the trends in genetic and environmental contributions to the variance in epigenetic aging across adolescence, emerging, and young adulthood. With our design, which includes four twin birth cohorts and saliva-based DNAm from two measurement occasions, it is possible to examine how genetic and environmental sources contribute to the variance in epigenetic aging and how these sources shift or remain stable across age groups and over time (approximately two and a half years in this study). DNAm was obtained from DNA extracted from saliva samples of early adolescent (*M*(age) = 10.7 years), late adolescent (*M*(age) = 16.6 years), emerging adult (*M*(age) = 22.7 years), and young adult (*M*(age) = 28.8 years) twins. The epigenetic aging estimators included two acceleration measures trained on chronological age, the Horvath clock [27] and the PedBE clock [47], and two epigenetic aging measures trained on biological indicators of aging, one of which, GrimAge [40], represents biological age acceleration and the other, DunedinPACE, represents the pace of biological aging [5].

With the use of these epigenetic aging estimators, we were able to answer the following research questions: (a) Do genetic and environmental contributions to the variance differ between differently developed epigenetic aging estimators? (b) To what extent do genetic and environmental contributions account for stable differences in epigenetic aging estimators across two measurement occasions? (c) Are there age-associated trends in the genetic and environmental contributions to the variance in epigenetic aging from early adolescence to young adulthood and do they differ for different epigenetic aging estimators? In addition, the findings allow comparisons of heritability estimates of epigenetic aging in young adults assessed using saliva samples with those that have previously been reported in other studies based on blood samples (see Figure 1 and Additional file 1: Table S1).

## Methods

### Sample

The research questions were examined in a sample of 976 twins from the TwinLife Epigenetic Change Satellite (TECS) project, which is a subsample of the German twin family panel TwinLife. TwinLife is a longitudinal study of four cohorts of same-sex twin pairs and their families recruited as a representative sample from the general population of Germany [15]. The study focuses on longitudinal measures of social inequality and the potential underlying biopsychosocial mechanisms [25]. The current analysis sample consisted of 263 monozygotic (MZ) and 225 dizygotic (DZ) complete twin pairs aged 8 to 29 years at the time of the first epigenetic measurement (*M* = 16.0) and aged 10 to 31 years at the second measurement (*M* = 18.3). Zygosity was confirmed by genotyping. The age difference between cohorts was approximately 6 years. At the time of assessment, members of the four cohorts were in the ages of early adolescence, late adolescence, emerging adulthood, and young adulthood, respectively (for more details see Table 1).

**Table 1.** Descriptive statistics for epigenetic aging measures.

| Variable | Early adolescence |  | Late adolescence |  | Emerging adulthood |  | Young adulthood |  |
| --- | --- | --- | --- | --- | --- | --- | --- | --- |
|  | Time 1: | Time 2: | Time 1: | Time 2: | Time 1: | Time 2: | Time 1: | Time 2: |
|  | age 9.5 | age 12 | age 15.5 | age 18 | age 21.5 | age 24 | age 27.5 | age 30 |
| *N* (MZ pairs/DZ pairs) | 326 (82/81) |  | 372 (87/99) |  | 152 (42/34) |  | 126 (52/11) |  |
| Chronological Age *M* (*SD*) | 9.51 (0.33) | 11.83 (0.58) | 15.48 (0.31) | 17.75 (0.58) | 21.58 (0.37) | 23.89 (0.62) | 27.60 (0.70) | 29.92 (1.04) |
| Horvath Accel. *M* (*SD*) | -0.02 (2.64) | -0.25 (3.35) | -0.07 (3.56) | 0.28 (3.99) | 0.47 (4.30) | 0.29 (4.38) | -0.30 (4.17) | -0.52 (4.20) |
| PedBE Accel. *M* (*SD*) | -0.10 (0.91) | -0.21 (0.98) | 0.10 (0.95) | 0.24 (1.14) | 0.16 (1.21) | 0.17 (1.36) | -0.22 (1.24) | -0.39 (1.30) |
| GrimAge Accel. *M* (*SD*) | 0.31 (2.03) | -0.08 (2.41) | -0.44 (2.32) | 0.11 (2.40) | 0.09 (2.23) | 0.01 (2.35) | 0.37 (2.59) | -0.14 (2.61) |
| DunedinPACE *M* (*SD*) | 0.00 (0.07) | -0.01 (0.08) | -0.01 (0.08) | 0.00 (0.08) | 0.00 (0.08) | 0.00 (0.08) | 0.02 (0.08) | 0.02 (0.09) |

*Note.* Four cohorts (early adolescents, late adolescents, emerging adults, and young adults) were measured twice, approximately 2.5 years apart. MZ, monozygotic; DZ, dizygotic; *M*, mean; *SD*, standard deviation; Accel., Acceleration.

### DNA methylation

#### Saliva sampling and DNA extraction

Saliva sampling for DNA extraction was conducted as a part of two TwinLife satellite projects, namely TwinSNPs and TECS. Twins and their family members were invited to participate in the molecular genetic analysis during TwinLife face-to-face interviews carried out in 2018–2020. During face-to-face interviews, interviewers offered to participate in the molecular genetic studies, which involved genotyping and epigenetic profiling. A short video explaining the study goals and procedures of the saliva sampling accompanied the recruitment procedure. The second saliva collection took place in 2021. This time, participants were contacted by mail. In addition to the invitation to participate, respondents received consent forms, saliva collection toolkits, and corresponding instructions. For all participants who provided valid consent, saliva samples were sent to the researchers at the University Hospital Bonn, Germany. The samples were collected using Oragene® saliva self-collection kits (OG-500 and OG-600; DNA Genotek, Canada) and DNA was extracted following the manufacturer’s protocol.

#### DNA methylation assessment

The assessment of DNA methylation profiles was conducted in a set of 2 128 samples, of which the present sample constituted a subset. The samples were randomized by age, sex, zygosity, and complete versus single twin pair across 23 plates (266 arrays) using the R package “Omixer” (version 1.4.0; [68]). The family identifier was used as a blocking variable to ensure that all samples and timepoints of a twin pair ran on the same batch/array. Bisulfite-conversion of 500 ng of DNA was performed using the D5033 EZ-96 DNA Methylation-Lightning Kit (Deep-Well; Zymo Research Corp., Irvine, USA), and methylation profiles were assessed using the Infinium MethylationEPIC BeadChip v1.0 (Illumina, San Diego, CA, USA).

Preprocessing of the methylation samples (*n* = 2 128) was performed following a standard pipeline by Maksimovic et al. [42] using the R package “minfi” (version 1.40.0; [2]). The following sample exclusion criteria were applied: a mean detection *p* value > .05 (*n* = 3) and sex mismatches between estimated sex from methylation data and confirmed phenotypic sex (*n* = 14). To normalize the beta values, we applied stratified quantile normalization^4^ [72], followed by BMIQ [70]. In further analysis, we removed probes containing SNPs (*n* = 24 038), X or Y chromosome probes (*n* = 16 263), cross-hybridizing probes (*n* = 46 867) and polymorphic probes (*n* = 297) according to Chen et al. [11] and McCartney et al. [45], and probes with a detection *p* value > .01 in at least one sample (*n* = 79 922). Afterwards, beta-values were transformed into M-values, and batch-effects were removed using Combat [36]. For this purpose, we examined the strength of associations between three possible batches (plate, slide, and array) and the first five principal components via principal component analysis. The strongest batch effects were iteratively removed. Sample mix-ups were checked with MixupMapper [74]. The comparison of beta values with the existing genotype data (assessed using Global Screening Arrays [GSA+MD-24v3.0-Psych-24v1.1, Illumina, San Diego, CA, USA]; for further details see [13]) revealed four mix-ups. For these four cases, both samples (first and second measurements) were removed (total of *n* = 8 individual samples). The resulting DNA methylation dataset comprises 2 102 samples of *n* = 1 055 participants (among which *n* = 488 complete twin pairs or 976 individuals with two timepoints of DNAm data) and 698 472 probes.

The cell type composition was estimated using the R package “EpiDish” (version 2.10; [71]) and the epidish() function. Because estimated cell type proportions of saliva consisted of three (highly correlated) cell types (leukocytes, epithelial cells, and fibroblasts), we obtained the first principal component (for each epigenetic measurement, explaining 99.6% and 99.7 % of cell type variation in the three estimated cell types at the first and second epigenetic measurements, respectively) to control for cell type composition in the following analyses.

#### Calculating epigenetic age, epigenetic age acceleration, and pace of epigenetic aging

There are several types of epigenetic aging estimators, each based on a unique training model with a variable number of CpGs, source tissues, and age ranges [3, 7]. To capture potential differences in genetic and environmental contributions between different epigenetic aging measures, we selected four commonly used ones (two trained on chronological age, and two trained on biological health indicators) for our analysis: the Horvath clock [27], the PedBE clock [47], GrimAge2 (an improved version of the original GrimAge; [40,41]), and DunedinPACE ([5]; Additional file 2: Table S2 and Figure S1). All four epigenetic aging measures have previously been shown to be associated with aging-related outcomes and to be sensitive to external drivers of aging (e.g., stress; [12, 19, 41, 46, 50, 63]). In particular, GrimAge and DunedinPACE have previously been associated with external drivers of aging in saliva of children and adolescents [17, 58, 60]. The specific background of the training sample of each epigenetic aging estimator is provided in Additional file 2: Table S2.

By the inclusion of the Horvath clock and the PedBE clock, we could address how differences in the training age range and tissue can potentially correspond to the estimates of the genetic and environmental contributions to the variance in epigenetic aging. The Horvath clock was trained as a pan-tissue epigenetic aging estimator in a large age range (from 0 to 101 years; [27]), while the PedBE clock was developed in buccal cells in younger age groups (from 0 to 20 years; [47]). By the inclusion of GrimAge2 and DunedinPACE, we addressed differences in the effects of the training phenotypes: both of these estimators, in contrast to Horvath and PedBE clocks, were trained on biological health indicators. In particular, GrimAge is trained on plasma proteins and smoking [41] and DunedinPACE is trained on biomarkers of organ systems integrity, such as blood pressure or serum leptin levels [5]. Finally, the inclusion of DunedinPACE allowed, in addition, to address the differences in the study design. Namely, the Horvath and PedBE clock, as well as GrimAge are trained on the cross-sectional data, while DunedinPACE accounts for longitudinal patterns of aging.

Epigenetic aging measures were calculated based on the normalized batch-corrected beta values. The Horvath and the PedBE epigenetic age were calculated using the R package “methylclock” (version 1.01; [54]). A total of 334 (94.6%) of the original 353 Horvath clock CpG sites and all 94 CpG sites of the PedBE clock were present in the dataset. Version 2 of GrimAge was computed based on 1 029 CpG sites present in our dataset (out of 1 030 mentioned in the publication) using analysis code from Lu et al. [40] provided by A. Lu and S. Horvath via personal correspondence. The DunedinPACE values were calculated with the DunedinPACE R package (https://github.com/danbelsky/DunedinPACE/) using all of 173 CpG sites.

The four epigenetic age estimators were significantly positively correlated with chronological age (*r*(Horvath epigenetic age) = .80, *p* < .001; *r*(PedBE epigenetic age) = .77, *p* < .001; *r*(GrimAge) = .80, *p* < .001; and *r*(DunedinPACE) = .08, *p* < .001; Additional file 2: Figure S2). For the Horvath epigenetic age, the Median Absolute Error (MAE) estimates ranged from 2.75 to 3.70 (3.26 to 3.65 for early adolescents, from 2.75 to 2.93 for late adolescents, from 3.33 to 3.34 for emerging adults, and from 2.89 to 3.71 for young adults). For the PedBE epigenetic age, the MAE estimates ranged from 1.07 to 12.72 (2.22 to 2.91 for early adolescents, from 1.08 to 2.01 for late adolescents, from 6.49 to 7.51 for emerging adults, and from 11.61 to 12.72 for young adults). For GrimAge, the MAEs ranged from 19.38 to 25.60 (22.96 to 24.38 for early adolescents, from 25.42 to 25.60 for late adolescents, from 21.84 to 22.57 for emerging adults, and from 19.39 to 20.25 for young adults; Additional file 2: Figure S3).

Acceleration estimates for the Horvath, PedBE, and GrimAge epigenetic ages were obtained as follows: epigenetic age was regressed on chronological age (in years and decimal months), biological sex, and cell type composition within each measurement occasion. In addition, the Horvath and PedBE epigenetic ages were regressed on smoking exposure (i.e., beta values of the cg05575921 probe which has previously been shown to predict cigarette consumption in both blood and saliva [55]). This method has been chosen to overcome the missing data on smoking exposure in our sample (especially in the age group of the onset of adolescence), the limited reliability of the self-reports on smoking status and to capture, beyond active smoking, the possible effects of passive smoking exposure. The resulting residuals (i.e., Horvath Acceleration, PedBE Acceleration, and GrimAge Acceleration), defined as *age acceleration* (positive score) or *age deceleration* (negative score) according to the direction of the deviation, were used in our analyses. The DunedinPACE values were controlled for smoking probe, biological sex, and cell type composition. For DunedinPACE, a value of one reflects a pace of aging in line with one year of chronological aging. Correspondingly, values greater than one reflect an increased and values lower than one a decreased pace of aging.

### Analytical strategy

#### Main analyses

To examine the contributions of genetic and environmental factors to the variance in epigenetic aging, we applied three biometrical variance decomposition models, including (1) univariate twin models, (2) bivariate twin models across measurement occasions, and (3) univariate twin models with age as a linear continuous moderator (Additional file 3: Figure S4; [51]). MZ twins share close to 100% of their genetic makeup, while DZ twins share, on average, 50% of genetic variants that differ among humans. As a consequence, the correlation of any genetic factors is approximately 1 for MZ twins, and is assumed to approach .50 for additive genetic factors and .25 for non-additive genetic factors due to allelic dominance deviation for DZ twins. Therefore, a greater correlation within MZ twin pairs than within DZ twin pairs signals the presence of genetic influences on the variance in a specific phenotype, such as epigenetic aging in the current study.

Furthermore, both MZ and DZ twins are assumed to share some prenatal, early life familial, and extra-familial factors [8] that act to increase their resemblance beyond genetic factors. These shared environmental factors are assumed to contribute to the similarity within MZ twins to the same degree as within DZ twins. Only individually-unique environmental factors are assumed to increase the dissimilarity between MZ twin siblings. Under the fulfillment of these assumptions, the classical biometrical model allows for a decomposition of the observed variance into variance attributable to genetic factors (*A* – additive genetic and *D* – non-additive genetic), environmental influences shared by twins (*C*), and environmental influences unique to each twin (*E*, including random error of measurement; [8]). For identification purposes, only three of the variance components can be estimated simultaneously in the classical twin model. Depending on the pattern of the correlations within the MZ and DZ twins, either *D* (if *r*MZ < 2 × *r*DZ) or *C* (if *r*MZ > 2 × *r*DZ) components have to be fixed to zero, resulting in the specification of either ACE or ADE models [16].

As outlined, we started our analysis with univariate twin models to decompose the variance in each epigenetic aging estimator into genetic and environmental components (Additional file 3: Figure S4a). Here, we pooled the data for the twin pairs across the two measurement occasions, resulting in a sample of 976 twin pairs for each univariate twin model. We set the variance of all variance components (*A*, *C*/*D*, and *E*) to 1 for identification purposes and ensured that the statistical conditions specified for twin modeling, namely nonsignificant differences in means and variances across co-twins and zygosity groups, were met in our sample. Then, we decided for each of the four epigenetic aging measures whether to use ACE or ADE models.

After that, among the different possible model variants (ACE, AE, CE, and E; or ADE, AE, and E), we identified the respective model that provided the best fit to the data considering model parsimony based on the chi-square difference test. In cases where alternative models (e.g., AE and CE) were nested in one more complex model (ACE) and did not differ significantly from the latter model based on the chi-square test, the model with a lower Akaike information criterion (AIC) was considered to fit the data better. Among the two nested models (e.g., AE vs. ADE) that differed significantly based on the chi-square test, the model with the higher number of parameters (e.g., ADE) was considered as the model that fits the data best [21]. For the best-fitting models, we calculated the standardized coefficients with 90% confidence intervals.

In the second step, we specified bivariate twin models across the two measurement occasions (488 twin pairs) to examine the genetic and environmental sources of variance in epigenetic aging considering the repeated measurement (Additional file 3: Figure S4b). In this bivariate factor model, we analyzed the first measurement of epigenetic aging as the first variable and the second measurement of epigenetic aging as the second variable. Genetic and environmental factors were specified for each measurement occasion and were allowed to correlate over time in the sense of a *correlated factors model* [39]. For identification purposes, the path coefficients of the latent factors were set to 1. This model specification allowed us to estimate, along with genetic and environmental contributions to the variance in epigenetic aging at the first and second measurement occasions, the genetic and environmental covariances between the two timepoints. First, we ensured that the statistical conditions specified for twin modeling were met in our sample and specified the full model. The full model included three estimates per factor and two estimates of expected means. As was done for the univariate twin models, we have chosen among the ACE, AE, CE, and E models or, alternatively, among the ADE, AE, and E models which of these models provided the best balance between parsimony and fit. The most parsimonious model showed the lowest AIC and did not significantly fit the data worse than a more complex model based on the chi-square difference test. Using the most parsimonious model, unstandardized and standardized variance components and genetic and environmental correlations were estimated. To examine whether or not the genetic and environmental contributions to the variance could be considered constant over time, we then restricted the genetic and environmental variance components to be equal across the first and second measurements of epigenetic aging. The model with equal variances was compared to the less constrained model with unequal variance components over time using the chi-square difference test.

In the third step, we specified twin models with age as a linear continuous moderator of the variance components. Four age cohorts with two measures each were pooled together, resulting in eight unique age groups from 9.5 to 30 (see Table 1) with age gaps of about two and a half years between measurement occasions and age gaps of about three and a half years between the second measure of the younger cohort and the first measure of the subsequent cohort. Here, we tested whether the age of participants moderated genetic and environmental contributions to the variance in epigenetic aging to unveil possible hints on genelenvironment interactions (Additional file 3: Figure S4c; [56]). Since genelenvironment interaction effects cannot be directly assessed in classical twin models, the estimates of genetic and environmental contributions could be confounded with interaction effects [56]. In the case of interactions between additive genetic factors (*A*) and environmental factors that are shared by siblings reared together (*C*), unconsidered interaction effects would be confounded with estimates of additive genetic effects. Potential but unconsidered interaction effects between genetic factors (*A*) and environmental factors not shared by siblings reared together (*E*) would be confounded with estimates of nonshared environmental effects. While the first interaction pattern is more plausible in younger twins, who are still living in a common household, the latter is more plausible in adult years, when twins live in different households and gain more separate life experiences. As a consequence, increasing estimates of genetic contributions to the variance in epigenetic aging are more plausible in adolescence and emerging adulthood than in adulthood when accumulating effects of individually-unique experiences of twins may or may not interact with genetic factors [10, 32].

To investigate the age-related moderation of the genetic and environmental variance components of epigenetic aging, age was added as a continuous linear moderator to the univariate models. This partitioned the ACE/ADE variance components into non-moderated baseline ACE/ADE components, age-moderated ACE/ADE components, a baseline intercept level, and independent linear and quadratic effects of age, producing a model with 9 parameters in total. The model was still identified as the saturated model had in total 10 measured parameters. We started with the full models (with 9 estimated parameters) and then dropped nonsignificant paths to obtain more parsimonious models with more degrees of freedom. We excluded parameters until we reached the model that fitted the data best, taking model parsimony into account (i.e., a reduced model with a lower AIC that did not fit the data significantly worse based on the chi-square test). The variance components, confidence intervals, and standardized estimates were then derived.

#### Sensitivity analyses

In the final step, two sensitivity analyses were performed. As a first sensitivity analysis, we scrutinized the models of the main analysis further not controlling the epigenetic aging measures either for sex, smoking probe, or cell type composition. We repeated the univariate, bivariate, and univariate age moderation analyses, first, for all chosen epigenetic aging measures unadjusted for sex, second, for three epigenetic aging measures unadjusted for smoking, and, finally, for four epigenetic aging measures unadjusted for cell type composition. For the second sensitivity analysis, we rerun the age-moderated univariate models excluding the second timepoint to support the patterns observed in age-moderated univariate twin models using the pooled sample. By doing this, we could check if the trends in the genetic and environmental contributions to the variance in epigenetic aging across age cohorts were similar to what we observed in the main analysis with the pooled data across age cohorts and measurement occasions, ensuring that drawing the same participants twice in the analysis sample, using the first and second timepoints, did not affect the main analysis results.

The analyses were performed in R studio 2023.06.2 with R version 4.2.3. For data preparation, we used the R packages “tidyverse” [75], “ltm” [62], “dlyr” [23], and “psych” [61]. For the variance decomposition, we used “OpenMX” [52].

#### Power analyses

For the estimation of power to detect the model with the best balance between parsimony and fit to the data, we applied the functions proposed by Verhulst [73]. In detail, we applied the function powerValue() along with acePow() function for univariate and bivPow() for bivariate twin models. To address moderation, we applied a function developed for binary moderators, sexLimPower(), and fitted paired age groups. We formulated two possible expectations regarding genetic factors (*A*) based on estimates obtained in larger sample studies. Previous research suggests that the contribution of genetic factors (*A*) to the variance of the Horvath clock is 13% across the lifespan [37] or equals 37% at age 15 (combination with middle-aged mothers; [67]). Our expectation regarding shared environmental factors (*C*) was based on plots for MZ and DZ twins in Li et al. ([37]; page 8, Figure 4, cohabitation-dependent CE model). Here, we identified approximated estimates of *C* for our sample’s mean age (17.1 years) of about 35%. The expectations for *A* and *C* were set to be the same across time. A power calculation for the univariate models without moderation demonstrated that 976 pairs (526 MZ and 450 DZ twin pairs) were sufficient for detecting the *C* component (power .99), while power for detecting the *A* component varied from .40 to .99 depending on the estimates of *A*. In the bivariate case without moderation, 488 pairs (263 MZ and 225 DZ twin pairs) were sufficient (.96 to .99) to detect the *C* component. For the detecting of the *A* component, the power ranged from .27 to .99 depending on the estimates of *A*. Finally, power calculation for models with moderation based on the power calculation for the binary moderation demonstrated that the power to detect the *A* component across the age groups varied in the univariate analysis from .12 to .61. The power to detect *C* varied from .52 to .94.

## Results

### Descriptive analysis

The means and standard deviations for the epigenetic aging measures across age groups are presented in Table 1. The twin correlations of epigenetic aging measures for MZ and DZ twins for each cohort and timepoint separately are given in Table 2 (see also Additional file 3: Table S3 for 95% confidence intervals). Rank-order stability estimates within and across twins’ measurements for MZ and DZ twins are shown in Additional file 3: Table S4.

**Table 2.**
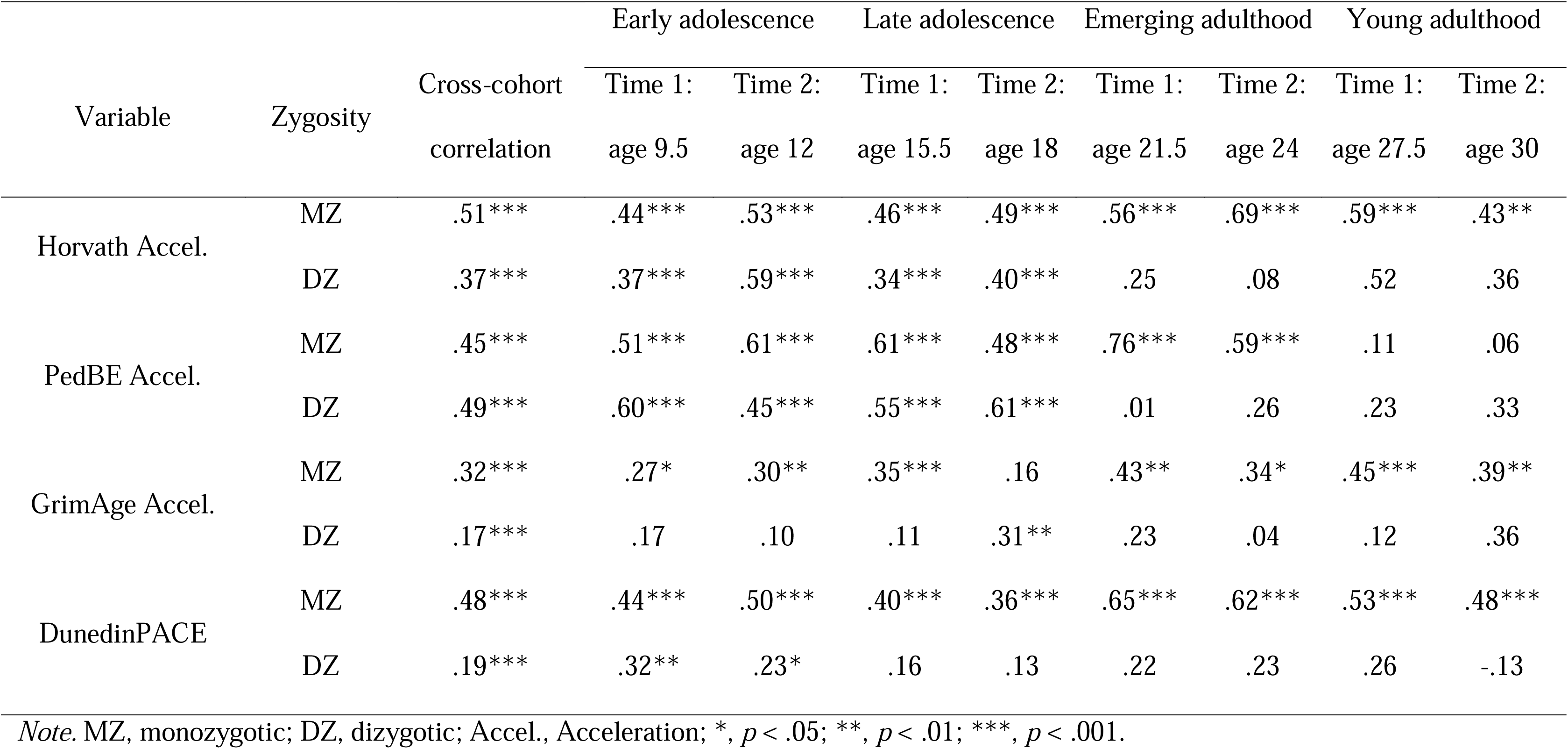
Spearman correlations for monozygotic and dizygotic twins for epigenetic aging measures.

| Variable | Zygosity | Cross-cohort<br>correlation | Early adolescence |  | Late adolescence |  | Emerging adulthood |  | Young adulthood |  |
| --- | --- | --- | --- | --- | --- | --- | --- | --- | --- | --- |
|  |  |  | Time 1: | Time 2: | Time 1: | Time 2: | Time 1: | Time 2: | Time 1: | Time 2: |
|  |  |  | age 9.5 | age 12 | age 15.5 | age 18 | age 21.5 | age 24 | age 27.5 | age 30 |
| Horvath Accel. | MZ | .51*** | .44*** | .53*** | .46*** | .49*** | .56*** | .69*** | .59*** | .43** |
|  | DZ | .37*** | .37*** | .59*** | .34*** | .40*** | .25 | .08 | .52 | .36 |
| PedBE Accel. | MZ | .45*** | .51*** | .61*** | .61*** | .48*** | .76*** | .59*** | .11 | .06 |
|  | DZ | .49*** | .60*** | .45*** | .55*** | .61*** | .01 | .26 | .23 | .33 |
| GrimAge Accel. | MZ | .32*** | .27* | .30** | .35*** | .16 | .43** | .34* | .45*** | .39** |
|  | DZ | .17*** | .17 | .10 | .11 | .31** | .23 | .04 | .12 | .36 |
| DunedinPACE | MZ | .48*** | .44*** | .50*** | .40*** | .36*** | .65*** | .62*** | .53*** | .48*** |
|  | DZ | .19*** | .32** | .23* | .16 | .13 | .22 | .23 | .26 | -.13 |

*Note.* MZ, monozygotic; DZ, dizygotic; Accel., Acceleration; \*, $p < .05$ ; \*\*, $p < .01$ ; \*\*\*, $p < .001$ .

Horvath and PedBE Acceleration, clocks trained on the chronological age, demonstrated age acceleration in the groups of late adolescents and emerging adults and age deceleration in the two other groups of early adolescents and young adults (see Table 1). For GrimAge Acceleration, estimates were higher across several periods, including the onset of adolescence, late adolescence, and young adulthood. The estimates of DunedinPACE showed a faster pace during young adulthood compared to other age periods.

The standard deviations of the four epigenetic aging estimators across age groups indicated patterns of increased variance across time and age stages (Table 1 and Additional file 3: Figure S5). The two measures trained on chronological age, Horvath and PedBE Acceleration, showed a slightly decreasing trend during young adulthood, though. In the eldest age group (*M* = 29.92) compared to the youngest age group (*M* = 9.51), the standard deviation was 60% greater for Horvath Acceleration, 43% greater for PedBE Acceleration, and 29% greater for both GrimAge Acceleration and DunedinPACE.

The twin correlations for Horvath and PedBE Acceleration demonstrated substantial similarity for adolescent twins, with only marginal differences between MZ and DZ twins (Table 2), indicating contributions of shared environmental influences to the variance in epigenetic aging during this period of life. In groups of emerging adults and later, the correlations for MZ twins were greater than the correlations for DZ twins (except the nonsignificant MZ and DZ twin correlations for PedBE Acceleration in young adulthood), indicating genetic influences on the variance in epigenetic aging in adulthood. In contrast, Spearman correlations for GrimAge Acceleration and DunedinPACE were greater for MZ twins than for DZ twins across almost all age groups, indicating genetic contributions to the variance in these epigenetic aging measures.

The rank-order stability across two measurement points was approximately *r* = .50 with no differences between ages and between MZ and DZ twin groups for all measures except for GrimAge Acceleration, where rank-order stability was lower in younger age groups, and PedBE Acceleration, where rank-order stability was greater for DZ twins as compared to MZ twins in young adulthood (Additional file 3: Table S4). Rank-order stability across matched MZ twin and co-twin measurements tended to be greater than those of DZ twins in adulthood, indicating declining contributions of shared environmental factors and increasing genetic contributions to rank-order stability with age.

### Univariate twin model analyses of epigenetic aging

To perform univariate twin modeling, we first examined whether the statistical conditions specified for twin modeling were met (Additional file 3: Table S5). We observed small differences in means across co-twins for Horvath Acceleration and DunedinPACE, but the models with the expected means and variances constrained to be equal across twin siblings’ order and zygosity did not fit the data significantly worse.

To make a choice regarding the ACE or ADE model, we examined if the correlations for MZ twins were more than two times higher than the DZ twin correlations. For Horvath, PedBE, and GrimAge Acceleration, the cross-cohort MZ twin correlations were not more than twice as strong as the DZ twin correlations, indicating the absence of non-additive genetic factors (*D*) and the presence of additive genetic factors (*A*) and shared environmental factors (*C*) instead: *r*MZ = .51 and *r*DZ = .37 regarding Horvath Acceleration, *r*MZ= .45 and *r*DZ = .49 regarding PedBE Acceleration, *r*MZ = .32 and *r*DZ = .17 regarding GrimAge Acceleration (see Table 2). The cross-cohort correlations for MZ twins for DunedinPACE were more than two times higher than correlations for DZ twins indicating, besides additive genetic factors (*A*), the presence of non-additive genetic factors (*D*): *r*MZ = .48 and *r*DZ = .19.

The results of the univariate twin model analyses of the epigenetic aging measures are presented in the Additional file 3: Tables S6-S7. Cross-cohort standardized genetic and environmental variance components for the full models are presented in Additional file 3: Figure S6 and the best-fitting models in Figure 2. For Horvath Acceleration, we observed a better fit of the full model (ACE), indicating that the variance in Horvath Acceleration was best explained by additive genetic as well as environmental factors shared and not shared by twins. PedBE Acceleration, GrimAge Acceleration, and DunedinPACE estimates in the univariate analysis were better represented by restricted models: a CE model for PedBE Acceleration and an AE model for both GrimAge Acceleration and DunedinPACE. The variance in PedBE Acceleration was thus best explained by environmental factors shared and not shared by twins, whereas the variance in GrimAge Acceleration and DunedinPACE was best explained by additive genetic and individually-unique environmental factors.

**Figure 2.**
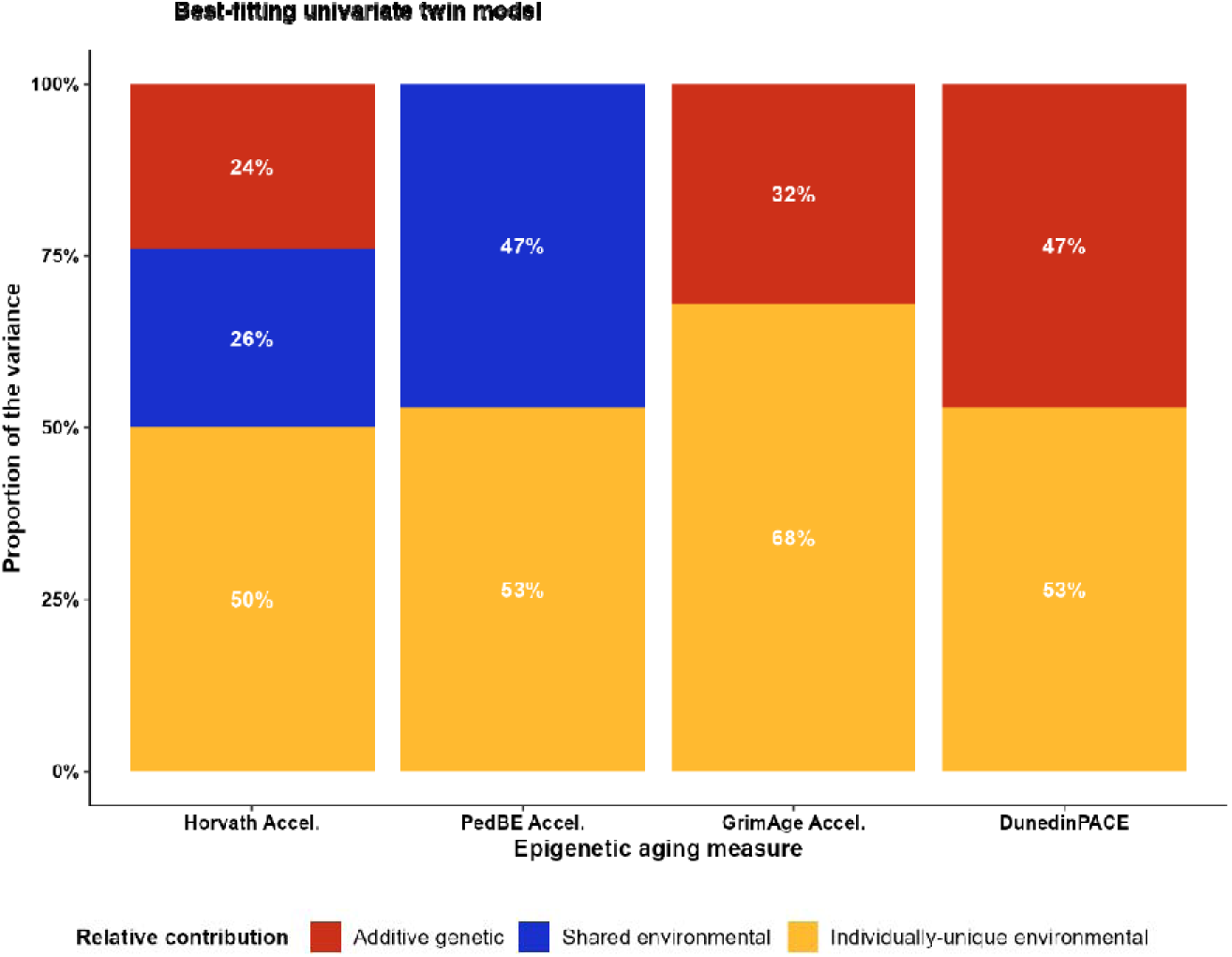
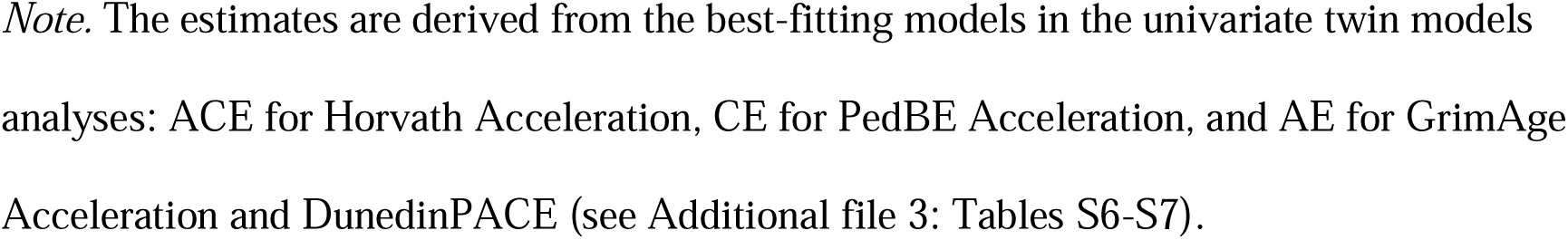
Cross-cohort standardized genetic and environmental variance components of epigenetic aging measures in the best-fitting model.

### Bivariate twin model analyses of epigenetic aging

In a further step, we decomposed the variance in the two repeated measures of epigenetic aging with the use of bivariate twin models. The models with the expected means and variances to be equal across co-twins and zygosity within each measurement occasion did not fit the data significantly worse than the saturated models (Additional file 3: Table S5). Thus, the results confirmed that the statistical conditions specified for bivariate twin modeling were met.

Following the previously reported MZ and DZ twin correlations (Table 2), we similarly specified ACE models for Horvath, PedBE, and GrimAge Acceleration, and an ADE model for DunedinPACE. As for the univariate analyses, a bivariate ACE model for Horvath, a CE model for PedBE, and an AE model for GrimAge and DunedinPACE provided the best balance between parsimony and fit (see Additional file 3: Tables S8-S9). Figure 3 presents the results from the variance decompositions of both timepoints based on the best-fitting reduced models. The results from the full models can be found in Additional file 3: Figure S7. The best-fitting reduced models were used as a basis to test whether genetic and environmental variance components could be constrained to be equal over time.

**Figure 3.**
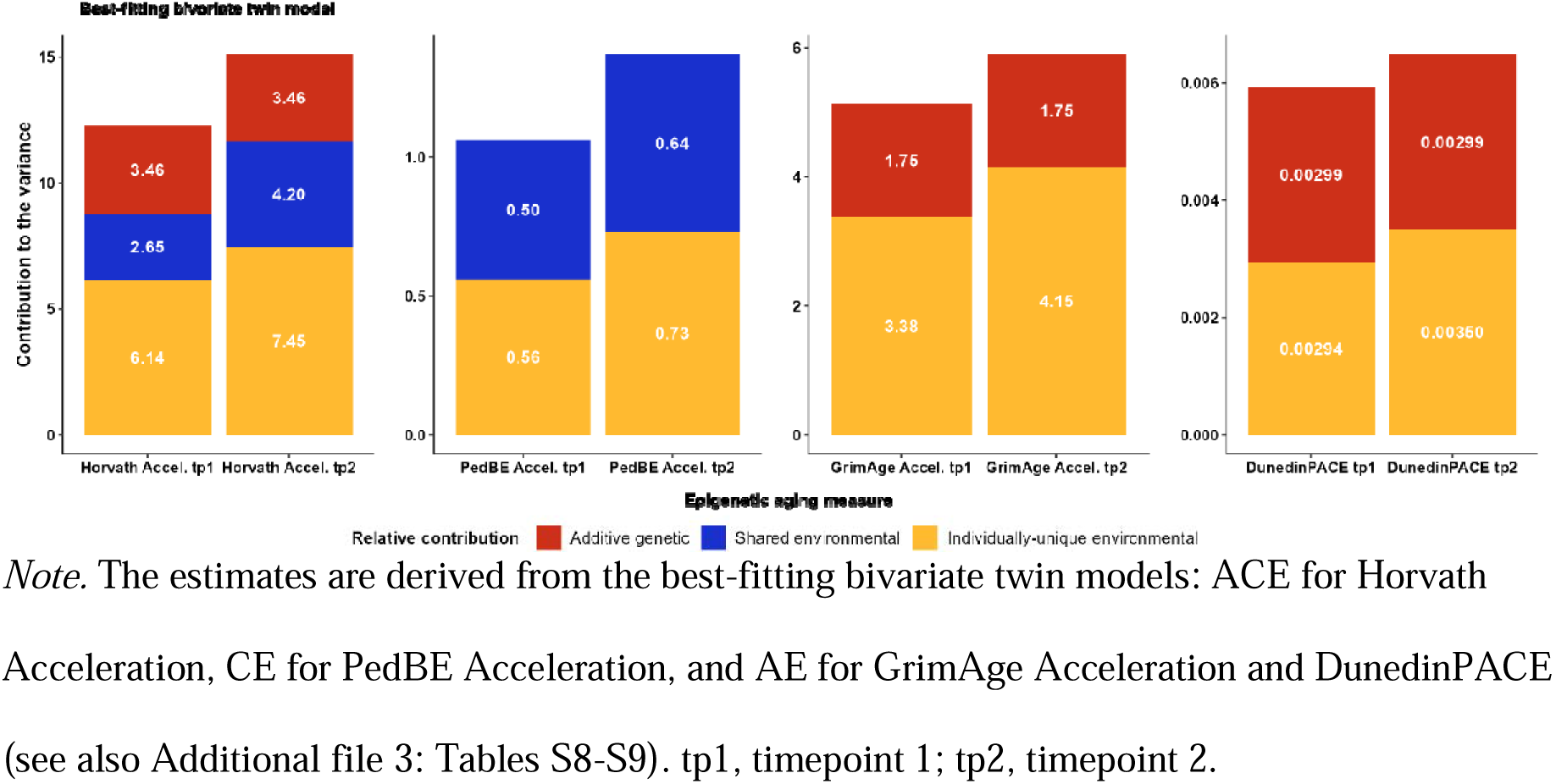
Unstandardized genetic and environmental variance components of epigenetic aging measures over 2.5 years.

**Figure 4.**
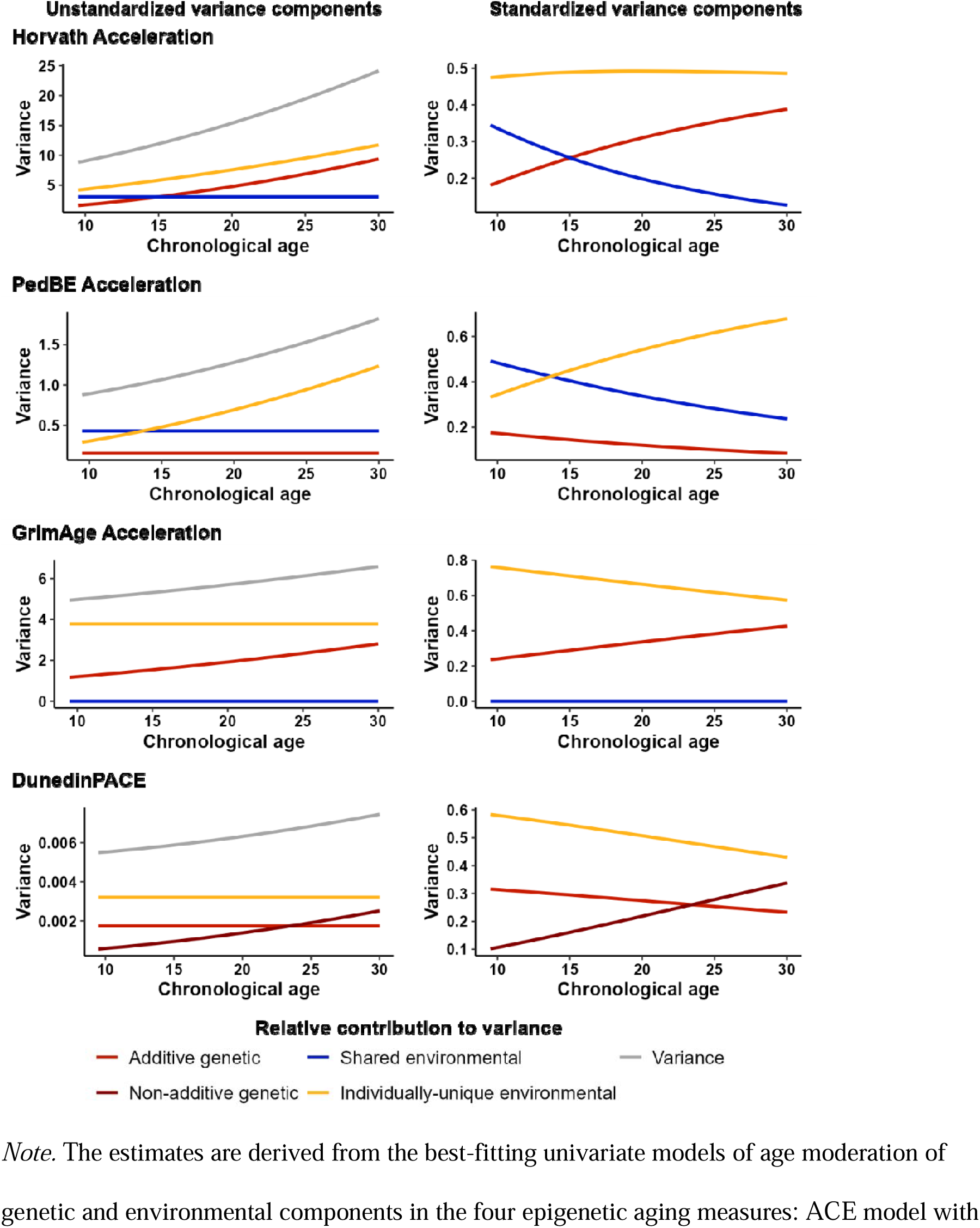
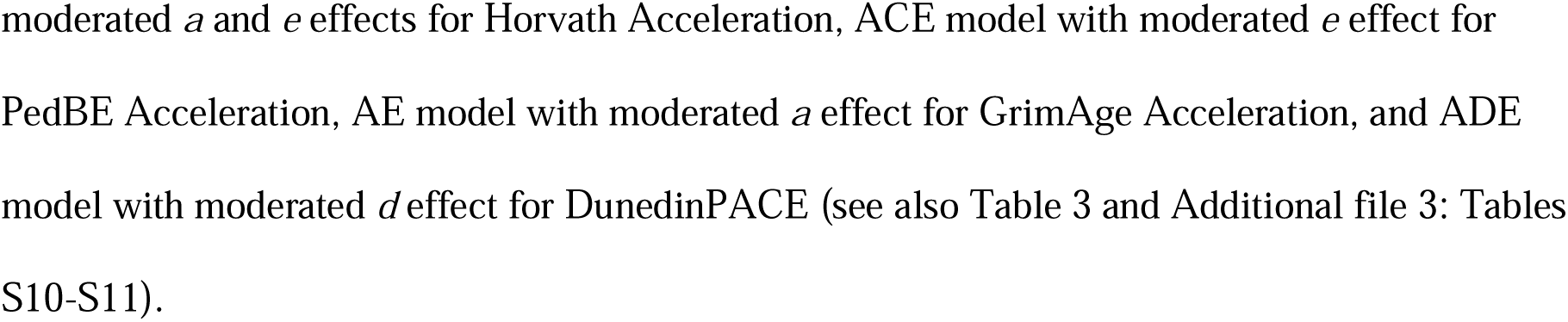
Unstandardized and standardized variance components derived from age-moderated models of epigenetic aging measures.

For each of the four epigenetic aging measures, restricting the variance components of all three or two genetic and environmental factors (*A*, *C*, and *E*) to be equal across the two timepoints, respectively, resulted in a significantly worse fit to the data (see Additional file 3: Table S8). Accordingly, our results suggested that the relative contributions of the genetic and/or environmental factors changed over time. However, setting only the variance component due to additive genetic factors to be equal across the two timepoints did not result in a significantly worse fit to the data for the epigenetic aging measures trained in adult samples (i.e., Horvath, GrimAge, DunedinPACE). These results suggested almost equal absolute contributions of additive genetic factors over time. In contrast, variance components due to environmental factors that are not shared by twins increased from the first to the second measurement timepoint for all epigenetic aging measures, resulting in increasing variance over time (see Figure 3). In addition, for epigenetic aging measures trained on the chronological age, the variance components due to shared environmental factors seemed to increase over time. This pattern was not observed for the epigenetic aging measures trained on health biomarkers, for which shared environmental factors were negligible.

Then, we estimated the degree to which genetic and environmental factors correlated over time. The three epigenetic aging measures trained in adult samples demonstrated significant positive genetic correlations from the first to the second measurement timepoint. The highest positive correlation of additive genetic factors was observed for Horvath Acceleration (*r_A_* = .90, *p* < .001), followed by GrimAge Acceleration (*r_A_* = .82, *p* < .001) and DunedinPACE (*r_A_* = .74, *p* < .001). Epigenetic aging estimators trained to predict chronological age, in turn, demonstrated statistically significant positive correlations of the shared environmental factors over time (*r_C_* = .92, *p* < .001 for Horvath Acceleration and *r_C_* = .87, *p* < .001 for PedBE Acceleration). Unique environmental factors correlated significantly for PedBE Acceleration (*r_E_* = .22, *p* < .001), DunedinPACE (*r_E_* = .18, *p* < .001), and Horvath Acceleration (*r_E_* = .15, *p* < .001), but not significantly for GrimAge Acceleration (*r_E_* = .06, *p* = .19). Generally, the correlations of individually-unique environmental factors were weak (≤ .22), while the correlations of additive genetic factors and shared environmental factors were substantial (≥ .74). In line with the a priori power analysis, the power to distinguish between the ACE/ADE, CE, and AE models varied from .41 to 1 for Horvath Acceleration, from .40 to 1 for PedBE Acceleration, from .39 to .93 for GrimAge Acceleration, and from .41 to .76 for DunedinPACE. The power to distinguish the best-fitting model from the E model was 1 for all epigenetic aging measures (Additional file 3: Table S8).

### Age moderation of genetic and environmental differences in epigenetic aging

In the final step, we explored whether the contributions of the *A*, *C* (or *D*), and *E* components to the variance of epigenetic aging differed across age groups (Table 3, Figure 4, Additional file 3: Tables S10-S11, Figure S8). Here, we only present the parameter estimates of the models that fitted the data best. The model fit statistics can be found in Additional file 3: Table S10.

**Table 3.**
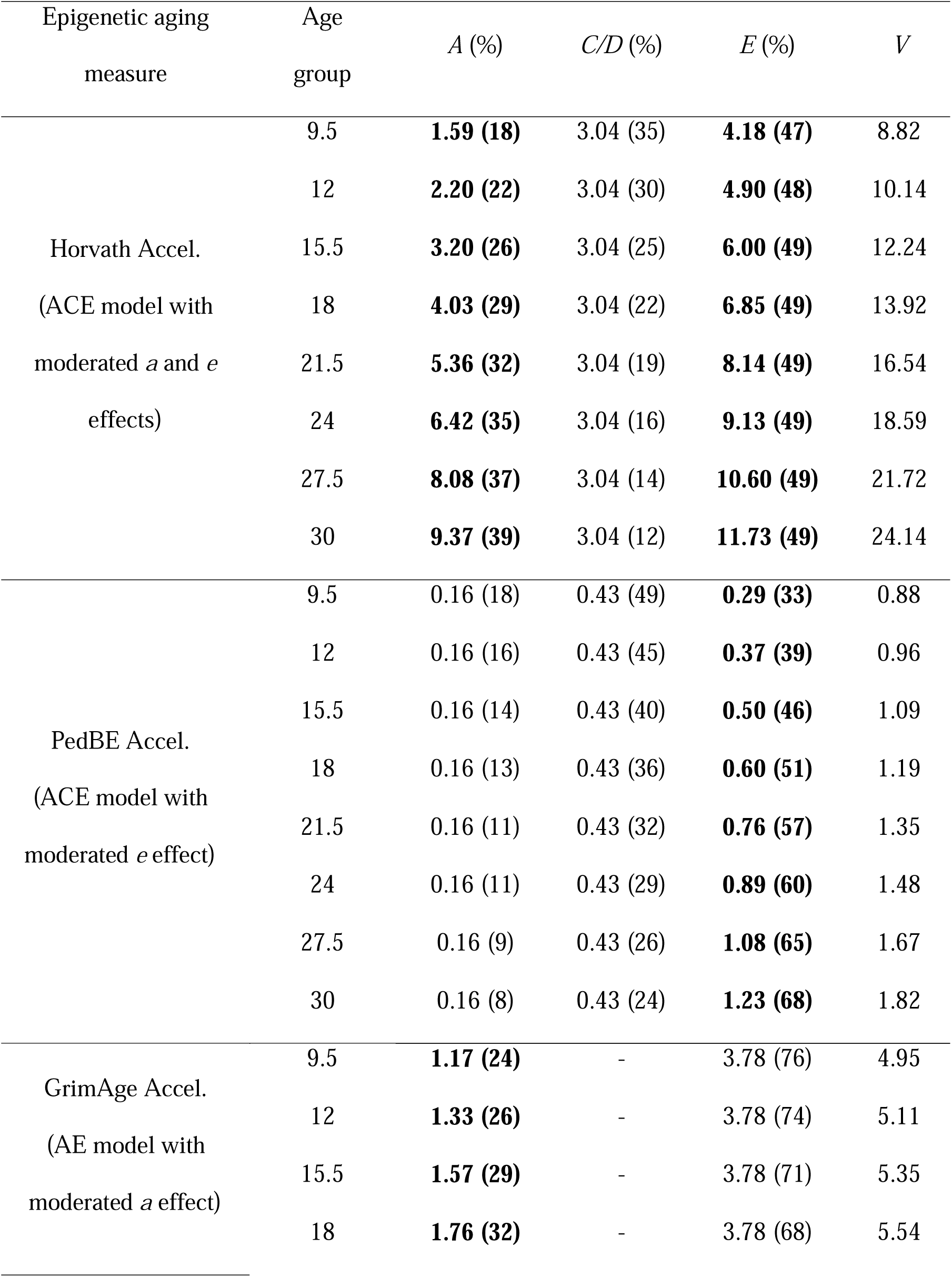
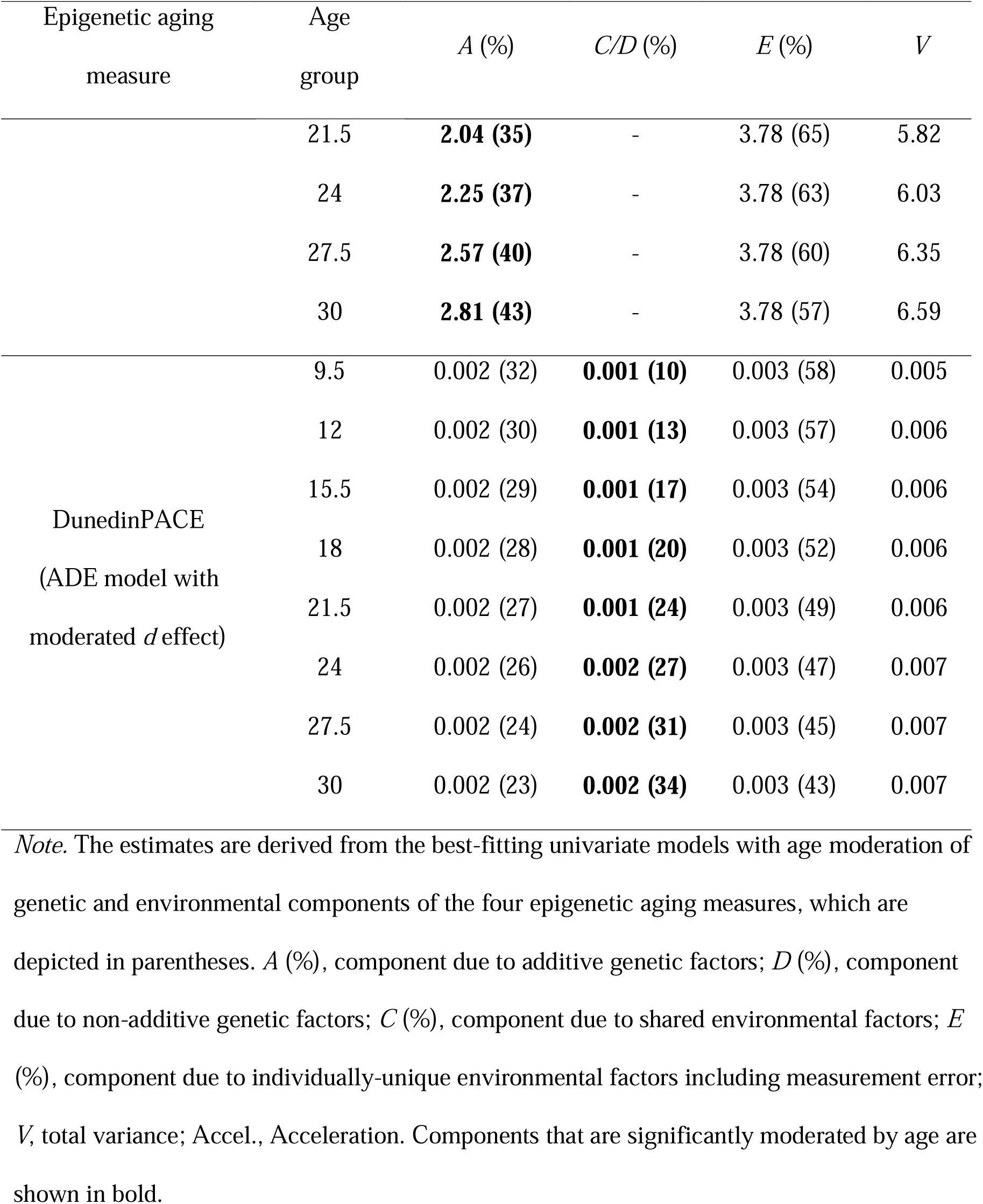
Variance components derived from age-moderated univariate models.

When moderation by age in the univariate models was allowed, statistically significant moderation was observed for all epigenetic aging measures resulting in generally increasing variance across age groups for all epigenetic aging measures. However, different factors turned out to be relevant for this increase in different epigenetic aging measures. For acceleration measures trained on chronological age, including Horvath and PedBE Acceleration, we observed the moderation of the individually-unique environmental component (*E*). That is, the unique environmental variance increased across age groups regarding both estimators. The contribution of the unique environmental factors to the variance in the Horvath and PedBE Acceleration was larger for the oldest age group (young adults) compared to the youngest ones (4.18 at the onset of adolescence versus 11.73 in young adults for Horvath Acceleration; 0.29 at the onset of adolescence versus 1.23 in young adults for PedBE Acceleration, see Table 3). Moderation of the variance attributable to genetic factors, with higher contribution in young adulthood compared to early adolescence, was found for Horvath and GrimAge Acceleration (1.59 versus 9.37 for Horvath Acceleration and 1.17 versus 2.81 for GrimAge Acceleration) as well as for DunedinPACE (non-additive genetic component was 0.00056 in younger adolescents and 0.00251 in young adults). Thus, the narrow-sense heritability potentially increased across age groups from 18% to 39% for Horvath Acceleration and from 24% to 43% for GrimAge Acceleration. The broad-sense heritability (*A + D*) of DunedinPACE increased from 42% to 57%.

In contrast to the bivariate model and the univariate twin model without age moderation, the CE model for PedBE (with moderated *e* effect) showed a higher AIC value (5 550.13) in comparison to the best-fitting model in the age-moderated univariate analysis (moderated *e* effect in ACE model; AIC = 5 548.89). However, the power to distinguish between these two models was limited (.30) and both models did not fit significantly worse to the data in comparison to the corresponding more complex models. Likewise, the AE model for DunedinPACE yielded a significantly worse fit to the data compared to the full model (–2LL = –4 562.38, *df* = 1 947, AIC = –4 552.38, *p* = .048) in the univariate analysis with age moderation.

### Sensitivity analysis

In the first part of the sensitivity analysis, we performed three separate analyses for univariate twin modeling, bivariate twin modeling, and univariate twin modeling with age as a moderator in epigenetic aging measures unadjusted either for biological sex, the smoking probe, or cell type composition (Additional file 4: Tables S12-S29). Not controlling for sex or smoking did not lead to different conclusions compared to those from the main analysis. Not controlling for cell type composition obscured some of the findings from the main analysis (Additional file 4: Tables S24-S29). For Horvath Acceleration, the AE model demonstrated a better fit in the univariate analysis without moderation, while the ACE model still yielded a better fit in the bivariate analysis and the analysis with age moderation. For PedBE Acceleration, we observed, along with the moderation of the *E* component, the moderation of the *A* component in the better fitting ACE age-moderated model. GrimAge Acceleration demonstrated a better fit of the CE models across the univariate and bivariate model in the sensitivity as opposed to the main analysis with the latter indicating a better fit of the AE models. Not controlling for cell type composition in DunedinPACE demonstrated, finally, a better fit of the ADE model in the univariate and bivariate twin models, as well as the moderation of the *E* instead of the *D* component in the age-moderated univariate twin model. The analysis of epigenetic aging measures unadjusted for cell type composition did not reach sufficient power to distinguish between models. The observed pattern is potentially explained by the fact that different tissues have heterogeneous cell type compositions [69], which, in turn, can have strong independent associations with age [76]. Therefore, the measures of epigenetic aging unadjusted for cell type composition might reflect, in addition to the genuine age-associated signal of DNAm, heterogeneity in different cell type compositions regarding DNAm as well as a link between cell type composition and age.

As the second part of the sensitivity analysis, we specified and fitted the univariate models of the variance decomposition with age moderation for the first epigenetic measurement to explore if drawing the data from two measurements to increase statistical power could affect the results. The models’ goodness-of-fit statistics are presented in the Additional file 4: Table S30. The same models were observed as the best-fitting to the data for Horvath Acceleration, GrimAge Acceleration, and DunedinPACE. Similar to the main analysis, we observed that the models with the moderated *a* and *e* paths for Horvath Acceleration, with the moderated *a* path for GrimAge Acceleration, and with the moderated *d* path for DunedinPACE showed a better fit to the data. For PedBE Acceleration, we did observe the moderation of the *e* path as also reported in the main analysis. In contrast to the main analysis, however, the model without *A* factors demonstrated a better fit to the data. The observed difference in results probably could originate from the halved sample size and decreased power to detect lower contributions of additive genetic factors to the variance.

## Discussion

In this study, we investigated the genetic and environmental contributions to the variance in epigenetic aging from early adolescence to young adulthood. Three main results can be summarized. First, genetic and environmental contributions to the variance in epigenetic aging varied between differently developed epigenetic aging estimators. Second, stable differences in epigenetic aging across two measurement occasions, 2.5 years apart, were mainly attributable to genetic and shared environmental factors, while individually-unique environmental factors represented the primary source of instability (i.e., rank-order change) over time. Third, the variance in epigenetic aging increased from early adolescence to young adulthood (i.e., from age 9.5 to 30 years), which was due to increasing genetic or environmental contributions to the variance, depending on the specific estimator for epigenetic aging.

### Genetic and environmental contributions to the variances of differently developed epigenetic aging estimators

The performed univariate and bivariate twin model analyses demonstrated both similarities and differences in the genetic and environmental contributions across the four differently developed epigenetic aging measures. One of the similarities was that at least half of the differences in every epigenetic aging estimator was attributable to individually-unique environmental factors not shared by twins reared together. The contributions of these factors to the variance could emphasize the role of individually-unique life experiences contributing to individual differences in epigenetic aging. Childhood and adolescence are a sequence of sensitive and critical developmental periods, during which the effects of positive and negative experiences can be particularly influential. Experiences of violence in children or institutional caring [29], for example, have already been mentioned to predict greater epigenetic age, affecting ongoing brain and emotional regulation development [65]. Individually-unique environmental factors can also reflect the *epigenetic drift*, stochastic changes in DNAm. When epigenetic drift occurs in epigenetic aging estimators’ locus-specific DNAm, the individually-unique environmental variance estimated in biometrical variance decomposition models would partly reflect epigenetic drift [20]. Epigenetic drift with age could also explain the greater power of epigenetic age to predict chronological age at the onset of life compared to childhood and adolescence [47, 67].

Genetic and shared environmental factors, in turn, contributed differently to the four epigenetic aging measures. In particular, the variance in the best-fitting univariate models was explained either by genetic (GrimAge Acceleration and DunedinPACE), by shared environmental (PedBE Acceleration), or by both genetic and shared environmental factors (Horvath Acceleration). The developmental design of the investigated epigenetic aging estimators suggests that these differences are not random. The larger contribution of the shared environmental factors to the variance in Horvath Acceleration and PedBE Acceleration corresponds to the training outcome of these estimators. Both are trained to predict chronological age, which is by definition shared by twins of one pair. The transition to adulthood, in turn, could explain why genetic contributions were observed for all epigenetic aging measures except for PedBE Acceleration. In contrast to the PedBE clock, which was initially developed in a sample of individuals up to 20 years of age, the Horvath clock, GrimAge, and DunedinPACE were trained in adults (see Additional file 2: Table S2). The genetic contributions we observed for the latter three epigenetic aging estimators, but not for PedBE Acceleration, might potentially characterize intrinsic epigenetic processes that are activated later in life and shared to a greater extent by genetically identical MZ twins compared to DZ twins. These processes can include the processes of physical development, for example the transition from puberty to the maintenance of biological systems and the high organism functionality. As these processes are less pronounced during adolescence, they may be not fully reflected in the genetic contributions to the variance in PedBE Acceleration.

The patterns observed for the genetic and environmental contributions to the variance in epigenetic aging were also reflected in stable differences across measurements, as shown in the bivariate twin analyses. Here, substantial positive genetic correlations over time were found in three estimators trained in adults. In contrast to environmental differences, the absolute size of genetic variance in epigenetic aging estimators remained constant over time, suggesting that genetic factors represent a primary source of stable differences in the epigenetic aging rate, at least across 2.5 years in our study. Estimators trained to predict chronological age (Horvath Acceleration and PedBE Acceleration) also demonstrated substantial positive correlations of shared environmental factors. However, their contributions to the variance increased from the first to the second measurement occasion, respectively. This indicates that the similarity of same-aged siblings and differences between families in epigenetic aging relative to chronological age tend to increase over time. While genetic and shared environmental factors represent the primary source of stability, individually-unique environmental factors were found to be only marginally stable over the period of 2.5 years. Moreover, individual differences in all epigenetic aging estimators increased over time due to an increase in those environmental influences not shared by twins. This indicates that individually-unique environmental sources represent the primary source of inter-individual differences in intra-individual change in epigenetic aging over time.

Finally, the training design could potentially explain the specific variance patterns for DunedinPACE, for which we observed a smaller variance in general and its modest increase. This is in line with previous findings suggesting a remarkable stability of DunedinPoAm, the predecessor of DunedinPACE, for the age period from 7 to 20 [53]. Beyond that, DunedinPACE was characterized by the greatest contribution of genetic factors to the variance and was the only epigenetic aging estimator for which the presence of non-additive genetic factors was indicated to some extent. These findings could also signal the differences between cross-sectional and longitudinal training designs. While the GrimAge, Horvath, and PedBE clocks were developed by analyzing cross-sectional samples, DunedinPACE was trained on longitudinal data, accounting for the within-individual trends of aging. Representing the speed of epigenetic aging based on 12 months, DunedinPACE does not directly relate to chronological age and the accumulation of aging processes, which are commonly represented by measures of epigenetic age acceleration. The smaller changes in the variance in DunedinPACE, along with the largest contribution of the genetic factors, suggest that their role in the pace of epigenetic aging across adolescence and young adulthood is potentially more stable than in epigenetic acceleration measures.

### Genetic and environmental contributions to age-associated differences in epigenetic aging estimators

The analysis of the age-associated differences demonstrated that the genetic and environmental contributions to the variance were not constant across age groups. These findings can be discussed in the context of potential gene[environment correlations and gene[environment interactions relevant for variance in epigenetic aging from early adolescence to young adulthood. That is, in addition to the patterns observed in the univariate and bivariate models, the three estimators Horvath Acceleration, GrimAge Acceleration, and DunedinPACE, trained in adults, demonstrated an increased variance across age groups due to genetic (additive or non-additive) factors. First, this potential increase is consistent with longitudinal studies on the SNP-based heritability of epigenetic aging and the heritability of other complex phenotypes. Specifically, in the sample of children and their parents, the SNP-based heritability was demonstrated to be 0% during the onset of life but higher later in life [67]. Similarly, genetic contributions to the variance in complex phenotypes, such as intelligence, depressive symptoms, and personality traits, demonstrate increasing heritability from childhood to adulthood [6, 32, 49].

The observed phenomenon of the increased role of genetic factors during this age is commonly explained by active and evocative gene[environment correlation in the presence of gene[environment interaction, “[…] when people become more and more autonomous and self-directed during adolescence. They have increasing opportunities to pave their own way, evoke, select and create environments that match their heritable tendencies. These environments in turn can provide experiences that have the potential to reinforce the pre-existing tendencies” ([32]; p. 238). One illustrative example of an active gene[environment correlation is the self-selection of individuals in environments that fit their genetic predispositions [6, 31]. For instance, when individuals with genetic predispositions for a slower epigenetic aging rather develop healthy habits and search for health-promoting environments that fit this genetic tendency, this would reflect processes of active gene[environment transactions resulting in non-random gene– environment correlation. As the increasing importance of active gene[environment correlation would come along with an increase of genetic differences, while a declining role of passive gene[environment correlation (i.e., genetic and environmental factors on offspring’s phenotypes are correlated due to parental phenotypes and behavior) would be reflected by decreasing effects of shared environmental influences [10, 56], our findings can be considered in line with this interpretation.

Since we did not detect increasing genetic variance over an interval of 2.5 years in our sample, which is not entirely compatible with the interpretation of gene–environment correlation, alternative explanations must also be considered. Specifically, the gene[environment interplay underlying estimates of genetic variance can also reflect interactions of genetic and environmental factors shared by twins, which would also account for the finding of non-perfect correlations of genetic factors over the 2.5-year period in our sample [32, 56]. This form of gene–environment interplay might be more relevant for twins reared together and, thus, more likely to occur during adolescence, when twins still share a common family household. Previous studies illuminating the role of shared environmental factors emphasized the driving role of the prenatal environment [38] and cohabitation for differences in epigenetic aging, implying that the longer relatives live together, the more similar they are epigenetically [37]. From adolescence to adulthood, the role of these factors seems to be steadily replaced by genetic and individually-unique environmental factors as well as by interactions between them. As cohabiting is more common before adulthood, our findings extend previous conclusions, implying an increasing role of active gene[environment correlation in the presence of interactions between genetic and environmental factors, when relatives, on average, transit to adulthood and tend to start living separately. These processes are reflected in the trajectories of gene[environment interactions and the relationships between individuals and their immediate environments. This would include, for instance, the transition from the sorting into fitting environments to the individual adjustment to changing environments, or finding a balance between person[environment fitting and adjustment [31].

Furthermore, the contributions of individually-unique environmental influences to the variance notably increased during adolescence and emerging adulthood in Horvath and PedBE Acceleration, epigenetic aging estimators trained on chronological age. In addition to the discussed accumulated effects of unique life experiences and epigenetic drift, the observed pattern can signal increasing measurement error in epigenetic aging measures, which was particularly evident for PedBE Acceleration. Being trained in a younger population [47], its application in adults was less precise. In our analysis, PedBE Acceleration had a larger error in predicting chronological age in older cohorts, while its MZ and DZ twin correlation estimates varied largely. Measurement error, however, is likely not the sole driver for the increasing variance in epigenetic aging due to individually-unique environmental factors, as the same trend of increasing environmental variance was observed for Horvath Acceleration. This estimator was also trained to predict chronological age and its mean prediction error was consistent across analyzed age stages and groups.

Although we observed that the individually-unique environmental factors increase from the first to the second timepoint in GrimAge Acceleration and DunedinPACE, the contributions of individually-unique environmental factors to variance in both estimators were not found to be moderated by age in the pooled univariate analysis. In contrast to epigenetic aging estimators trained on chronological age, the two latter measures were developed in (exclusively adult) blood samples and rely on intrinsic biological functioning. In particular, GrimAge is linked to the level of plasma proteins [41] and DunedinPACE reflects, among other factors, the level of glycated hemoglobin, leptin, and serum C-reactive proteins [5]. Because these two estimators are also intrinsically driven, these processes may be more elusive and less sensitive to trends in individually-unique environmental contributions when they are measured in saliva.

### The heritability of epigenetic aging based on saliva samples

The age-specific patterns of genetic contributions to the variance in epigenetic aging in saliva obtained in our analysis were rather consistent with the assessments reported in blood. However, our heritability estimates were, on average, lower than those reported in studies based on blood tissue samples. For example, previous research reported that genetic differences accounted for 69%, 73%, and 68% of the variance in Horvath Acceleration, GrimAge Acceleration, and DunedinPACE in 22-year-olds, respectively [34]. The broad-sense heritability for the 21.5-year-old twins in our analysis was substantially lower (32%, 35%, and 51% respectively). The comparison of the heritability of PedBE Acceleration in saliva with its heritability in other sources of DNAm (e.g., buccal cells) was not possible, because no study so far, to our knowledge, provided heritability estimates for PedBE Acceleration and our study indicated decreasing or zero heritability. The observed differences in heritability estimates across sources of DNAm for epigenetic aging measures could be related to variations in the age compositions of the studies’ samples, the presence of age moderation, or the genuine differences in epigenetic aging across tissues [77]. Therefore, further studies are needed to investigate whether the observed pattern of the lower heritability of epigenetic aging based on saliva can be replicated.

### Limitations and future research

Among the strengths of our study are the relatively large sample size of twin pairs in the field of epigenetics, the focus on both epigenetic age acceleration and the pace of epigenetic aging measures, the examination of genetic and environmental contributions to the covariance in epigenetic aging measures over two timepoints, and, finally, the consideration of age differences during the second and third decades of life. Nevertheless, our study is not without limitations.

First, differences in the performance of epigenetic aging measures across different ages and tissues may have introduced additional sources of measurement error. The Horvath clock, for example, is less precise in younger samples, while the PedBE clock could be more biased in adults [47]. Similarly, our study exclusively utilized saliva samples, but not all applied epigenetic aging estimators were trained on saliva-based DNA methylation profiles. While the Horvath clock is a pantissue epigenetic clock and the PedBE clock was developed in buccal cells, DunedinPACE and GrimAge were trained in blood. Although the pace of aging can be assessed based on saliva [58, 59] and the second version of GrimAge demonstrated good performance in saliva-based DNA methylation data [40], applying these epigenetic aging estimators in a tissue different from the initial design may affect the estimates. The Horvath clock, at the same time, being a pantissue clock, could lose some precision when applied to any tissue.

Second, despite the advantages of a comparatively large sample size with epigenetic data and the use of two measurement points in the biometrical variance decomposition, the post hoc power analysis demonstrated that the differentiation between several nested models was limited, as confidence intervals overlapped. Previous literature has suggested a minimum sample size of 600 twin pairs for univariate analysis and larger samples for detecting possible non-additive genetic factors, which, unfortunately, could not be reached [44, 64]. Nevertheless, we had enough power to detect the presence of genetic or environmental age moderation in epigenetic aging measures.

Third, the utilization of measurements from two timepoints in four birth cohorts could introduce additional sources of variance. Different age groups, for instance, might introduce cohort effects that cannot be entirely disentangled from age effects. In addition, the second measurement was realized during the COVID-19 pandemic. While prior studies did not confirm specific effects of the COVID-19 pandemic on epigenetic aging in adolescents [14], we observed an increased contribution of the individually-unique environmental factors to variance in epigenetic aging at the second measurement. It thus might be that pandemic-related factors, such as changes in health status, lockdowns, and restrictions, formed potential new influential environmental sources that could act very individually. Hence, the role of these factors in epigenetic aging dynamics should be clarified further.

Fourth, classical twin analysis is an efficient way to estimate genetic and environmental sources of variance but at the cost of several assumptions. One assumption is that environmental experiences shared by twins contribute to the similarity within MZ twin pairs to the same degree as within DZ twin pairs. One study proposed that this assumption might not hold true for epigenetic aging, as MZ and DZ pairs can possibly share their preadult environmental effects to different extents [37]. Such phenomenon can have both an environmental or genetic origin [18]. In addition, the classical twin design does not allow, so far, the estimation of shared environmental (*C*) and non-additive genetic (*D*) contributions in one model, while both sources of variance can be relevant for variance in epigenetic aging at the same or during different developmental stages. These aspects should be addressed in further investigations.

## Conclusion

Altogether, the present study demonstrated that the genetic and environmental sources of variance in epigenetic aging differ across different epigenetic aging estimators. For estimators trained in adult samples, larger contributions of genetic factors to the variance in epigenetic aging were observed. Shared environmental factors accounted for variance in epigenetic aging measures trained on chronological age only. Genetic factors and environmental factors shared by twins reared together represented the primary sources contributing to the stability of epigenetic aging differences in corresponding estimators over time, while individually-unique environmental factors represented the primary source of rank-order change.

The increase of the variance in epigenetic aging measures from early adolescence to young adulthood was associated with differences in the contributions of genetic and environmental factors. The variance of epigenetic aging estimators trained in adult samples was characterized by increasing contributions of genetic factors across age. Increasing differences in epigenetic aging measures trained on chronological age, in addition, were driven by increasing contributions of individually-unique environmental factors. These findings can be interpreted in terms of an interplay between genetic and environmental factors and contribute to a better understanding of the epigenetic aging processes during the transition to adulthood.

## Declarations

### Ethics approval and consent to participate

The TwinLife study was reviewed and approved by the German Psychological Society (Deutsche Gesellschaft für Psychologie; protocol number: RR 11.2009). The TwinSNPs project received ethical approval from the Ethical Committee of Bielefeld University (No. 2020-180) and the TwinSNPs and TECS projects received ethical approval from the Ethical Committee of the Medical Faculty of the University of Bonn (Lfd. Nr. 113/18). All participants provided written informed consent. Participation was voluntary and participants could withhold participation at any timepoint. The personal data collection and analysis complied with the corresponding data protection regulations. The statistical analysis was performed on pseudonymized data from the servers of Bielefeld University.

## Availability of data and materials

The TwinLife data are archived in the GESIS data catalog: https://search.gesis.org/research_data/ZA6701. The data are released for academic research and teaching after the data depositor’s written authorization. The epigenetic data are available for collaborators via TwinLife data management.

## Authors’ contributions

DVK, CK, and BM were involved in the study conception and design. DVK analyzed the data and interpreted the results. DVK, YL, AMS, and CK drafted the manuscript. DVK, YL, AMS, DC, JI, CKLP, MMN, EBB, MD, AJF, CK, and BM provided valuable feedback for the manuscript revision. MD & MMN were responsible for obtaining ethical approval. MD, BM, MMN, AJF, CKLP, FMS, EBB, DC, and CK collected the data. AMS, DC, DVK, and CKLP assessed DNA methylation and generated the epigenetic aging scores. BM supervised the study. All the authors have read and approved the final manuscript.

## Supporting information

Previously published estimates of heritability

Performance of epigenetic aging measures

Performance of epigenetic aging measures

Sensitivity analysis

Tables within the manuscript

## Acknowledgments

We thank M. Ruks, A. Kottwitz, and K. Krell from TwinLife data management team for their support; S. Zare, A. Vogelsang, and N. Fricker for their contribution to the DNA extraction and epigenetic profiling; M. Ruks and L. Weigel for valuable advice on the manuscript.

## Additional files

**Additional file 1.** The SNP-, pedigree-, and twin-based heritability of epigenetic aging estimated in previous studies.

**Additional file 2.** Performance and training characteristics of epigenetic aging measures. The figures and tables represent CpG sites overlap between the epigenetic aging measures, Median Absolute Errors and mean deviations for three epigenetic age estimators.

**Additional file 3.** Supplementary Tables and Figures.

**Additional file 4.** Sensitivity analysis results. Tables S12-29 report goodness-of-fit statistics and standardized variance components for univariate, bivariate, and univariate age-moderated twin models of epigenetic aging measures unadjusted for biological sex, smoking probe, or cell type composition. Table S30 reports the univariate age-moderated twin models of epigenetic aging measures for the first epigenetic measurement.

## Footnotes

1 The proportion of the variance explained by all single nucleotide polymorphisms (SNPs) used in a genome-wide association study in conventionally unrelated individuals [78].

2 The proportion of the variance explained by the patterns of genetic resemblance among not-too-distantly related family members [24].

3 The proportion of the variance explained by the patterns of genetic resemblance between monozygotic and dizygotic twins.

4 The chemical chip used to measure DNA methylation signal from extracted DNA consists of two types of probes (I and II). These two types of probes produce different distributions of signals that can introduce a technical noise when analyzed together. Stratified quantile normalization fixes this by stratifying the probes into type I and II, then using type I signals as ‘anchors’ to normalize Infinium II signals at the level of probe coverage categories [72].

