## Supplementary material for "Genetic and Environmental Contributions to Epigenetic Aging Across Adolescence and Young Adulthood": Previously published estimates of heritability

**Additional file 1: Previously published estimates of heritability of the analyzed epigenetic aging measures**

**Table S1**

*Estimates of heritability of epigenetic aging measures from twin-, SNP-, and pedigree-based studies*

| Study | *M* (*SD*/Range) | Heritability | Twin pairs | Individuals | Measure | Method |
| --- | --- | --- | --- | --- | --- | --- |
| Horvath, 2013 [2] | 0 (0-0) | 1.00 | - | 53 | Horvath Accel. | Twin |
| Horvath, 2013 [2] | 63^1^ (45-75) | 0.39 | - | 93 | Horvath Accel. | Twin |
| Levine et al., 2015 [6] | 89 (6.4) | 0.41^5^ | - | 700 | Horvath Accel. | SNP |
| Marioni et al., 2015 [8] | 14^2^ (9-23)  47^3^ (33-75) | 0.43 | 178 | 614 | Horvath Diff. | Pedigree (twins, siblings, parents) |
| Lu et al., 2016 [7] | 48 (16-96) | 0.69/ 0.01/0.01/0.59^6^ | - | 112 | Horvath Accel. | SNP |
| Lu et al., 2016 [7] | 52 (1-102) | 0.15/0.14^7^ | - | 201 | Horvath Accel. | SNP |
| Simpkin et al., 2016 [11] | 15^4^ (15-NA) | 0.37 | - | 2 036 | Horvath Accel. | SNP (mothers, children) |
| Jylhävä et al., 2019 [3] | 70 (8.4) | 0.55 | 104 | 208 | Horvath Accel. | Twin |
| Jylhävä et al., 2019 [3] | 79 (8.3) | 0.51 | 104 | 208 | Horvath Accel. | Twin |
| Sillanpää et al., 2019 [10] | 23 (0.9) | 0.74 | 289 | 578 | Horvath Accel. | Twin |
| Sillanpää et al., 2019 [10] | 61 (3.7) | 0.53 | 164 | 328 | Horvath Accel. | Twin |
| Kankaanpää et al., 2021 [4] | 22 (0.7) | 0.69 | 285 | 570 | Horvath Accel. | Twin |
| Kankaanpää et al., 2021 [4] | 62 (4.1) | 0.61 | 235 | 470 | Horvath Accel. | Twin |
| Kankaanpää et al., 2021 [4] | 22 (0.7) | 0.62 | 285 | 570 | GrimAge Accel. | Twin |
| Kankaanpää et al., 2021 [4] | 62 (4.1) | 0.58 | 235 | 470 | GrimAge Accel. | Twin |
| Jylhävä et al., 2019 [3] | 70 (8.4) | 0.55 | 104 | 208 | Horvath Accel. | Twin |
| Jylhävä et al., 2019 [3] | 79 (8.3) | 0.51 | 104 | 208 | Horvath Accel. | Twin |
| Kankaanpää et al., 2022 [5] | 22 (0.7) | 0.73 | 365 | 730 | GrimAge Accel. | Twin |
| Kankaanpää et al., 2022 [5] | 22 (0.7) | 0.62 | 365 | 730 | DunedinPoAm | Twin |
| Kankaanpää et al., 2022 [5] | 22 (0.7) | 0.68 | 365 | 730 | DunedinPACE | Twin |
| Hong et al., 2024^8^ [1] | 31 (31-31) | 0.66 |  | 986 | DunedinPACE | Twin |
| Hong et al., 2024 [1] | 70 (70-70) | 0.44 |  | 986 | DunedinPACE | Twin |
| Hong et al., 2024 [1] | 31 (31-31) | 0.76 |  | 986 | PC Horvath Accel. | Twin |
| Hong et al., 2024 [1] | 70 (70-70) | 0.49 |  | 986 | PC Horvath Accel. | Twin |
| Hong et al., 2024 [1] | 31 (31-31) | 0.60 |  | 986 | PC GrimAge Accel. | Twin |
| Hong et al., 2024 [1] | 70 (70-70) | 0.55 |  | 986 | PC GrimAge Accel. | Twin |
| Miao et al., 2024 [9] | 50.24 (26-77) | 0.69 | 134 | **268** | DunedinPACE | Twin |
| Miao et al., 2024 [9] | 54.87 (31-82) | 0.72 | 134 | **268** | DunedinPACE | Twin |
| Miao et al., 2024 [9] | 50.24 (26-77) | 0.70 | 134 | **268** | GrimAge Accel. | Twin |
| Miao et al., 2024 [9] | 54.87 (31-82) | 0.69 | 134 | **268** | GrimAge Accel. | Twin |

*Note.* SNP, single nucleotide polymorphisms-based heritability analysis; Pedigree, pedigree-based heritability analysis; Twin, patterns of genetic resemblance between monozygotic and dizygotic twins; *M*, mean age; *SD*, standard deviation; NA, not available; Accel., the residuals from a linear regression model of epigenetic age on chronological age; Diff., the difference between epigenetic age and chronological age, PC, principal components from the CpG sites derived from the correspondent epigenetic aging measure.

^1^ Median of age.

^2^ Mean age of children.

^3^ Mean age of parents.

^4^ Oldest age combination (15-year-old children and middle age mothers).

^5^ Estimates of heritability for cerebellum/frontal cortex/pons/temporal cortex.

^6^ Estimates of heritability for cerebellum/frontal cortex.

^7^ Estimates of heritability for dorsolateral prefrontal cortex.

^8^ Estimates for the youngest and for the oldest age groups. The heritability estimates for epigenetic aging measures were derived in local structural equation modeling. The total sample size for ages between 30 and 70 is 986, with a raw sample size of 16 at age 30 and 8 at age 70.
