## Supplementary material for "Genetic and Environmental Contributions to Epigenetic Aging Across Adolescence and Young Adulthood": Performance of epigenetic aging measures

**Additional file 2: Performance of epigenetic aging measures**

**Table S2**

| Epigenetic aging measure | Reference | Training Sample | | | TECS | |
| --- | --- | --- | --- | --- | --- | --- |
|  |  | Tissue | Age range (years) | Phenotype | Available CpGs/original epigenetic aging measure CpGs | Correlation with chronological age |
| Horvath clock | Horvath [2] | Multiple incl. blood and saliva | 1–101 | Chronological age | 334/353 | .80 |
| PedBE clock | McEwen et al. [4] | Buccal | 0-20 | Chronological age | 94/94 | .77 |
| GrimAge2 | Lu et al. [3] | Blood | 40–92 | 12 biomarkers of aging (10 of GrimAge version 1 plus log C-reactive protein and log haemoglobin A1C) | 1 030/1 030 | .80 |
| DunedinPACE | Belsky et al. [1] | Blood | 26, 32, 38, and 45 | Longitudinal change in 19 biomarkers of aging (e.g., blood pressure, leptin levels) | 173/173 | .08 |

*Specific backgrounds of training sample of the epigenetic aging estimators*

*Note.* TECS, TwinLife Epigenetics Change Satellite project.

**Figure S1**

*CpG sites overlap between the epigenetic aging measures*

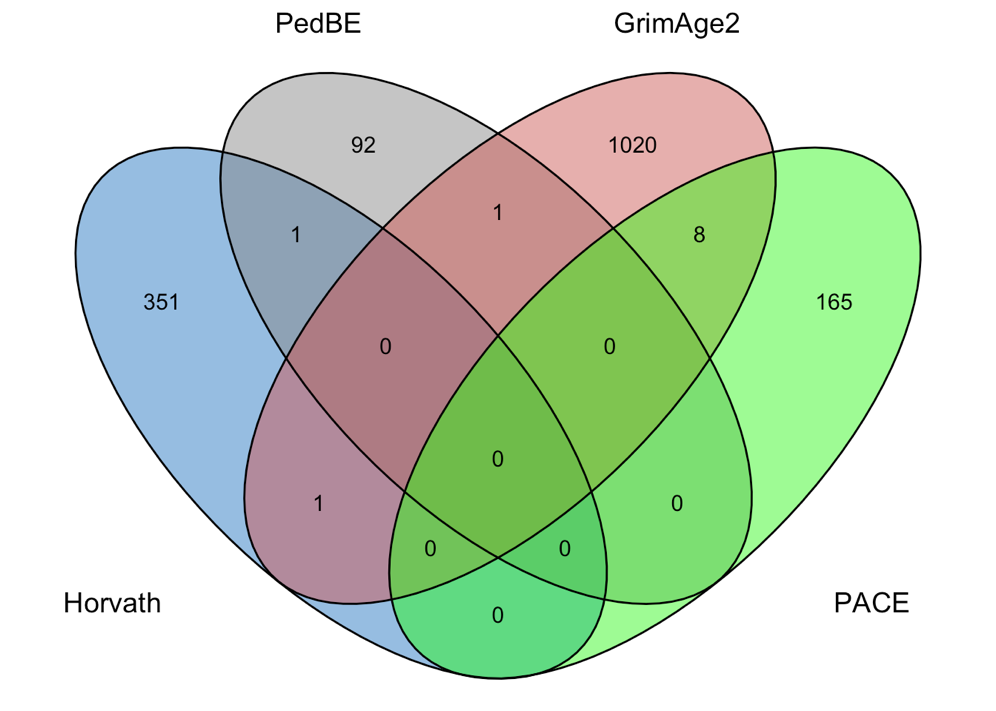

B

A

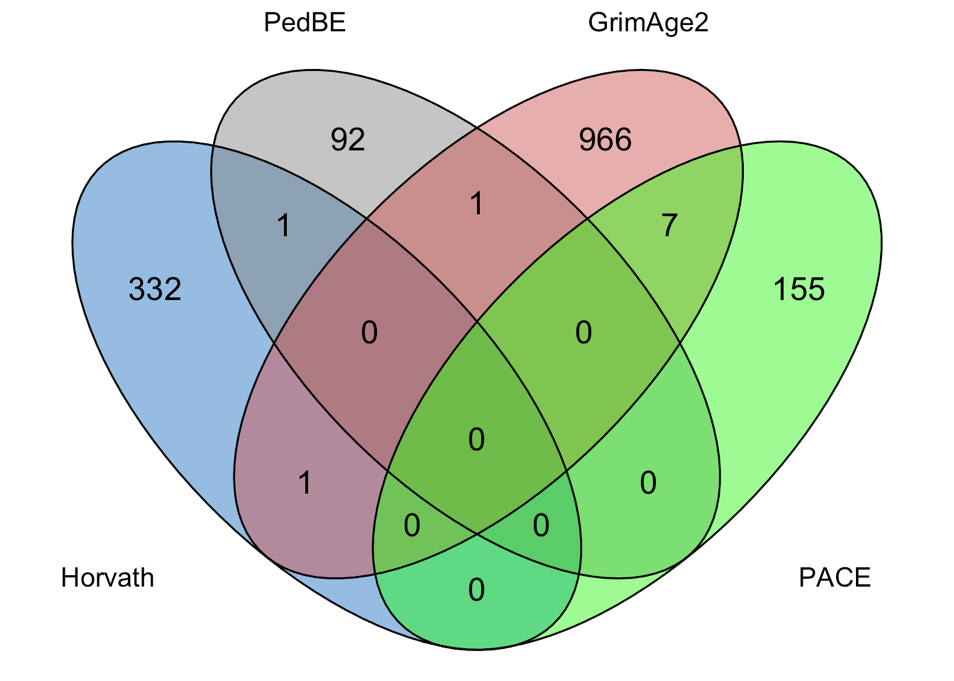

*Note*. Venn diagram of the number of CpG sites overlapping between the Horvath clock, the PedBE clock, GrimAge2 and DunedinPACE estimate using (A) all original epigenetic aging measures CpGs and (B) only CpGs available in the TECS DNA methylation dataset.

**Figure S2**

*Mean deviations for three epigenetic age measures: Horvath epigenetic age, PedBE epigenetic age, and Grimage2*

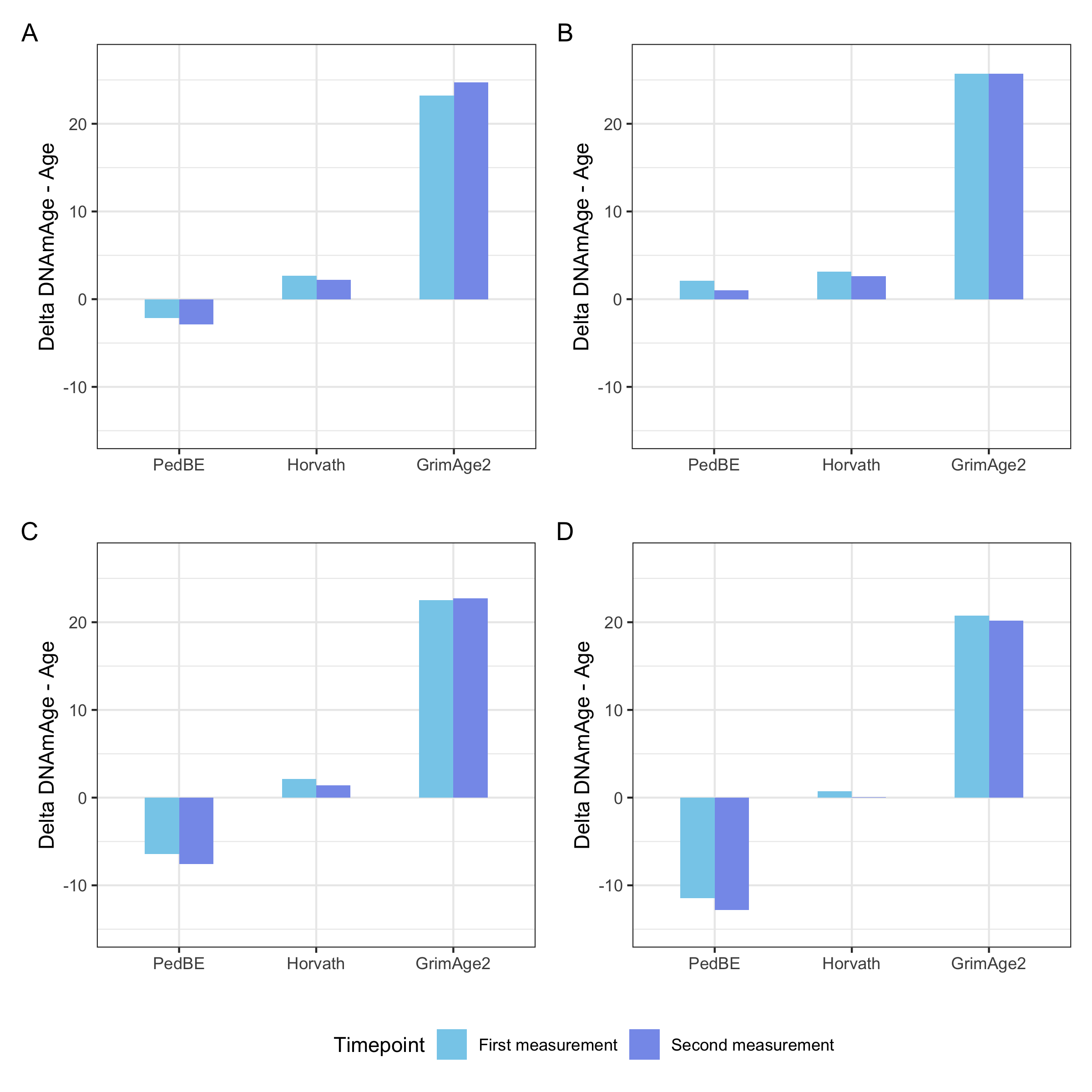
*Note*. Mean deviation computed as the mean differences between chronological and epigenetic age for the first (blue) and second measurement (purple) in early (A) and late adolescence (B), emerging adulthood (C), and young adulthood (D). The closer to zero, the more accurate the corresponding epigenetic estimate.

**Figure S3**

*Median Absolute Errors for three epigenetic age measures: Horvath epigenetic age, PedBE epigenetic age, and Grimage2*

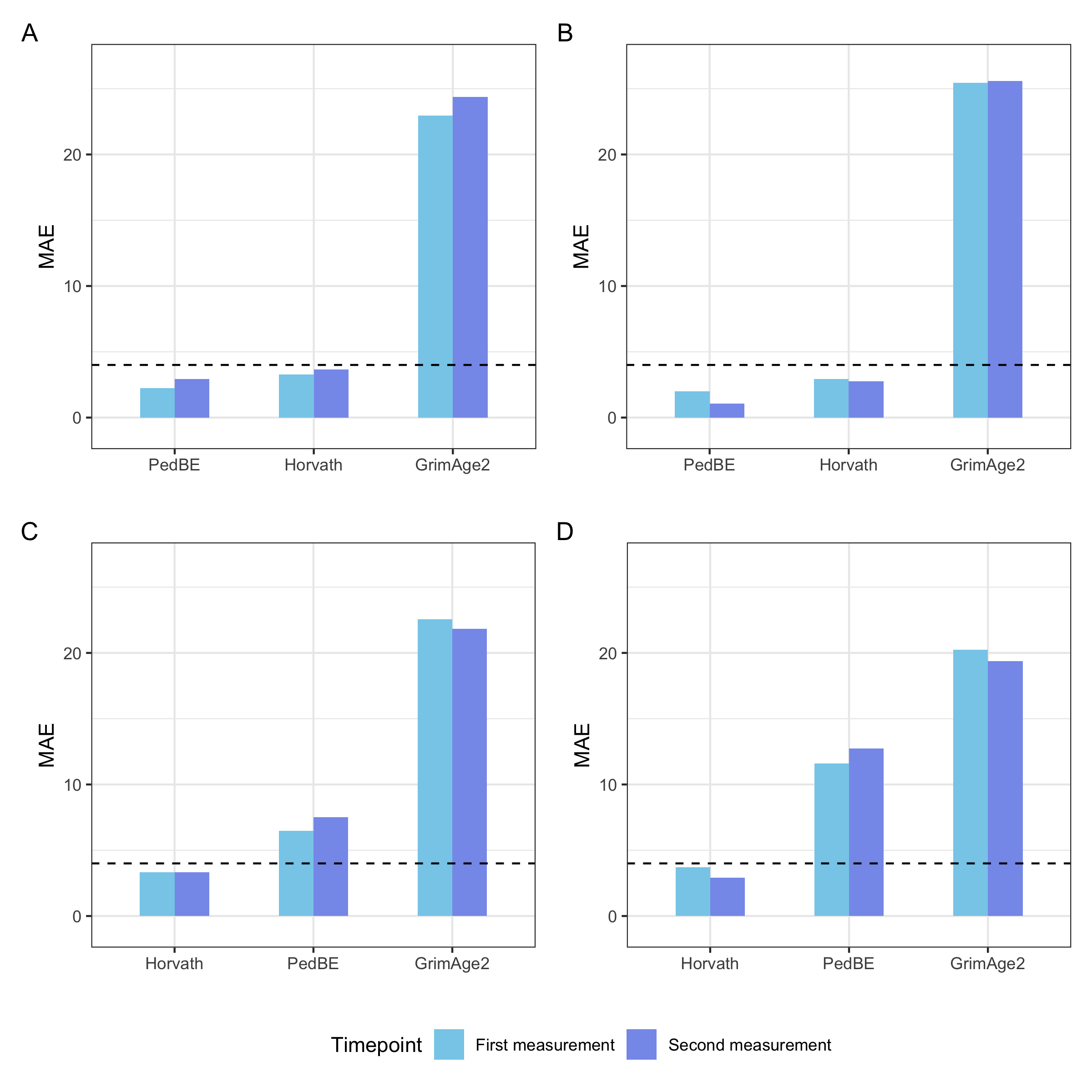

*Note.* Median absolute error computed as the absolute median differences between chronological and epigenetic age for the first (blue) and second measurement (purple) in early (A) and late adolescence (B), emerging adulthood (C), and young adulthood (D). The black dotted line represents MAE = 4. MAE, median absolute error.
