## Supplementary material for "Genetic and Environmental Contributions to Epigenetic Aging Across Adolescence and Young Adulthood": Performance of epigenetic aging measures

**Additional file 3: Supplementary Tables and Figures**

**Table S3**

*Spearman correlations and their 95% confidence intervals for monozygotic and dizygotic twins’ epigenetic aging measures*

|  |  | Early adolescence | | | Late adolescence | | | |
| --- | --- | --- | --- | --- | --- | --- | --- | --- |
| Variable | Zygosity | Time 1:  age 9.5 | Time 2:  age 12 | | Time 1:  age 15.5 | | Time 2:  age 18 | |
| Horvath Accel. | MZ | .44*** [.25,.60] | .53*** [.36,.67] | | .46*** [.28,.61] | | .49*** [.31,.64] | |
|  | DZ | .37*** [.16,.54] | .59*** [.43,.72] | | .34*** [.16,.51] | | .40*** [.22,.55] | |
| PedBE Accel. | MZ | .51*** [.33,.65] | .61*** [.45,.73] | | .61*** [.46,.73] | | .48*** [.30,.63] | |
|  | DZ | .60*** [.43,.72] | .45*** [.26,.61] | | .55*** [.39,.67] | | .61*** [.46,.72] | |
| GrimAge Accel. | MZ | .27* [.05,.46] | .30** [.09,.49] | | .35*** [.15,.52] | | .16 [-.05,.36] | |
|  | DZ | .17 [-.05,.38] | .10 [-.13,.31] | | .11 [-.09,.30] | | .31** [.12,.48] | |
| DunedinPACE | MZ | .44*** [.25,.60] | .50*** [.32,.65] | | .40*** [.20,.56] | | .36*** [.17,.53] | |
|  | DZ | .32** [.11,.51] | .23* [.01,.43] | | .16 [-.04,.35] | | .13 [-.07,.32] | |
| *Table continues on the next page* | | | | | | | | |
|  |  | Emerging adulthood | | | | Young adulthood | | |
| Variable | Zygosity | Time 1:  age 21.5 | | Time 2:  age 24 | | Time 1:  age 27.5 | | Time 2:  age 30 |
| Horvath Accel. | MZ | .56*** [.31,.74] | | .69*** [.49,.82] | | .59*** [.37,.74] | | .43** [.18,.63] |
|  | DZ | .25 [-.10,.54] | | .08 [-.26,.41] | | .52 [-.12,.85] | | .36 [-.30,.79] |
| PedBE Accel. | MZ | .76*** [.60,.87] | | .59*** [.35,.76] | | .11 [-.17,.37] | | .06 [-.21,.33] |
|  | DZ | .01 [-.33,.35] | | .26 [-.09,.55] | | .23 [-.43,.73] | | .33 [-.33,.78] |
| GrimAge Accel. | MZ | .43** [.15,.65] | | .34* [.04,.58] | | .45*** [.20,.64] | | .39** [.13,.60] |
|  | DZ | .23 [-.12,.52] | | .04 [-.30,.38] | | .12 [-.51,.67] | | .36 [-.30,.79] |
| DunedinPACE | MZ | .65*** [.44,.80] | | .62*** [.39,.78] | | .53*** [.30,.70] | | .48*** [.24,.67] |
|  | DZ | .22 [-.13,.52] | | .23 [-.12,.53] | | .26 [-.40,.74] | | -.13 [-.68,.51] |

*Note.* MZ, monozygotic; DZ, dizygotic; Accel., Acceleration; *, *p* < .05; **, *p* < .01; ***, *p* < .001.

**Table S4**

*The rank-order stability estimates of epigenetic aging measures across the two measurement occasions*

| Epigenetic aging measures |  | Early adolescence | | Late adolescence | | Emerging adulthood | | Young adulthood | |
| --- | --- | --- | --- | --- | --- | --- | --- | --- | --- |
|  |  | Time 1:  age 9.5 | Time 2:  age 12 | Time 1:  age 15.5 | Time 2:  age 18 | Time 1:  age 21.5 | Time 2:  age 24 | Time 1:  age 27.5 | Time 2:  age 30 |
| Rank-order stability within twins | | | | | | | | | |
| Horvath Accel. | MZ | .52*** [.40, .63] | | .50*** [.38, .60] | | .53*** [.36, .67] | | .55*** [.40, .67] | |
|  | DZ | .48*** [.35, .59] | | .54*** [.43, .63] | | .56*** [.37, .70] | | .59** [.23, .81] | |
| PedBE Accel. | MZ | .53*** [.41, .63] | | .44*** [.31, .55] | | .54*** [.37, .68] | | .31** [.13, .48] | |
|  | DZ | .52*** [.40, .63] | | .59*** [.50, .68] | | .55*** [.36, .70] | | .82*** [.61, .92] | |
| GrimAge Accel. | MZ | .17* [.02, .32] | | .25** [.10, .38] | | .51*** [.33, .65] | | .52*** [.36, .65] | |
|  | DZ | .14 [-.02, .28] | | .36*** [.23, .47] | | .44*** [.22, .61] | | .26 [-.17, .62] | |
| DunedinPACE | MZ | .39*** [.25, .51] | | .32*** [.18, .45] | | .55*** [.37, .68] | | .53*** [.38, .66] | |
|  | DZ | .43*** [.30, .55] | | .52*** [.41, .62] | | .50*** [.29, .66] | | .45* [.03, .73] | |
| *Table continues on the next page* | | | | | | | | | |
| Epigenetic aging measures \| \| Early adolescence \| \| Late adolescence \| \| Emerging adulthood \| \| Young adulthood \| \|; \| \| Time 1: age 9.5 \| Time 2: age 12 \| Time 1: age 15.5 \| Time 2: age 18 \| Time 1: age 21.5 \| Time 2: age 24 \| Time 1: age 27.5 \| Time 2: age 30 \| | | | | | | | | | |
| Rank-order stability between twins and co-twins | | | | | | | | | |
| Horvath Accel. | MZ | .39*** [.25, .52] | | .45*** [.33, .56] | | .51*** [.33, .65] | | .47*** [.31, .61] | |
|  | DZ | .38*** [.25, .52] | | .37*** [.24, .48] | | .10 [-.14, .33] | | .46* [.05, .74] | |
| PedBE Accel. | MZ | .44*** [.30, .55] | | .48*** [.35, .58] | | .53*** [.36, .67] | | .12 [-.08, .31] | |
|  | DZ | .36*** [.22, .49] | | .51*** [.40, .61] | | .22 [-.01, .44] | | .26 [-.18, .62] | |
| GrimAge Accel. | MZ | .28*** [.13, .42] | | .19* [.04, .33] | | .38*** [.18, .55] | | .34*** [.16, .50] | |
|  | DZ | .17* [.02, .32] | | .08 [-.06, .22] | | .19 [-.06, .41] | | .02 [-.40, .44] | |
| DunedinPACE | MZ | .28*** [.13, .42] | | .21** [.07, .35] | | .59*** [.43, .72] | | .44*** [.28, .59] | |
|  | DZ | .18* [.03, .33] | | .07 [-.07, .21] | | .23 [-.01, .44] | | -.05 [-.46, .38] | |

*Note.* The estimates refer to Spearman's correlations*.* The rank-order stability within pairs of twins represents the stability of differences between individuals across two measurement occasions (in a pair of twins *t* and *u*: twin *t*, timepoint 1 – twin *t*, timepoint 2 and twin *u*, timepoint 1 – twin *u*, timepoint 2). The rank-order stability between twins and co-twins represents the twin correlations across the two measurement occasions (in a pair of twins *t* and *u*: twin *t*, timepoint 1 – twin *u*, timepoint 2 and twin *u*, timepoint 1 – twin *t*, timepoint 2). MZ, monozygotic; DZ, dizygotic; *, *p* < .05; **, *p* < .01; ***, *p* < .001. 95% confidence intervals are shown in brackets.

**Table S5**

*Goodness-of-fit statistics of the nested model in the saturated model for epigenetic aging measures*

| Model specification | ep | -2LL | *df* | AIC | *p* |
| --- | --- | --- | --- | --- | --- |
| Horvath Acceleration (univariate twin modeling) | | | | | |
| Saturated model | 10 | 10 415.49 | 1 942 | 10 435.49 |  |
| mT1=mT2 | 8 | 10 422.80 | 1 944 | 10 438.80 | .026 |
| mT1=mT2 & varT1=varT2 | 6 | 10 423.34 | 1 946 | 10 435.34 | .097 |
| Zygosity MZ=DZ | 4 | 10 424.09 | 1 948 | 10 432.09 | .198 |
| PedBE Acceleration (univariate twin modeling) | | | | | |
| Saturated model | 10 | 5 676.84 | 1 942 | 5 696.84 |  |
| mT1=mT2 | 8 | 5 677.86 | 1 944 | 5 693.86 | .601 |
| mT1=mT2 & varT1=varT2 | 6 | 5 680.27 | 1 946 | 5 692.27 | .488 |
| Zygosity MZ=DZ | 4 | 5 682.94 | 1 948 | 5 690.94 | .412 |
| GrimAge Acceleration (univariate twin modeling) | | | | | |
| Saturated model | 10 | 8 796.82 | 1 942 | 8 816.82 |  |
| mT1=mT2 | 8 | 8 798.90 | 1 944 | 8 814.90 | .353 |
| mT1=mT2 & varT1=varT2 | 6 | 8 803.84 | 1 946 | 8 815.84 | .135 |
| Zygosity MZ=DZ | 4 | 8 804.94 | 1 948 | 8 812.94 | .230 |
| DunedinPACE (univariate twin modeling) | | | | | |
| Saturated model | 10 | -4 556.84 | 1 942 | -4 536.84 |  |
| mT1=mT2 | 8 | -4 549.84 | 1 944 | -4 533.84 | .030 |
| mT1=mT2 & varT1=varT2 | 6 | -4 548.88 | 1 946 | -4 536.88 | .093 |
| Zygosity MZ=DZ | 4 | -4 546.53 | 1 948 | -4 538.53 | .112 |
| Horvath Acceleration (bivariate twin modeling) | | | | | |
| Saturated model | 28 | 10 115.38 | 1 924 | 10 171.38 |  |
| mT1=mT2 | 24 | 10 123.03 | 1 928 | 10 171.03 | .105 |
| mT1=mT2 & varT1=varT2 | 20 | 10 124.36 | 1 932 | 10 164.36 | .343 |
| Zygosity MZ=DZ | 16 | 10 125.44 | 1 936 | 10 157.44 | .610 |
| PedBE Acceleration (bivariate twin modeling) | | | | | |
| Saturated model | 28 | 5 366.26 | 1 924 | 5 422.26 |  |
| mT1=mT2 | 24 | 5 367.73 | 1 928 | 5 415.73 | .833 |
| mT1=mT2 & varT1=varT2 | 20 | 5 370.38 | 1 932 | 5 410.38 | .846 |
| Zygosity MZ=DZ | 16 | 5 372.32 | 1 936 | 5 404.32 | .913 |
| GrimAge Acceleration (bivariate twin modeling) | | | | | |
| Saturated model | 28 | 8 679.91 | 1 924 | 8 735.91 |  |
| mT1=mT2 | 24 | 8 685.18 | 1 928 | 8 733.18 | .261 |
| mT1=mT2 & varT1=varT2 | 20 | 8 691.69 | 1 932 | 8 731.69 | .161 |
| Zygosity MZ=DZ | 16 | 8 693.46 | 1 936 | 8 725.46 | .330 |
| DunedinPACE (bivariate twin modeling) | | | | | |
| Saturated model | 28 | -3 566.40 | 1 924 | -3 510.40 |  |
| mT1=mT2 | 24 | -3 559.32 | 1 928 | -3 511.32 | .132 |
| mT1=mT2 & varT1=varT2 | 20 | -3 559.32 | 1 932 | -3 519.32 | .528 |
| Zygosity MZ=DZ | 16 | -3 558.84 | 1 936 | -3 526.84 | .819 |

*Note.* mT1=mT2, expected means to be equal across co-twins; varT1=varT2, expected variances to be equal across twin order, Zygosity MZ=DZ, expected means and variances to be equal across twin order and zygosity; ep, the number of estimated parameters; -2LL, log-likelihood statistics; *df*, degrees of freedom; AIC, Akaike information criterion; *p*, results of the comparison of a parsimonious model with a more complex model based on the chi-square difference test.

**Table S6**

*Goodness-of-fit statistics for univariate twin models of variance in epigenetic aging measures (976 twin pairs)*

| Comparison | ep | -2LL | *df* | AIC | *p* | 1 − β |
| --- | --- | --- | --- | --- | --- | --- |
| Horvath Acceleration | | | | | | |
| **ACE model** | **4** | **10 424.09** | **1 948** | **10 432.09** |  |  |
| AE model | 3 | 10 432.86 | 1 949 | 10 438.86 | .003 | .91 |
| CE model | 3 | 10 430.41 | 1 949 | 10 436.41 | .012 | .81 |
| E model | 2 | 10 648.19 | 1 950 | 10 652.19 | < .001 | 1 |
| PedBE Acceleration | | | | | | |
| ACE model | 4 | 5 683.24 | 1 948 | 5 691.24 |  |  |
| AE model | 3 | 5 719.40 | 1 949 | 5 725.40 | < .001 | 1 |
| **CE model** | **3** | **5 683.24** | **1 949** | **5 689.24** | **1.00** |  |
| E model | 2 | 5 924.38 | 1 950 | 5 928.38 | < .001 | 1 |
| GrimAge Acceleration | | | | | | |
| ACE model | 4 | 8 804.94 | 1 948 | 8 812.94 |  |  |
| **AE model** | **3** | **8 805.03** | **1 949** | **8 811.03** | **.766** |  |
| CE model | 3 | 8 810.85 | 1 949 | 8 816.85 | .015 | .78 |
| E model | 2 | 8 874.16 | 1 950 | 8 878.16 | < .001 | 1 |
| DunedinPACE | | | | | | |
| ADE model | 4 | -4 546.53 | 1 948 | -4 538.53 |  |  |
| **AE model** | **3** | **-4 545.14** | **1 949** | **-4 539.14** | **1.00** |  |
| E model | 2 | -4 392.87 | 1 950 | -4 388.87 | < .001 | 1 |

*Note.* *A*, additive genetic factors; *C*, shared environmental factors; *D*, non-additive genetic factors; *E*, individually-unique environmental factors including measurement error; ep, the number of estimated parameters; -2LL, log-likelihood statistics; *df*, degrees of freedom; AIC, Akaike information criterion; *p*, results of the comparison of a parsimonious model with a more complex model based on the chi-square difference test; 1 *–* β, achieved statistical power. The respective best fitting or most parsimonious model that did not fit the data worse compared to less restricted models is shown in bold.

**Table S7**

*Standardized variance components and confidence intervals from univariate twin models*

| Epigenetic aging measure | Model | *A* % [90% CI] | *C/D* % [90% CI] | *E* % [90% CI] |
| --- | --- | --- | --- | --- |
| Horvath Accel. | Full model | 24 [6, 43] | 26 [9, 42] | 50 [44, 56] |
|  | Best-fitting model | 24 [6, 43] | 26 [9, 42] | 50 [44, 56] |
| PedBE Accel. | Full model | 0 [0, 14] | 47 [42, 52] | 53 [48, 58] |
|  | Best-fitting model | - | 47 [42, 52] | 53 [48, 58] |
| GrimAge Accel. | Full model | 28 [7, 39] | 3 [0, 22] | 69 [61, 76] |
|  | Best-fitting model | 32 [25, 39] | - | 68 [61, 75] |
| DunedinPACE | Full model | 27 [0, 52] | 22 [0, 53] | 51 [46, 58] |
|  | Best-fitting model | 47 [41, 53] | - | 53 [47, 59] |

*Note*. The estimates are derived from the full and best-fitting univariate twin models for the four epigenetic aging measures. *A* %, relative contribution of the additive genetic factors; *D* %, relative contribution of the non-additive genetic factors; *C* %, relative contribution of the shared environmental factors; *E* %, relative contribution of the individually-unique environmental factors including measurement error; [90% CI], 90% confidence intervals; Accel., Acceleration.

**Table S8**

*Goodness-of-fit statistics for bivariate twin models of epigenetic aging measures (488 twin pairs)*

| Comparison | ep | -2LL | *df* | AIC | *p* | 1 − *β* |
| --- | --- | --- | --- | --- | --- | --- |
| Horvath Acceleration | | | | | | |
| ACE model | 11 | 10 127.74 | 1 941 | 10 149.74 |  | .41 |
| ACE model with *a*²*1*=*a*²*2*, *c*²*1*=*c*²*2*, and *e*²*1*=*e*²*2* variances | 8 | 10 142.03 | 1 944 | 10 158.03 | .003 | 1 |
| **ACE model with *a*²*1*=*a*²*2* variances** | **10** | **10 127.75** | **1 942** | **10 147.75** | **.945** |  |
| ACE model with *c*²*1*=*c*²*2* variances | 10 | 10 128.38 | 1 942 | 10 148.38 | .427 |  |
| ACE model with *e*²*1*=*e*²*2* variances | 10 | 10 130.66 | 1 942 | 10 150.66 | .088 | .52 |
| ACE model with *a*²*1*=*a*²*2* and *c*²*1*=*c*²*2* variances | 9 | 10 131.89 | 1 943 | 10 149.89 | .126 | .43 |
| AE model | 8 | 10 135.55 | 1 944 | 10 151.55 | .050 | .62 |
| CE model | 8 | 10 136.12 | 1 944 | 10 152.12 | .039 | .67 |
| E model | 5 | 10 321.01 | 1 947 | 10 331.01 | < .001 | 1 |
| PedBE Acceleration | | | | | | |
| ACE model | 11 | 5 390.70 | 1 941 | 5 412.70 |  | .40 |
| AE model | 8 | 5 416.91 | 1 944 | 5 432.91 | < .001 | 1 |
| **CE model** | **8** | **5 394.75** | **1 944** | **5 410.75** | **.256** |  |
| CE model with *e*²*1*=*e*²*2* variances | 7 | 5 403.72 | 1 945 | 5 417.72 | .003 | .84 |
| CE model with *c*²*1*=*c*²*2* variances | 7 | 5 399.04 | 1 945 | 5 413.04 | .038 | .45 |
| E model | 5 | 5 598.51 | 1 947 | 5 608.51 | < .001 | 1 |
| GrimAge Acceleration | | | | | | |
| ACE model | 11 | 8 700.78 | 1 941 | 8 722.78 |  | .85 |
| AE model | 8 | 8 701.44 | 1 944 | 8 717.44 | .884 | .39 |
| AE model with *e*²*1*=*e*²*2* variances | 7 | 8 705.89 | 1 945 | 8 719.89 | .035 | .67 |
| **AE model with *a*²*1*=*a*²*2* variances** | **7** | **8 701.56** | **1 945** | **8 715.56** | **.728** |  |
| CE model | 8 | 8 709.28 | 1 944 | 8 725.28 | .037 | .93 |
| E model | 5 | 8 775.54 | 1 947 | 8 785.54 | < .001 | 1 |
| DunedinPACE | | | | | | |
| ADE model | 11 | -4 751.77 | 1 941 | -4 729.77 |  | .76 |
| AE model | 8 | -4 749.24 | 1 944 | -4 733.24 | .470 | .41 |
| AE model with *e*²*1*=*e*²*2* variances | 7 | -4 746.88 | 1 945 | -4 732.88 | .125 | .46 |
| **AE model with *a***²***1*=*a***²***2* variances** | **7** | **-4 749.24** | **1 945** | **-4 735.24** | **.953** |  |
| E model | 5 | -4 614.29 | 1 947 | -4 604.29 | < .001 | 1 |

*Note.* *A/a*², additive genetic factor/factor’s variances; *D/d*², non-additive genetic factor/factor’s variances; *C/c*², shared environmental factor/factor’s variances; *E/e*², individually-unique environmental factor/factor’s variances including measurement error; ep, the number of estimated parameters; -2LL, log-likelihood statistics; *df*, degrees of freedom; AIC, Akaike information criterion; *p*, results of the comparison of a parsimonious model with a more complex model based on the chi-square difference test; 1 *–* β, achieved statistical power. The parsimonious model that did not fit the data worse compared to less restricted models, but better than more restricted models is shown in bold. Models dropping single variance components were compared to the full model, while models constraining paths to be equal were compared to the most parsimonious models that did not fit the data worse than more complex models.

**Table S9**

*Variance components and confidence intervals derived from bivariate twin models (488 twin pairs)*

| Epigenetic aging measure | Model | Measurement | VA *(A* %  [90% CI]) | VC/VD *(C/D* %  [90% CI]) | VE *(E* %  [90% CI]) |
| --- | --- | --- | --- | --- | --- |
| Horvath Accel. | Full model | First measurement | 3.53 (29 [3, 56]) | 2.59 (21 [-4, 43]) | 6.13 (50 [43, 59]) |
|  |  | Covariance over time | 3.09 (43 [8, 81]) | 3.07 (43 [9, 73]) | 0.99 (14 [4, 25]) |
|  |  | Second measurement | 3.36 (22 [-3, 48]) | 4.28 (28 [5, 49]) | 7.46 (49 [42, 58]) |
|  | Best-fitting model | First measurement | 3.46 (28 [8, 50]) | 2.65 (22 [1, 40]) | 6.14 (50 [43, 58]) |
|  |  | Covariance over time | 3.10 (43 [8, 82]) | 3.07 (43 [9, 73]) | 0.99 (14 [3, 25]) |
|  |  | Second measurement | 3.46 (23 [6, 41]) | 4.20 (28 [11, 43]) | 7.45 (49 [42, 57]) |
| PedBE Accel.  PedBE Accel. | Full model | First measurement | 0.18 (17 [-8, 42]) | 0.36 (34 [12, 54]) | 0.52 (49 [41, 58]) |
|  |  | Covariance over time | 0.15 (24 [-11, 59]) | 0.38 (60 [29, 87]) | 0.11 (17 [6, 29]) |
|  |  | Second measurement | -0.12 (-9 [-33, 17]) | 0.74 (54 [32, 73]) | 0.75 (55 [46, 64]) |
|  | Best-fitting model | First measurement | - | 0.50 (47 [40, 54]) | 0.56 (53 [46, 60]) |
|  |  | Covariance over time | - | 0.49 (78 [69, 87]) | 0.14 (22 [13, 31]) |
|  |  | Second measurement | - | 0.64 (47 [39, 53]) | 0.73 (53 [47, 61]) |
| GrimAge Accel. | Full model | First measurement | 1.93 (37 [6, 70]) | -0.11 (-2 [-30, 24]) | 3.33 (65 [55, 75]) |
|  |  | Covariance over time | 1.51 (91 [14, 100]) | -0.07 (-4 [-73, 60]) | 0.23 (14 [-13, 38]) |
|  |  | Second measurement | 1.01 (17 [-15, 50]) | 0.56 (10 [-18, 36]) | 4.30 (73 [63, 85]) |
|  | Best-fitting model | First measurement | 1.75 (34 [26, 42]) | - | 3.38 (66 [58, 74]) |
|  |  | Covariance over time | 1.43 (86 [63, 110]) | - | 0.24 (14 [-10, 37]) |
|  |  | Second measurement | 1.75 (30 [23, 36]) | - | 4.15 (70 [64, 77]) |
| Dunedin  PACE  Dunedin  PACE | Full model | First measurement | 0.0019 (32 [-17, 78]) | 0.0011 (18 [-29, 70]) | 0.0029 (49 [41, 59]) |
|  |  | Covariance over time | 0.0004 (15 [-73, 96]) | 0.0019 (68 [-17, NA]) | 0.0005 (18 [5, 32]) |
|  |  | Second measurement | 0.0012 (19 [-34, 69]) | 0.0018 (28 [-23, 83]) | 0.0034 (53 [45, 62]) |
|  | Best-fitting model | First measurement | 0.00299 (50 [43, 57]) | - | 0.00294 (50 [43, 57]) |
|  |  | Covariance over time | 0.00222 (79 [65, 91]) | - | 0.00059 (21 [9, 35]) |
|  |  | Second measurement | 0.00299 (46 [39, 53]) | - | 0.00350 (54 [47, 61]) |

*Note*. The estimates of the unstandardized variance components, standardized variance components, and their confidence intervals are derived from the full and the best-fitting bivariate twin models across measurement occasions for the four epigenetic aging measures. VA, VD, VC, and VE, the unstandardized variance components for additive genetic, non-additive genetic, shared environmental, and individually-unique environmental factors (the latter including measurement error) correspondingly; *A* %, *D* %, *C* %, and *E* %, the relative contribution of additive genetic, non-additive genetic, shared environmental, and individually-unique environmental factors including measurement error; [90% CI], 90% confidence intervals.

**Table S10**

*Goodness-of-fit statistics for univariate twin models with age as a moderator (976 twin pairs)*

| Comparison | ep | -2LL | *df* | AIC | *p* | 1 − β |
| --- | --- | --- | --- | --- | --- | --- |
| Horvath Acceleration | | | | | | |
| Moderated ACE model | 9 | 10 338.95 | 1 943 | 10 356.95 |  | .38 |
| Non-moderated *a* effect in ACE model | 8 | 10 343.80 | 1 944 | 10 359.80 | .028 | .69 |
| **Non-moderated *c* effect in ACE model** | **8** | **10 339.16** | **1 944** | **10 355.16** | **.646** |  |
| Non-moderated *e* effect in ACE model | 8 | 10 373.90 | 1 944 | 10 389.90 | < .001 | 1 |
| Moderated AE model | 7 | 10 349.21 | 1 945 | 10 363.21 | .006 | .88 |
| Non-moderated ACE model | 6 | 10 420.41 | 1 946 | 10 432.41 | < .001 | 1 |
| PedBE Acceleration | | | | | | |
| Moderated ACE model | 9 | 5 534.79 | 1 943 | 5 552.79 |  | .63 |
| Non-moderated *a* effect in ACE model | 8 | 5 534.80 | 1 944 | 5 550.80 | .946 | .40 |
| Non-moderated *c* effect in ACE model | 8 | 5 534.85 | 1 944 | 5 550.85 | .806 | .41 |
| Non-moderated *e* effect in ACE model | 8 | 5 630.07 | 1 944 | 5 646.06 | < .001 | 1 |
| **Non-moderated *a* and *c* effects in ACE model** | **7** | **5 534.89** | **1 945** | **5 548.89** | **.953** |  |
| Moderated AE model | 7 | 5 562.17 | 1 945 | 5 576.17 | < .001 | 1 |
| Moderated CE model | 7 | 5 537.94 | 1 945 | 5 551.94 | .207 | .54 |
| Non-moderated *c* effect in CE model | 6 | 5 538.13 | 1 946 | 5 550.13 | .342 | .30 |
| Non-moderated ACE model | 6 | 5 650.11 | 1 946 | 5 662.11 | < .001 | 1 |
| GrimAge Acceleration | | | | | | |
| Moderated ACE model | 9 | 8 793.38 | 1 943 | 8 811.38 |  | .73 |
| Non-moderated *a* effect in ACE model | 8 | 8 794.15 | 1 944 | 8 810.15 | .382 | .63 |
| Non-moderated *c* effect in ACE model | 8 | 8 793.39 | 1 944 | 8 809.39 | .926 | .55 |
| Non-moderated *e* effect in ACE model | 8 | 8 794.15 | 1 944 | 8 810.15 | .382 | .63 |
| Moderated CE model | 7 | 8 798.92 | 1 945 | 8 812.92 | .063 | .82 |
| Moderated AE model | 7 | 8 793.47 | 1 945 | 8 807.47 | .958 | .29 |
| Non-moderated *a* effect in AE model | 6 | 8 796.57 | 1 946 | 8 808.57 | .363 | .45 |
| **Non-moderated *e*** **effect in AE model** | **6** | **8 794.28** | **1 946** | **8 806.28** | **.826** |  |
| Non-moderated ACE model | 6 | 8 802.06 | 1 946 | 8 814.06 | .034 | .87 |
| Non-moderated AE model | 5 | 8 802.13 | 1 947 | 8 812.13 | .068 | .78 |
| DunedinPACE | | | | | | |
| Moderated ADE model | 9 | -4 571.98 | 1 943 | -4 553.98 |  | .56 |
| Moderated AE model | 7 | -4 464.40 | 1 945 | -4 450.40 | < .001 | 1 |
| Non-moderated *a* effect in ADE model | 8 | -4 571.45 | 1 944 | -4 555.45 | .467 | .37 |
| Non-moderated *d* effect in ADE model | 8 | -4 570.15 | 1 944 | -4 554.15 | .176 | .54 |
| Non-moderated *e* effect in ADE model | 8 | -4 571.67 | 1 944 | -4 555.67 | .581 | .34 |
| Non-moderated *a* and *d* effects in ADE model | 7 | -4 567.48 | 1 945 | -4 553.48 | .105 | .61 |
| **Non-moderated *a* and *e* effects in ADE model** | **7** | **-4 571.20** | **1 945** | **-4 557.20** | **.676** |  |
| Non-moderated *d* and *e* effects in ADE model | **7** | -4 569.56 | 1 945 | -4 555.56 | .298 | .36 |
| Non-moderated ADE model | 6 | -4 563.85 | 1 946 | -4 551.85 | .043 | .75 |
| Non-moderated AE model | 5 | -4 562.38 | 1 947 | -4 552.38 | .048 | .71 |

*Note.* Goodness-of-fit statistics are presented for univariate twin models with age-moderated paths, quadratic, and linear main effects of age. *A/a,* additive genetic factor/effect; *D/d,* non-additive genetic factor/effect; *C/c,* shared environmental factor/effect; *E/e*, individually-unique environmental factor/effect including measurement error; ep, the number of estimated parameters; -2LL, log-likelihood statistics; *df*, degrees of freedom; AIC, Akaike information criterion; *p,* results of the comparison of a parsimonious model with a more complex model based on the chi-square difference test; 1 *–* β, achieved statistical power. The parsimonious model that did not fit the data worse compared to less restricted models but better than more restricted models is shown in bold.

**Table S11**

*Variance components and confidence intervals derived from univariate twin models with age as a moderator*

| Epigenetic aging measure | Age group | *A* %  [90% CI] | *C/D* %  [90% CI] | *E* %  [90% CI] | *V* |
| --- | --- | --- | --- | --- | --- |
| Horvath Accel. | 9.5 | **18 [3, 41]** | 35 [23, 53] | **47 [39, 57]** | 8.82 |
|  | 12 | **22 [6, 42]** | 30 [20, 46] | **48 [42, 56]** | 10.14 |
|  | 15.5 | **26 [11, 44]** | 25 [17, 38] | **49 [44, 55]** | 12.24 |
|  | 18 | **29 [14, 46]** | 22 [15, 34] | **49 [44, 56]** | 13.92 |
|  | 21.5 | **32 [17, 49]** | 19 [12, 28] | **49 [43, 57]** | 16.54 |
|  | 24 | **35 [18, 52]** | 16 [11, 25] | **49 [42, 58]** | 18.59 |
|  | 27.5 | **37 [20, 56]** | 14 [9, 22] | **49 [40, 60]** | 21.72 |
|  | 30 | **39 [20, 59]** | 12 [8, 19] | **49 [39, 61]** | 24.14 |
| PedBE Accel. | 9.5 | 18 [0, 38] | 49 [32, 66] | **33 [27, 41]** | 0.88 |
|  | 12 | 16 [0, 34] | 45 [29, 61] | **39 [33, 46]** | 0.96 |
|  | 15.5 | 14 [0, 30] | 40 [26, 53] | **46 [41, 52]** | 1.09 |
|  | 18 | 13 [0, 28] | 36 [24, 49] | **51 [45, 57]** | 1.19 |
|  | 21.5 | 11 [0, 24] | 32 [21, 43] | **57 [50, 64]** | 1.35 |
|  | 24 | 11 [0, 22] | 29 [19, 39] | **60 [53, 69]** | 1.48 |
|  | 27.5 | 9 [0, 20] | 26 [17, 35] | **65 [56, 75]** | 1.67 |
|  | 30 | 8 [0, 18] | 24 [15, 32] | **68 [58, 80]** | 1.82 |
| GrimAge Accel. | 9.5 | **24 [14, 35]** | - | 76 [69, 85] | 4.95 |
|  | 12 | **26 [17, 36]** | - | 74 [67, 82] | 5.11 |
|  | 15.5 | **29 [21, 38]** | - | 71 [64, 79] | 5.35 |
|  | 18 | **32 [24, 40]** | - | 68 [61, 76] | 5.54 |
|  | 21.5 | **35 [27, 44]** | - | 65 [58, 72] | 5.82 |
|  | 24 | **37 [28, 48]** | - | 63 [56, 70] | 6.03 |
|  | 27.5 | **40 [29, 55]** | - | 60 [54, 66] | 6.35 |
|  | 30 | **43 [29, 60]** | - | 57 [52, 64] | 6.59 |
| DunedinPACE | 9.5 | 32 [0, 53] | **10 [0, 10]** | 58 [52, 66] | 0.005 |
|  | 12 | 30 [0, 51] | **13 [0, 50]** | 57 [50, 64] | 0.006 |
|  | 15.5 | 29 [0, 49] | **17 [0, 52]** | 54 [48, 61] | 0.006 |
|  | 18 | 28 [0, 47] | **20 [1, 53]** | 52 [46, 59] | 0.006 |
|  | 21.5 | 27 [0, 45] | **24 [3, 57]** | 49 [44, 56] | 0.006 |
|  | 24 | 26 [0, 43] | **27 [6, 59]** | 47 [42, 54] | 0.007 |
|  | 27.5 | 24 [0, 40] | **31 [8, 62]** | 45 [40, 51] | 0.007 |
|  | 30 | 23 [0, 39] | **34 [10, 64]** | 43 [38, 48] | 0.007 |

*Note.* The estimates are derived from the best-fitting univariate twin models with age as a moderator in a pooled sample for four epigenetic aging measures. *A* %, relative contribution of the additive genetic factors; *D* %, relative contribution of the non-additive genetic factors; *C* %, relative contribution of the shared environmental factors; *E* %, relative contribution of the individually-unique environmental factors including measurement error; *V*, total variance; [90% CI], 90% confidence intervals; Accel., Acceleration. Components that are significantly moderated by age are shown in bold.

**Figure S4**

*Variance decomposition models fitted in the performed analyses*

a) *Univariate twin model*

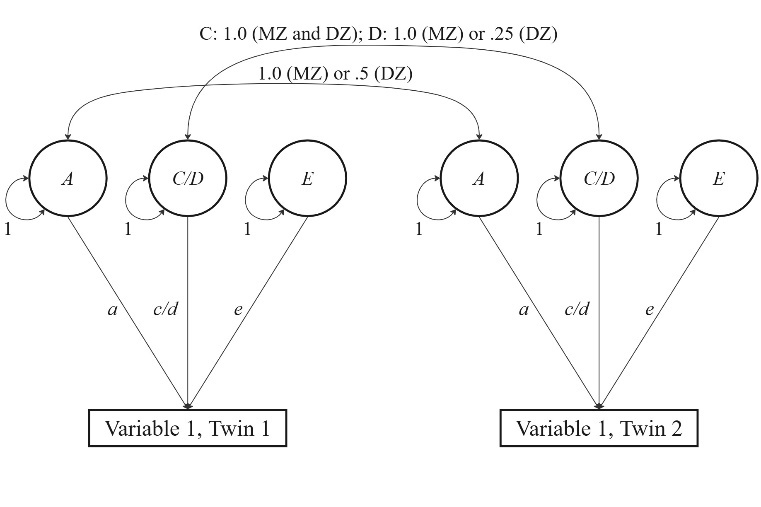

*Note.* Univariate model for data from monozygotic (MZ) and dizygotic (DZ) twins reared together. *A/a*, additive genetic factor/effect; *D/d*, non-additive genetic factor/effect; *C/c*, shared environmental factor/effect; *E/e*, individually-unique environmental factor/effect including measurement error. Variances of genetic and environmental factors *A*, *D*, *C*, and *E* are fixed to one in order to estimate path coefficients (effects), which allows estimations of the relative contributions of genetic (*a*² and *d*²) and environmental influences (*c*² and *e*²) to the variance in a measured variable.

b) *Bivariate twin model* *across* *measurement* *occasions*

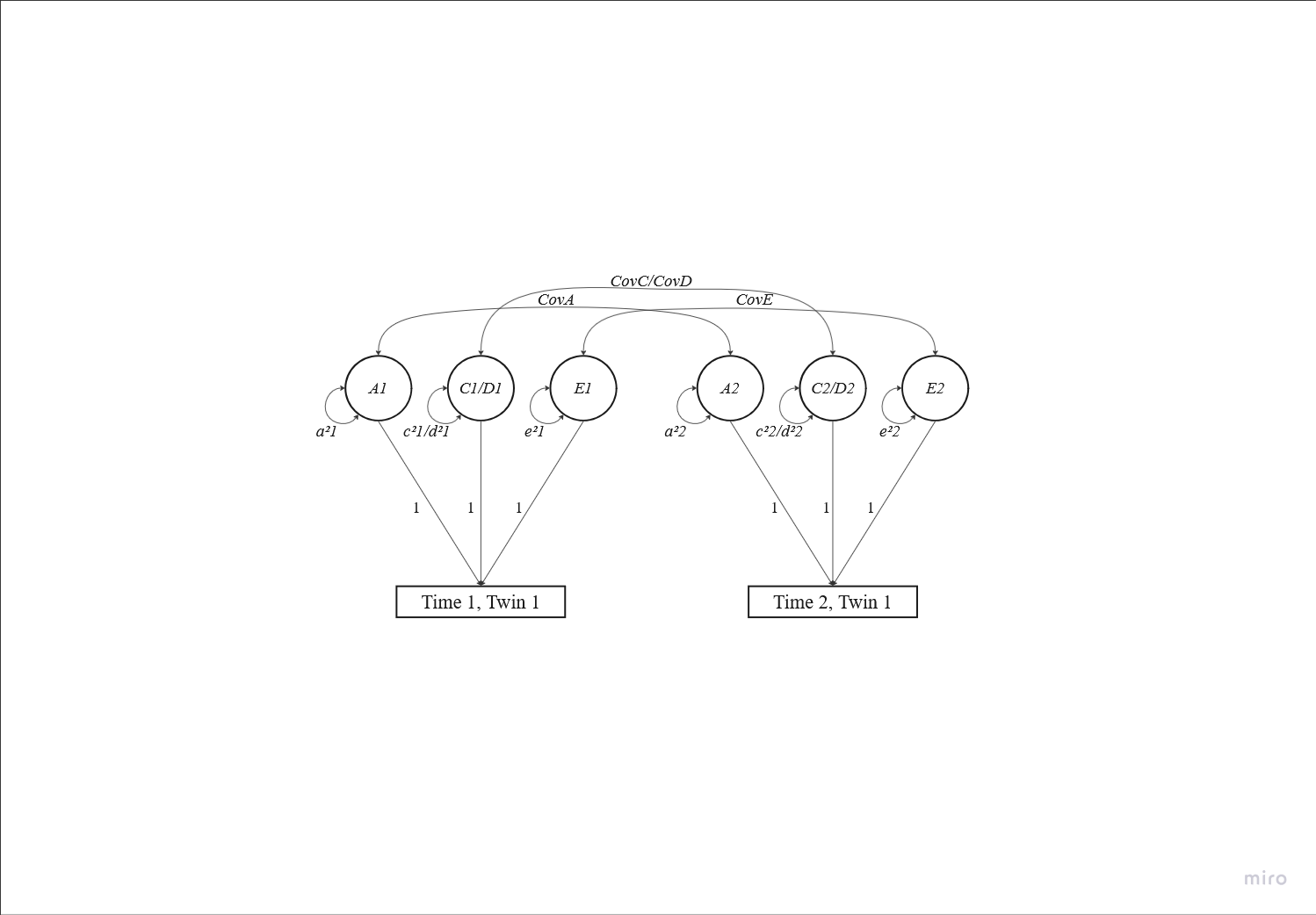

*Note.* Bivariate model for data from monozygotic (MZ) and dizygotic (DZ) twins reared together specified as a correlated factors model across timepoints 1 and 2. For simplicity, the model is only depicted for one twin. *A1/a*²*1* and *A2/a*²*2*, additive genetic factors/factors’ variances; *D1/d*²*1* and *D2/d*²*2*, non-additive genetic factors/factors’ variances; *C1/c*²*1* and *C2/c*²*2*, shared environmental factors/factors’ variances; *E1/e*²*1* and *E2/e*²*2*, individually-unique environmental factors/factors’ variances including measurement error; path coefficients (effects) of genetic and environmental factors *A1*, *A2*, *C1*, *C2*, *D1*, *D2*, *E1* and *E2* are fixed to one in order to estimate the variance components. This model allows estimations of the genetic (*a*² and *d*²) and environmental (*c*² and *e*²) variance components at time 1 and 2 as well as cross-time covariances of additive and non-additive genetic factors (*CovA*, *CovD*) and shared and individually-unique environmental factors (*CovC*, *CovE*).

c) *Univariate twin model with age as a continuous moderator*

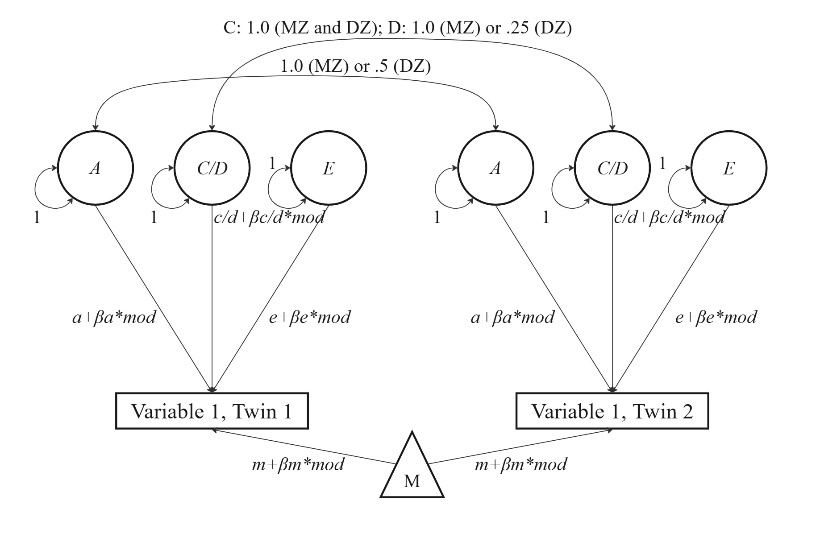

*Note.* Univariate model with age as continuous moderator for data from monozygotic (MZ) and dizygotic (DZ) twins reared together. *A/a*, additive genetic factor/effect; *D/d*, non-additive genetic factor/effect; *C/c*, shared environmental factor/effect; *E/e*, individually-unique environmental factor/effect incl. measurement error; variances of genetic and environmental factors *A*, *D*, *C*, and *E* are fixed to one in order to estimate path coefficients (effects); M, moderator; *m*, means; *βm*mod,* effect of the moderator on the variable, *βa*mod, βc/d*mod,* and *βe*mod*, the moderated effects of the corresponding genetic and environmental factors.

**Figure S5**

*Box plots with mean-levels and standard deviations of epigenetic aging across age groups*

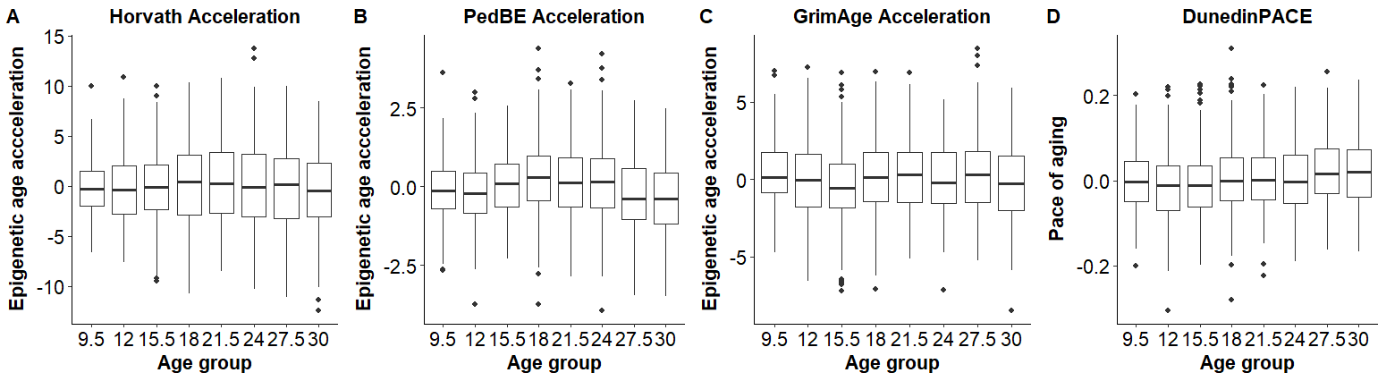

**Figure S6**

*Cross-cohort standardized genetic and environmental variance components of epigenetic aging measures* *in the full model*

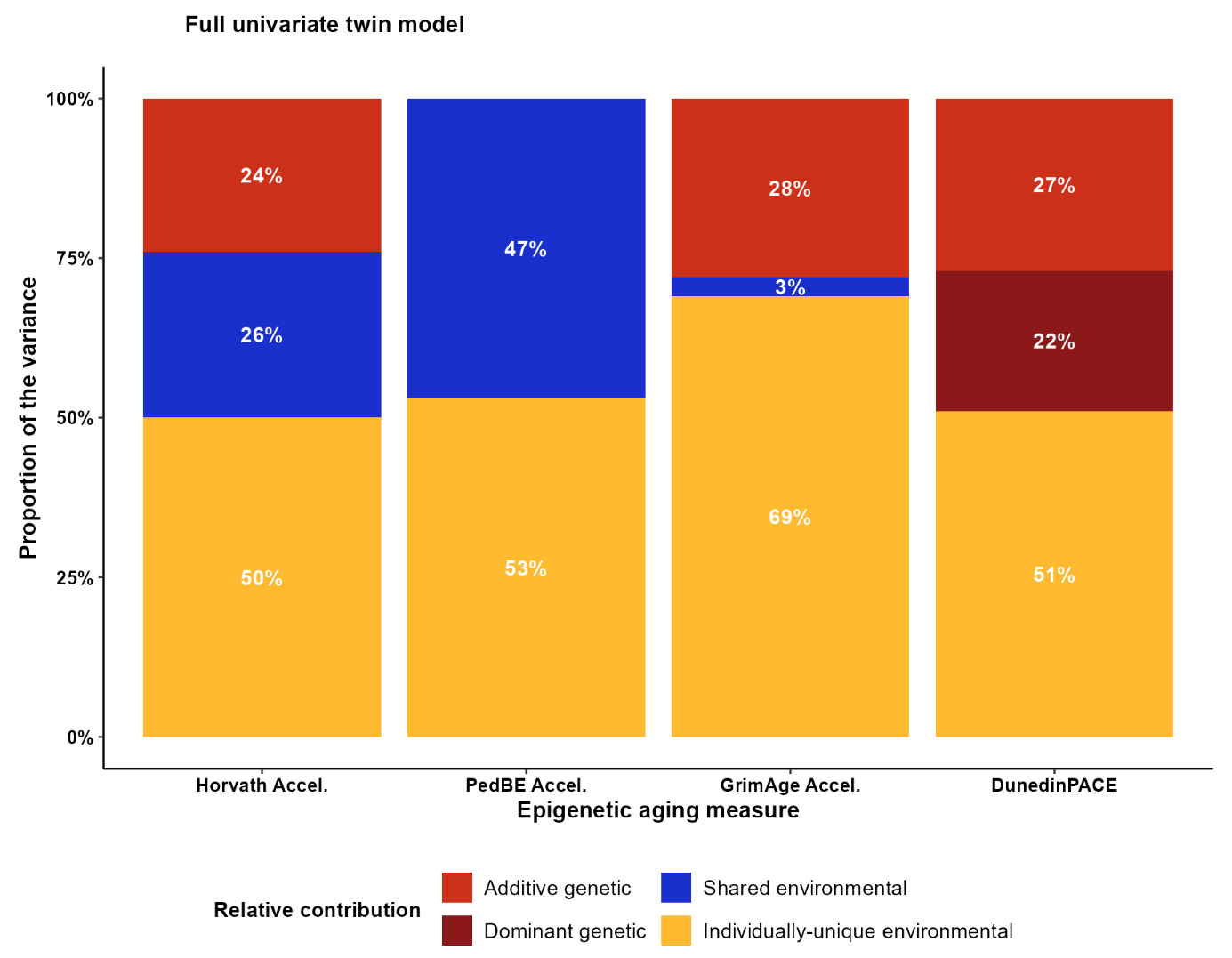

*Note.* The estimates are derived from the full models in the univariate twin model analysis.

**Figure S7**

*Unstandardized genetic and environmental variance components of epigenetic aging measures over 2.5 years*
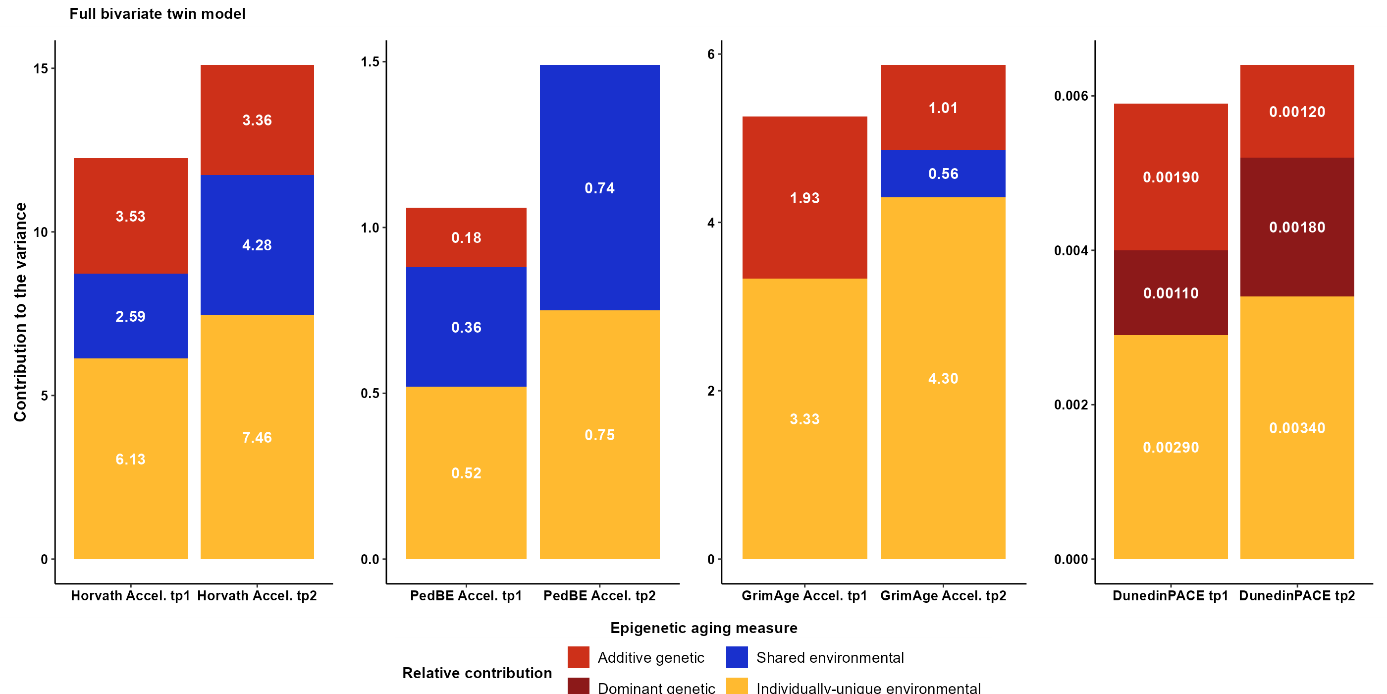

*Note.* The estimates are derived from the full bivariate twin models. tp.1, timepoint 1; tp. 2, timepoint 2.

**Figure S8**

*Unstandardized variance components and their confidence intervals derived from the univariate twin models with age as a moderator*

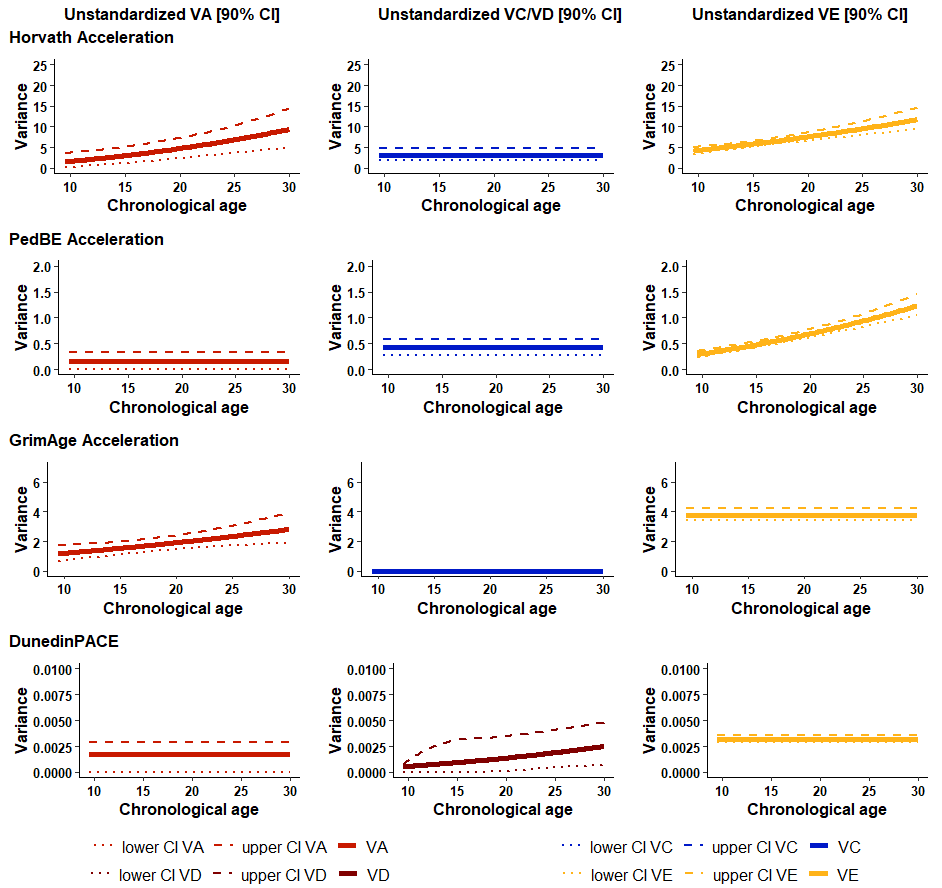

*Note.* The estimates are derived from the best-fitting univariate twin models with age as a moderator in a pooled sample. VA, VD, VC, and VE represent the unstandardized variance components for additive genetic, non-additive genetic, shared environmental, and individually-unique environmental factors (the latter including measurement error) correspondingly; [90% CI], 90% confidence intervals.
