## Supplementary material for "Genetic and Environmental Contributions to Epigenetic Aging Across Adolescence and Young Adulthood": Sensitivity analysis

**Additional file 4: Sensitivity analysis**

**Table S12**

*Goodness-of-fit statistics for univariate models of epigenetic aging measures* ***unadjusted for biological sex*** *(976 twin pairs)*

| Comparison | ep | -2LL | *df* | AIC | *p* | 1 − β |
| --- | --- | --- | --- | --- | --- | --- |
| Horvath Acceleration | | | | | | |
| ***ACE model*** | ***4*** | ***10 424.55*** | ***1 948*** | ***10 432.55*** |  |  |
| AE model | 3 | 10 433.43 | 1 949 | 10 439.43 | .003 | .83 |
| CE model | 3 | 10 430.83 | 1 949 | 10 436.83 | .012 | .66 |
| E model | 2 | 10 648.99 | 1 950 | 10 652.99 | < .001 | 1 |
| PedBE Acceleration | | | | | | |
| ACE model | 4 | 5 683.31 | 1 948 | 5 691.31 |  | .41 |
| AE model | 3 | 5 719.22 | 1 949 | 5 725.22 | < .001 | 1 |
| ***CE model*** | ***3*** | ***5 683.31*** | ***1 949*** | ***5 689.31*** | ***1.00*** |  |
| E model | 2 | 5 924.71 | 1 950 | 5 928.71 | < .001 | 1 |
| GrimAge Acceleration | | | | | | |
| ACE model | 4 | 8 806.11 | 1 948 | 8 814.11 |  | .40 |
| ***AE model*** | ***3*** | ***8 806.20*** | ***1 949*** | ***8 812.20*** | ***.762*** |  |
| CE model | 3 | 8 812.05 | 1 949 | 8 818.05 | .015 | .79 |
| E model | 2 | 8 875.90 | 1 950 | 8 879.90 | < .001 | 1 |
| DunedinPACE | | | | | | |
| ADE model | 4 | -4 515.84 | 1 948 | -4 507.84 |  | .30 |
| ***AE model*** | ***3*** | ***-4 515.08*** | ***1 949*** | ***-4 509.08*** | ***.383*** |  |
| E model | 2 | -4 354.30 | 1 950 | -4 350.30 | < .001 | 1 |

*Note.* *A*, additive genetic factors; *C*, shared environmental factors; *D*, non-additive genetic factors; *E*, unique environmental factors including measurement error; ep, the number of estimated parameters; -2LL, log-likelihood statistics; *df*, degrees of freedom; AIC, Akaike information criterion; *p*, results of the comparison of a parsimonious model with a more complex model based on the chi-square difference test; 1 *–* β, achieved statistical power. The parsimonious models that did not fit the data worse compared to less restricted models in the sensitivity analysis are shown in italics and those in the main analysis are shown in bold.

**Table S13**

*Standardized variance components in the best-fitting univariate models for the four epigenetic measures* ***unadjusted for biological sex***

| Epigenetic aging measure | *A* (%) | *C* (%) | *E* (%) |
| --- | --- | --- | --- |
| Horvath Accel. | 24 | 26 | 50 |
| PedBE Accel. | - | 47 | 53 |
| GrimAge Accel. | 32 | - | 68 |
| DunedinPACE | 48 | - | 52 |

*Note*. The estimates are derived from the best-fitting univariate twin models for the four epigenetic aging measures. *A*, relative contribution of the additive genetic factors; *C*, relative contribution of the shared environmental factors; *E*, relative contribution of the unique environmental factors including measurement error; Accel., Acceleration.

**Table S14**

*Goodness-of-fit statistics for bivariate models of epigenetic aging measures* ***unadjusted for biological sex*** *(488 twin pairs)*

| Comparison | ep | -2LL | *df* | AIC | *p* | 1 − β |
| --- | --- | --- | --- | --- | --- | --- |
| Horvath Acceleration | | | | | | |
| ACE model | 11 | 10 127.96 | 1 941 | 10 149.96 |  | .41 |
| ACE model with *a*²*1*=*a*²*2*, *c*²*1*=*c*²*2*, and *e*²*1*=*e*²*2* variances | 8 | 10 142.24 | 1 944 | 10 158.24 | .003 | .95 |
| ***ACE model with a*²*1=a*²*2 variances*** | ***10*** | ***10 127.97*** | ***1 942*** | ***10 147.97*** | ***.935*** |  |
| ACE model with *c*²*1*=*c*²*2* variances | 10 | 10 128.61 | 1 942 | 10 148.61 | .420 | .21 |
| ACE model with *e*²*1*=*e*²*2* variances | 10 | 10 130.90 | 1 942 | 10 150.90 | .087 | .53 |
| ACE model with *a*²*1*=*a*²*2* and *c*²*1*=*c*²*2* variances | 9 | 10 132.11 | 1 943 | 10 150.11 | .126 | .43 |
| AE model | 8 | 10 135.84 | 1 944 | 10 151.84 | .049 | .63 |
| CE model | 8 | 10 136.33 | 1 944 | 10 152.33 | .039 | .67 |
| E model | 5 | 10 321.56 | 1 947 | 10 331.56 | < .001 | 1 |
| PedBE Acceleration | | | | | | |
| ACE model | 11 | 5 390.66 | 1 941 | 5 412.66 |  | .39 |
| AE model | 8 | 5 416.72 | 1 944 | 5 432.72 | < .001 | 1 |
| ***CE model*** | ***8*** | ***5 394.79*** | ***1 944*** | ***5 410.79*** | ***.248*** |  |
| CE model with *e*²*1*=*e*²*2* variances | 7 | 5 403.80 | 1 945 | 5 417.80 | .003 | .84 |
| CE model with *c*²*1*=*c*²*2* variances | 7 | 5 399.03 | 1 945 | 5 413.03 | .039 | .44 |
| E model | 5 | 5 598.77 | 1 947 | 5 608.77 | < .001 | 1 |
| GrimAge Acceleration | | | | | | |
| ACE model | 11 | 8 701.76 | 1 941 | 8 723.76 |  | .85 |
| AE model | 8 | 8 702.42 | 1 944 | 8 718.42 | .883 | .39 |
| AE model with *e*²*1*=*e*²*2* variances | 7 | 8 706.94 | 1 945 | 8 720.94 | .034 | .67 |
| ***AE model with a*²*1=a*²*2 variances*** | ***7*** | ***8 702.57*** | ***1 945*** | ***8 716.57*** | ***.701*** |  |
| CE model | 8 | 8 710.31 | 1 944 | 8 726.31 | .036 | .93 |
| E model | 5 | 8 777.13 | 1 947 | 8 787.13 | < .001 | 1 |
| DunedinPACE | | | | | | |
| ADE model | 11 | -4 726.99 | 1 941 | -4 704.99 |  | .80 |
| AE model | 8 | -4 725.25 | 1 944 | -4 709.25 | .629 | .39 |
| AE model with *e*²*1*=*e*²*2* variances | 7 | -4 723.00 | 1 945 | -4 709.00 | .134 | .41 |
| ***AE model with a*²*1=a*²*2 variances*** | ***7*** | ***-4 725.11*** | ***1 945*** | ***-4 711.11*** | ***.706*** |  |
| E model | 5 | -4 583.65 | 1 947 | -4 573.65 | < .001 | 1 |

*Note.* *A/a*, additive genetic factor/factor’s variances; *D/d*, non-additive genetic factor/factor’s variances; *C/c*, shared environmental factor/factor’s variances; *E/e*, individually-unique environmental factor/factor’s variances including measurement error; ep, the number of estimated parameters; -2LL, log-likelihood statistics; *df*, degrees of freedom; AIC, Akaike information criterion; *p*, results of the comparison of a parsimonious model with a more complex model based on the chi-square difference test; 1 *–* β, achieved statistical power*.* The parsimonious models that did not fit the data worse compared to less restricted models in the sensitivity analysis are shown in italics and those in the main analysis are shown in bold.

**Table S15**

*Variance components in the best-fitting bivariate models for the four epigenetic measures* ***unadjusted for biological sex***

| Epigenetic aging measure | Measurement | *A* (%) | *C* (%) | *E* (%) |
| --- | --- | --- | --- | --- |
| Horvath Acceleration | First measurement | 28 | 22 | 50 |
|  | Covariance over time | 43 | 43 | 14 |
|  | Second measurement | 23 | 28 | 49 |
| PedBE Acceleration | First measurement | - | 47 | 53 |
|  | Covariance over time | - | 78 | 22 |
|  | Second measurement | - | 47 | 53 |
| GrimAge Acceleration | First measurement | 34 | - | 66 |
|  | Covariance over time | 86 | - | 14 |
|  | Second measurement | 30 | - | 70 |
| DunedinPACE | First measurement | 52 | - | 48 |
|  | Covariance over time | 80 | - | 20 |
|  | Second measurement | 47 | - | 53 |

*Note*. The estimates are derived from the best-fitting bivariate twin models across measurement occasions for the four epigenetic aging measures. *A*, relative contribution of the additive genetic factors; *C*, relative contribution of the shared environmental factors; *E*, relative contribution of the unique environmental factors including measurement error.

**Table S16**

*Goodness-of-fit statistics for univariate age-moderated models of epigenetic aging measures* ***unadjusted for biological sex*** *(976 twin pairs)*

| Comparison | ep | -2LL | *df* | AIC | *p* | 1 − β |
| --- | --- | --- | --- | --- | --- | --- |
| Horvath Acceleration | | | | | | |
| Moderated ACE model | 9 | 10 340.14 | 1 943 | 10 358.14 |  | .38 |
| Non-moderated  *a* effect in ACE model | 8 | 10 344.92 | 1 944 | 10 360.92 | .029 | .69 |
| ***Non-moderated c effect in ACE model*** | ***8*** | ***10 340.36*** | ***1 944*** | ***10 356.36*** | ***.639*** |  |
| Non-moderated *e* effect in ACE model | 8 | 10 375.10 | 1 944 | 10 391.10 | < .001 | .99 |
| Moderated AE model | 7 | 10 350.49 | 1 945 | 10 364.49 | .006 | .88 |
| Non-moderated ACE model | 6 | 10 420.84 | 1 946 | 10 432.84 | < .001 | 1 |
| PedBE Acceleration | | | | | | |
| Moderated ACE model | 9 | 5 534.89 | 1 943 | 5 552.89 |  | .63 |
| Non-moderated *a* effect in ACE model | 8 | 5 534.89 | 1 944 | 5 550.89 | .952 | .39 |
| Non-moderated *c* effect in ACE model | 8 | 5 534.96 | 1 944 | 5 550.96 | .794 | .40 |
| Non-moderated *e* effect in ACE model | 8 | 5 629.93 | 1 944 | 5 645.93 | < .001 | 1 |
| ***Non-moderated a and c effect in ACE model*** | ***7*** | ***5 535.00*** | ***1 945*** | ***5 549.00*** | ***.943*** |  |
| Moderated AE model | 7 | 5 562.02 | 1 945 | 5 576.02 | < .001 | 1 |
| Moderated CE model | 7 | 5 538.08 | 1 945 | 5 552.08 | .203 | .54 |
| Non-moderated *c* effect in CE model | 6 | 5 538.30 | 1 946 | 5 550.30 | .332 | .31 |
| Non-moderated ACE model | 6 | 5 650.22 | 1 946 | 5 662.22 | < .001 | 1 |
| GrimAge Acceleration | | | | | | |
| Moderated ACE model | 9 | 8 795.17 | 1 943 | 8 813.17 |  | .73 |
| Non-moderated *a* effect in ACE model | 8 | 8 795.83 | 1 944 | 8 811.83 | .418 | .62 |
| Non-moderated *c* effect in ACE model | 8 | 8 795.18 | 1 944 | 8 811.18 | .950 | .55 |
| Non-moderated *e* effect in ACE model | 8 | 8 795.94 | 1 944 | 8 811.94 | .380 | .63 |
| Moderated CE model | 7 | 8 800.83 | 1 945 | 8 814.83 | .059 | .83 |
| Moderated AE model | 7 | 8 795.25 | 1 945 | 8 809.25 | .959 | .29 |
| Non-moderated *a* effect in AE model | 6 | 8 797.98 | 1 946 | 8 809.98 | .422 | .40 |
| ***Non-moderated e effect in AE model*** | ***6*** | ***8 796.08*** | ***1 946*** | ***8 808.08*** | ***.824*** |  |
| Non-moderated ACE model | 6 | 8 803.18 | 1 946 | 8 815.18 | .046 | .85 |
| Non-moderated AE model | 5 | 8 803.24 | 1 947 | 8 813.24 | .089 | .73 |
| DunedinPACE | | | | | | |
| Moderated ADE model | 9 | -4 550.08 | 1 943 | -4 532.08 |  | .60 |
| Moderated AE model | 7 | -4 547.91 | 1 945 | -4 533.91 | .339 | .38 |
| Non-moderated *a* effect in ADE model | 8 | -4 549.89 | 1 944 | -4 533.89 | .664 | .38 |
| Non-moderated *d* effect in ADE model | 8 | -4 548.36 | 1 944 | -4 532.36 | .190 | .57 |
| Non-moderated *e* effect in ADE model | 8 | -4 549.84 | 1 944 | -4 533.84 | .629 | .39 |
| Non-moderated *a* and *d* effects in ADE model | 7 | -4 542.51 | 1 945 | -4 528.51 | .023 | .85 |
| ***Non-moderated a and e effects in ADE model*** | ***7*** | ***-4 549.69*** | ***1 945*** | ***-4 535.69*** | ***.825*** |  |
| Non-moderated *d* and *e* effects in ADE model | 7 | -4 547.93 | 1 945 | -4 533.93 | .342 | .38 |
| Non-moderated ADE model | 6 | -4 537.25 | 1 946 | -4 525.25 | .005 | .94 |

*Note.* *A/a,* additive genetic factor/effect; *D/d,* non-additive genetic factor/effect; *C/c,* shared environmental factor/effect; *E/e*, individually-unique environmental factor/effect including measurement error; -2LL, log-likelihood statistics; *df*, degrees of freedom; AIC, Akaike information criterion; *p*, results of the comparison of a parsimonious model with a more complex model based on the chi-square difference test; 1 *–* β, achieved statistical power*.* The parsimonious models that did not fit the data worse compared to less restricted models in the sensitivity analysis are shown in italics and those in the main analysis are shown in bold.

**Table S17**

*Standardized variance components in the best-fitting univariate models of age moderation four epigenetic measures* ***unadjusted for biological sex***

| Epigenetic aging measure | Age group | *A* (%) | *C/D* (%) | *E* (%) |
| --- | --- | --- | --- | --- |
| Horvath Accel. | 9.5 | **18** | 35 | **47** |
|  | 12 | **22** | 30 | **48** |
|  | 15.5 | **26** | 25 | **49** |
|  | 18 | **29** | 22 | **49** |
|  | 21.5 | **32** | 19 | **49** |
|  | 24 | **34** | 16 | **49** |
|  | 27.5 | **37** | 14 | **49** |
|  | 30 | **39** | 13 | **49** |
| PedBE Accel. | 9.5 | 18 | 49 | **33** |
|  | 12 | 16 | 45 | **39** |
|  | 15.5 | 14 | 40 | **46** |
|  | 18 | 13 | 36 | **51** |
|  | 21.5 | 12 | 32 | **57** |
|  | 24 | 11 | 29 | **60** |
|  | 27.5 | 9 | 26 | **65** |
|  | 30 | 9 | 24 | **68** |
| GrimAge Accel. | 9.5 | **24** | - | 76 |
|  | 12 | **26** | - | 74 |
|  | 15.5 | **30** | - | 70 |
|  | 18 | **32** | - | 68 |
|  | 21.5 | **35** | - | 65 |
|  | 24 | **37** | - | 63 |
|  | 27.5 | **40** | - | 60 |
|  | 30 | **42** | - | 58 |
| DunedinPACE | 9.5 | 36 | **05** | 59 |
|  | 12 | 35 | **08** | 57 |
|  | 15.5 | 33 | **12** | 54 |
|  | 18 | 32 | **16** | 52 |
|  | 21.5 | 30 | **22** | 48 |
|  | 24 | 28 | **26** | 46 |
|  | 27.5 | 26 | **32** | 42 |
|  | 30 | 24 | **36** | 40 |

*Note.* The estimates are derived from the best-fitting univariate models of age moderation of genetic and environmental components in the four epigenetic aging measures. *A*, relative contribution of the additive genetic factors; *D*, relative contribution of the non-additive genetic factors; *C*, relative contribution of the shared environmental factors; *E*, relative contribution of the unique environmental factors including measurement error; Accel., Acceleration. Components that are moderated by age are shown in bold.

**Table S18**

*Goodness-of-fit statistics for univariate models of epigenetic aging measures* ***unadjusted for smoking probe*** *(976 twin pairs)*

| Comparison | ep | -2LL | *df* | AIC | *p* | 1 − β |
| --- | --- | --- | --- | --- | --- | --- |
| Horvath Acceleration | | | | | | |
| ***ACE model*** | ***4*** | ***10 546.42*** | ***1 948*** | ***10 554.42*** |  |  |
| AE model | 3 | 10 554.62 | 1 949 | 10 560.62 | .004 | .80 |
| CE model | 3 | 10 551.02 | 1 949 | 10 557.02 | .032 | .49 |
| E model | 2 | 10 742.82 | 1 950 | 10 746.82 | < .001 | 1 |
| PedBE Acceleration | | | | | | |
| ACE model | 4 | 5 718.99 | 1 948 | 5 726.99 |  | .41 |
| AE model | 3 | 5 756.65 | 1 949 | 5 762.65 | < .001 | 1 |
| ***CE model*** | ***3*** | ***5 718.99*** | ***1 949*** | ***5 724.99*** | ***1.00*** |  |
| E model | 2 | 5 956.59 | 1 950 | 5 960.59 | < .001 | 1 |
| DunedinPACE | | | | | | |
| ADE model | 4 | -4 349.53 | 1 948 | -4 341.53 |  | .38 |
| ***AE model*** | ***3*** | ***-4 349.28*** | ***1 949*** | ***-4 343.28*** | ***.615*** |  |
| E model | 2 | -4 167.37 | 1 950 | -4 163.37 | < .001 | 1 |

*Note.* *A*, additive genetic factors; *C*, shared environmental factors; *D*, non-additive genetic factors; *E*, unique environmental factors including measurement error; ep, the number of estimated parameters; -2LL, log-likelihood statistics; *df*, degrees of freedom; AIC, Akaike information criterion; *p*, results of the comparison of a parsimonious model with a more complex model based on the chi-square difference test; 1 *–* β, achieved statistical power. The parsimonious models that did not fit the data worse compared to less restricted models in the sensitivity analysis are shown in italics and those in the main analysis are shown in bold.

**Table S19**

*Standardized variance components in the best-fitting univariate models for the three epigenetic measures* ***unadjusted for smoking probe***

| Epigenetic aging measure | *A* (%) | *C* (%) | *E* (%) |
| --- | --- | --- | --- |
| Horvath Accel. | 21 | 26 | 53 |
| PedBE Accel. | - | 46 | 54 |
| DunedinPACE | 50 | - | 50 |

*Note*. The estimates are derived from the best-fitting univariate twin models for the three epigenetic aging measures. *A*, relative contribution of the additive genetic factors; *C*, relative contribution of the shared environmental factors; *E*, relative contribution of the unique environmental factors including measurement error; Accel., Acceleration.

**Table S20**

*Goodness-of-fit statistics for bivariate models of epigenetic aging measures* ***unadjusted for smoking probe*** *(488 twin pairs)*

| Comparison | ep | -2LL | *df* | AIC | *p* | 1 − β |
| --- | --- | --- | --- | --- | --- | --- |
| Horvath Acceleration | | | | | | |
| ACE model | 11 | 10 259.44 | 1 941 | 10 281.44 |  | .41 |
| ACE model with *a*²*1*=*a*²*2*, *c*²*1*=*c*²*2*, and *e*²*1*=*e*²*2* variances | 8 | 10 269.47 | 1 944 | 10 285.47 | .018 | .79 |
| ***ACE model with a*²*1=a*²*2 variances*** | ***10*** | ***10 259.45*** | ***1 942*** | ***10 279.45*** | ***.915*** |  |
| ACE model with *c*²*1*=*c*²*2* variances | 10 | 10 259.81 | 1 942 | 10 279.81 | .544 | .16 |
| ACE model with *e*²*1*=*e*²*2* variances | 10 | 10 260.45 | 1 942 | 10 280.45 | .315 | .26 |
| ACE model with *a*²*1*=*a*²*2* and *c*²*1*=*c*²*2* variances | 9 | 10 263.51 | 1 943 | 10 281.51 | .130 | .42 |
| AE model | 8 | 10 268.14 | 1 944 | 10 284.14 | .034 | .70 |
| CE model | 8 | 10 264.72 | 1 944 | 10 280.72 | .152 | .31 |
| E model | 5 | 10 426.59 | 1 947 | 10 436.59 | < .001 | 1 |
| PedBE Acceleration | | | | | | |
| ACE model | 11 | 5 424.32 | 1 941 | 5 446.32 |  | .34 |
| AE model | 8 | 5 452.02 | 1 944 | 5 468.02 | < .001 | 1 |
| ***CE model*** | ***8*** | ***5 428.83*** | ***1 944*** | ***5 444.83*** | ***.212*** |  |
| CE model with *e*²*1*=*e*²*2* variances | 7 | 5 438.37 | 1 945 | 5 452.37 | .002 | .86 |
| CE model with *c*²*1*=*c*²*2* variances | 7 | 5 433.70 | 1 945 | 5 447.70 | .027 | .52 |
| E model | 5 | 5 630.53 | 1 947 | 5 640.53 | < .001 | 1 |
| DunedinPACE | | | | | | |
| ADE model | 11 | -4 561.77 | 1 941 | -4 539.77 |  | .86 |
| AE model | *8* | -4 561.40 | 1 944 | -4 545.40 | .947 | .39 |
| ***AE model with a*²*1=a*²*2 variances*** | ***7*** | ***-4 561.23*** | ***1 945*** | ***-4 547.23*** | ***.683*** |  |
| AE model with *e*²*1*=*e*²*2* variances | **7** | -4 557.46 | 1 945 | -4 543.46 | .047 | .62 |
| E model | 5 | -4 229.32 | 1 947 | -4 219.32 | < .001 | 1 |

*Note.* *A/a*, additive genetic factor/factor’s variances; *D/d*, non-additive genetic factor/factor’s variances; *C/c*, shared environmental factor/factor’s variances; *E/e*, individually-unique environmental factor/factor’s variances including measurement error; ep, the number of estimated parameters; *p*, results of the comparison of a parsimonious model with a more complex model based on the chi-square difference test; 1 *–* β, achieved statistical power*.* The parsimonious models that did not fit the data worse compared to less restricted models in the sensitivity analysis are shown in italics and those in the main analysis are shown in bold.

**Table S21**

*Standardized variance components in the best-fitting bivariate models for the three epigenetic measures* ***unadjusted for smoking probe***

| Epigenetic aging measure | Measurement | *A* (%) | *C* (%) | *E* (%) |
| --- | --- | --- | --- | --- |
| Horvath Acceleration | First measurement | 25 | 21 | 54 |
|  | Covariance over time | 26 | 52 | 22 |
|  | Second measurement | 21 | 28 | 51 |
| PedBE Acceleration | First measurement | - | 47 | 53 |
|  | Covariance over time | - | 78 | 22 |
|  | Second measurement | - | 46 | 54 |
| DunedinPACE | First measurement | 52 | - | 48 |
|  | Covariance over time | 83 | - | 17 |
|  | Second measurement | 48 | - | 52 |

*Note*. The estimates are derived from the best-fitting bivariate twin models across measurement occasions for the three epigenetic aging measures. *A*, relative contribution of the additive genetic factors; *C*, relative contribution of the shared environmental factors; *E*, relative contribution of the unique environmental factors including measurement error.

**Table S22**

*Goodness-of-fit statistics for univariate age-moderated models of epigenetic aging measures* ***unadjusted for smoking probe*** *(976 twin pairs)*

| Comparison | ep | -2LL | *df* | AIC | *p* | 1 − β |
| --- | --- | --- | --- | --- | --- | --- |
| Horvath Acceleration | | | | | | |
| Moderated ACE model | 9 | 10 454.70 | 1 943 | 10 472.70 |  | .39 |
| Non-moderated *a* effect in ACE model | 8 | 10 458.68 | 1 944 | 10 474.68 | .046 | .62 |
| ***Non-moderated c effect in ACE model*** | ***8*** | ***10 454.86*** | ***1 944*** | ***10 470.86*** | ***.688*** |  |
| Non-moderated *e* effect in ACE model | 8 | 10 495.52 | 1 944 | 10 511.52 | < .001 | 1 |
| Moderated AE model | 7 | 10 464.49 | 1 945 | 10 478.49 | .007 | .87 |
| Non-moderated ACE model | 6 | 10 539.59 | 1 946 | 10 551.59 | < .001 | 1 |
| PedBE Acceleration | | | | | | |
| Moderated ACE model | 9 | 5 562.30 | 1 943 | 5 580.30 |  | .63 |
| Non-moderated *a* effect in ACE model | 8 | 5 562.34 | 1 944 | 5 578.34 | .842 | .41 |
| Non-moderated *c* effect in ACE model | 8 | 5 562.32 | 1 944 | 5 578.32 | .887 | .41 |
| Non-moderated *e* effect in ACE model | 8 | 5 672.59 | 1 944 | 5 688.59 | < .001 | 1 |
| ***Non-moderated a and c effects in ACE model*** | ***7*** | ***5 562.34*** | ***1 945*** | ***5 576.34*** | ***.980*** |  |
| Moderated AE model | 7 | 5 590.28 | 1 945 | 5 604.28 | < .001 | 1 |
| Moderated CE model | 7 | 5 566.23 | 1 945 | 5 580.23 | .140 | .63 |
| Non-moderated *c* effect in CE model | 6 | 5 566.25 | 1 946 | 5 578.25 | .267 | .38 |
| Non-moderated CE model | 5 | 5 690.16 | 1 947 | 5 700.16 | < .001 | 1 |
| DunedinPACE | | | | | | |
| Moderated ADE model | 9 | -4 387.17 | 1 943 | -4 369.17 |  | .54 |
| Moderated AE model | 7 | -4 385.06 | 1 945 | -4 371.06 | .348 | .29 |
| Non-moderated *a* effect in ADE model | 8 | -4 386.50 | 1 944 | -4 370.50 | .413 | .37 |
| Non-moderated *d* effect in ADE model | 8 | -4 385.28 | 1 944 | -4 369.28 | .168 | .53 |
| Non-moderated *e* effect in ADE model | 8 | -4 386.79 | 1 944 | -4 370.79 | .534 | .33 |
| Non-moderated *a* and *d* effects in ADE model | 7 | -4 383.10 | 1 945 | -4 369.10 | .131 | .55 |
| ***Non-moderated a and e effects in ADE model*** | ***7*** | ***-4 386.21*** | ***1 945*** | ***-4 372.21*** | ***.616*** |  |
| Non-moderated *d* and *e* effects in ADE model | 7 | -4 384.69 | 1 945 | -4 370.69 | .288 | .34 |
| Non-moderated ADE model | 6 | -4 380.07 | 1 946 | -4 368.07 | .069 | .65 |

*Note.* *A/a,* additive genetic factor/effect; *D/d,* non-additive genetic factor/effect; *C/c,* shared environmental factor/effect; *E/e*, individually-unique environmental factor/effect including measurement error; ep, the number of estimated parameters; -2LL, log-likelihood statistics; *df*, degrees of freedom; AIC, Akaike information criterion; *p*, results of the comparison of a parsimonious model with a more complex model based on the chi-square difference test; 1 *–* β, achieved statistical power*.* The parsimonious models that did not fit the data worse compared to less restricted models in the sensitivity analysis are shown in italics and those in the main analysis are shown in bold.

**Table S23**

*Standardized variance components in the best-fitting univariate models of age moderation three epigenetic measures* ***unadjusted for smoking probe***

| Epigenetic aging measure | Age group | *A* (%) | *C/D* (%) | *E* (%) |
| --- | --- | --- | --- | --- |
| Horvath Accel. | 9.5 | **17** | 35 | **49** |
|  | 12 | **20** | 30 | **50** |
|  | 15.5 | **24** | 25 | **52** |
|  | 18 | **26** | 22 | **52** |
|  | 21.5 | **29** | 18 | **53** |
|  | 24 | **30** | 16 | **53** |
|  | 27.5 | **33** | 14 | **54** |
|  | 30 | **34** | 12 | **54** |
| PedBE Accel. | 9.5 | 19 | 49 | **31** |
|  | 12 | 17 | 45 | **37** |
|  | 15.5 | 15 | 40 | **45** |
|  | 18 | 14 | 36 | **50** |
|  | 21.5 | 12 | 31 | **56** |
|  | 24 | 11 | 29 | **60** |
|  | 27.5 | 10 | 25 | **65** |
|  | 30 | 9 | 23 | **68** |
| DunedinPACE | 9.5 | 42 | **2** | 56 |
|  | 12 | 42 | **4** | 55 |
|  | 15.5 | 40 | **7** | 53 |
|  | 18 | 39 | **10** | 51 |
|  | 21.5 | 37 | **14** | 49 |
|  | 24 | 36 | **17** | 47 |
|  | 27.5 | 34 | **22** | 44 |
|  | 30 | 32 | **26** | 42 |

*Note.* The estimates are derived from the best-fitting univariate models of age moderation of genetic and environmental components of the variance in the three epigenetic aging measures. *A*, relative contribution of the additive genetic factors; *D*, relative contribution of the non-additive genetic factors; *C*, relative contribution of the shared environmental factors; *E*, relative contribution of the unique environmental factors including measurement error; Accel., Acceleration. Components that are moderated by age are shown in bold.

**Table S24**

*Goodness-of-fit statistics for univariate models of epigenetic aging measures* ***unadjusted for cell type composition*** *(976 twin pairs)*

| Comparison | ep | -2LL | *df* | AIC | *p* | 1 − β |
| --- | --- | --- | --- | --- | --- | --- |
| Horvath Acceleration | | | | | | |
| **ACE model** | **4** | **10 455.53** | **1 948** | **10 463.53** |  | .41 |
| *AE model* | *3* | *10 455.53* | *1 949* | *10 461.53* | *1.00* |  |
| CE model | 3 | 10 455.66 | 1 949 | 10 461.66 | .717 | .12 |
| E model | 2 | 10 669.87 | 1 950 | 10 673.87 | < .001 | 1 |
| PedBE Acceleration | | | | | | |
| ACE model | 4 | 5 978.15 | 1 948 | 5 986.15 |  | .41 |
| AE model | 3 | 5 999.49 | 1 949 | 6 005.49 | < .001 | 1 |
| ***CE model*** | ***3*** | ***5 978.15*** | ***1 949*** | ***5 984.15*** | ***1.00*** |  |
| E model | 2 | 6 166.56 | 1 950 | 6 170.56 | < .001 | 1 |
| GrimAge Acceleration | | | | | | |
| ACE model | 4 | 10 322.96 | 1 948 | 10 330.96 |  | .58 |
| **AE model** | **3** | **10 322.96** | **1 949** | **10 328.96** | **1.00** | .33 |
| *CE model* | *3* | *10 321.52* | *1 949* | *10 327.52* | *1.00* |  |
| E model | 2 | 10 345.19 | 1 950 | 10 349.19 | < .001 | 1 |
| DunedinPACE | | | | | | |
| *ADE model* | *4* | *-3 771.59* | *1 948* | *-3 763.59* |  |  |
| **AE model** | **3** | **-3 767.21** | **1 949** | **-3 761.21** | **.036** | .46 |
| E model | 2 | -3 700.45 | 1 950 | -3 696.45 | < .001 | 1 |

*Note.* *A*, additive genetic factors; *C*, shared environmental factors; *D*, non-additive genetic factors; *E*, unique environmental factors including measurement error; ep, the number of estimated parameters; -2LL, log-likelihood statistics; *df*, degrees of freedom; AIC, Akaike information criterion; *p*, results of the comparison of a parsimonious model with a more complex model based on the chi-square difference test; 1 *–* β, achieved statistical power. The parsimonious models that did not fit the data worse compared to less restricted models in the sensitivity analysis are shown in italics and those in the main analysis are shown in bold.

**Table S25**

*Standardized variance components in the best-fitting univariate models for the four epigenetic measures* ***unadjusted for cell type composition***

| Epigenetic aging measure | *A* (%) | *C/D* (%) | *E* (%) |
| --- | --- | --- | --- |
| Horvath Accel. | 52 | - | 48 |
| PedBE Accel. | - | 42 | 58 |
| GrimAge Accel. | - | 15 | 85 |
| DunedinPACE | 0 | 37 | 63 |

*Note*. The estimates are derived from the best-fitting univariate twin models for the four epigenetic aging measures. *A*, relative contribution of the additive genetic factors; *D*, relative contribution of the non-additive genetic factors; *C*, relative contribution of the shared environmental factors; *E*, relative contribution of the unique environmental factors including measurement error; Accel., Acceleration.

**Table S26**

*Goodness-of-fit statistics for bivariate models of epigenetic aging measures* ***unadjusted for cell type composition*** *(488 twin pairs)*

| Comparison | ep | -2LL | *df* | AIC | *p* | 1 − β |
| --- | --- | --- | --- | --- | --- | --- |
| Horvath Acceleration | | | | | | |
| ACE model | 11 | 10 150.28 | 1 941 | 10 172.28 |  | .41 |
| ACE model with *a*²*1*=*a*²*2*, *c*²*1*=*c*²*2*, and *e*²*1*=*e*²*2* variances | 8 | 10 166.32 | 1 944 | 10 182.32 | .001 | .97 |
| ***ACE model with a*²*1=a*²*2 variances*** | ***10*** | ***10 150.30*** | ***1 942*** | ***10 170.30*** | ***.899*** |  |
| ACE model with *c*²*1*=*c*²*2* variances | 10 | 10 150.91 | 1 942 | 10 170.91 | .426 | .20 |
| ACE model with *e*²*1*=*e*²*2* variances | 10 | 10 154.47 | 1 942 | 10 174.47 | .041 | .65 |
| ACE model with *a*²*1*=*a*²*2* and *c*²*1*=*c*²*2* variances | 9 | 10 181.01 | 1 943 | 10 199.01 | < .001 | 1 |
| AE model | 8 | 10 156.82 | 1 944 | 10 172.82 | .088 | .48 |
| CE model | 8 | 10 160.32 | 1 944 | 10 176.32 | .018 | .79 |
| E model | 5 | 10 343.21 | 1 947 | 10 353.21 | < .001 | 1 |
| PedBE Acceleration | | | | | | |
| ACE model | 11 | 5 698.01 | 1 941 | 5 720.01 |  | .60 |
| AE model | 8 | 5 712.90 | 1 944 | 5 728.90 | .002 | .97 |
| ***CE model*** | ***8*** | ***5 700.43*** | ***1 944*** | ***5 716.43*** | ***.490*** |  |
| CE model with *e*²*1*=*e*²*2* variances | 7 | 5 711.74 | 1 945 | 5 725.74 | .001 | .92 |
| CE model with *c*²*1*=*c*²*2* variances | 7 | 5 704.70 | 1 945 | 5 718.70 | .039 | .45 |
| E model | 5 | 5 858.56 | 1 947 | 5 868.56 | < .001 | 1 |
| GrimAge Acceleration | | | | | | |
| ACE model | 11 | 10 254.12 | 1 941 | 10 276.12 |  | .86 |
| AE model | 8 | 10 255.85 | 1 944 | 10 271.85 | .632 | .55 |
| CE model | 8 | 10 254.70 | 1 944 | 10 270.70 | .901 | .41 |
| *CE model with c*²*1=c*²*2 variances* | *7* | *10 254.72* | *1 945* | *10 268.72* | *1.00* |  |
| CE model with *e*²*1*=*e*²*2* variances | 7 | 10 283.23 | 1 945 | 10 297.23 | < .001 | 1 |
| E model | 5 | 10 281.56 | 1 947 | 10 291.56 | < .001 | 1 |
| DunedinPACE | | | | | | |
| ADE model | 11 | -3 919.63 | 1 941 | -3 897.63 |  | .62 |
| ADE model with *a*²*1=a*²*2*, *c*²*1=c*²*2*, and *e*²*1=e*²*2* variances | *8* | -3 901.10 | 1 944 | -3 885.10 | < .001 | .99 |
| ADE model with *a*²*1=a*²*2* variances | 10 | -3 919.50 | 1 942 | -3 899.50 | .724 | .40 |
| ADE model with *c*²*1=c*²*2* variances | 10 | -3 919.47 | 1 942 | -3 899.47 | .687 | .40 |
| ADE model with *e*²*1=e*²*2* variances | 10 | -3 911.30 | 1 942 | -3 891.30 | .004 | .93 |
| *ADE model with a*²*1=a*²*2 and c*²*1=c*²*2 effects* | *9* | *-3 919.43* | *1 943* | *-3 901.43* | *.907* |  |
| AE model | 7 | -3 912.84 | 1 944 | -3 896.84 | .079 | .69 |
| E model | 5 | -3 847.99 | 1 947 | -3 837.99 | < .001 | 1 |

*Note.* *A/a*, additive genetic factor/factor’s variances; *D/d*, non-additive genetic factor/factor’s variances; *C/c*, shared environmental factor/factor’s variances; *E/e*, individually-unique environmental factor/factor’s variances including measurement error; ep, the number of estimated parameters; *p*, results of the comparison of a parsimonious model with a more complex model based on the chi-square difference test; -2LL, log-likelihood statistics; *df*, degrees of freedom; AIC, Akaike information criterion; *p*, results of the comparison of a parsimonious model with a more complex model based on the chi-square difference test; 1 *–* β, achieved statistical power. The parsimonious models that did not fit the data worse compared to less restricted models in the sensitivity analysis are shown in italics and those in the main analysis are shown in bold.

**Table S27**

*Standardized variance components in the best-fitting bivariate models for the four epigenetic measures* ***unadjusted for cell type composition***

| Epigenetic aging measure | Measurement | *A* (%) | *C/D* (%) | *E* (%) |
| --- | --- | --- | --- | --- |
| Horvath Acceleration | First measurement | 31 | 20 | 49 |
|  | Covariance over time | 48 | 39 | 13 |
|  | Second measurement | 25 | 25 | 50 |
| PedBE Acceleration | First measurement | - | 42 | 58 |
|  | Covariance over time | - | 70 | 30 |
|  | Second measurement | - | 42 | 58 |
| GrimAge Acceleration | First measurement | - | 19 | 81 |
|  | Covariance over time | - | 11 | 89 |
|  | Second measurement | - | 13 | 87 |
| DunedinPACE | First measurement | -1 | 45 | 56 |
|  | Covariance over time | -50 | 122 | 28 |
|  | Second measurement | -1 | 36 | 65 |

*Note*. The estimates are derived from the best-fitting bivariate twin models across measurement occasions for the four epigenetic aging measures. *A*, relative contribution of the additive genetic factors; *D*, relative contribution of the non-additive genetic factors; *C*, relative contribution of the shared environmental factors; *E*, relative contribution of the unique environmental factors including measurement error.

**Table S28**

*Goodness-of-fit statistics for univariate age-moderated models of epigenetic aging measures* ***unadjusted for cell type composition*** *(976 twin pairs)*

| Comparison | ep | -2LL | *df* | AIC | *p* | 1 − β |
| --- | --- | --- | --- | --- | --- | --- |
| Horvath Acceleration | | | | | | |
| Moderated ACE model | 9 | 10 362.46 | 1 943 | 10 380.46 |  | .38 |
| Non-moderated *a* effect in ACE model | 8 | 10 367.98 | 1 944 | 10 383.98 | .019 | .75 |
| ***Non-moderated c effect in ACE model*** | ***8*** | ***10 362.67*** | ***1 944*** | ***10 378.67*** | ***.640*** |  |
| Non-moderated *e* effect in ACE model | 8 | 10 394.81 | 1 944 | 10 410.81 | < .001 | 1 |
| Moderated AE model | 7 | 10 371.23 | 1 945 | 10 385.23 | .012 | .82 |
| Non-moderated ACE model | 6 | 10 445.72 | 1 946 | 10 457.72 | < .001 | 1 |
| PedBE Acceleration | | | | | | |
| Moderated ACE model | 9 | 5 759.41 | 1 943 | 5 777.41 |  | .39 |
| Non-moderated *a* effect in ACE model | 8 | 5 760.29 | 1 944 | 5 776.29 | .347 | .22 |
| *Non-moderated c effect in ACE model* | *8* | *5 759.56* | *1 944* | *5 775.56* | *.697* |  |
| Non-moderated *e* effect in ACE model | 8 | 5 876.93 | 1 944 | 5 892.93 | < .001 | 1 |
| **Non-moderated *a* and *c* effects in ACE model** | **7** | **5 763.69** | **1 945** | **5 777.69** | **.117** | **.43** |
| Moderated AE model | 7 | 5 777.34 | 1 945 | 5 791.34 | < .001 | .99 |
| Moderated CE model | 7 | 5 762.62 | 1 945 | 5 776.62 | .201 | .27 |
| Non-moderated *c* effect in CE model | 6 | 5 766.26 | 1 946 | 5 778.26 | .077 | .50 |
| Non-moderated ACE model | 6 | 5 932.48 | 1 946 | 5 944.48 | < .001 | 1 |
| GrimAge Acceleration | | | | | | |
| Moderated ACE model | 9 | 10 306.86 | 1 943 | 10 324.86 |  | .65 |
| Non-moderated *a* effect in ACE model | 8 | 10 306.89 | 1 944 | 10 322.89 | .870 | .44 |
| Non-moderated *c* effect in ACE model | 8 | 10 308.00 | 1 944 | 10 324.00 | .287 | .57 |
| Non-moderated *e* effect in ACE model | 8 | 10 320.43 | 1 944 | 10 336.43 | < .001 | .99 |
| Moderated CE model | 7 | 10 306.97 | 1 945 | 10 320.97 | .948 | .15 |
| Moderated AE model | 7 | 10 309.25 | 1 945 | 10 323.25 | .303 | .48 |
| **Non-moderated *e* effect in AE model** | **6** | **10 310.32** | **1 946** | **10 322.32** | **.326** | .36 |
| *Non-moderated c effect in CE model* | *6* | *10 308.70* | *1 946* | *10 320.70* | *.608* |  |
| Non-moderated *e* effect in CE model | 6 | 10 321.06 | 1 946 | 10 333.06 | .003 | .97 |
| Non-moderated ACE model | 6 | 10 321.11 | 1 946 | 10 333.11 | .003 | .97 |
| Non-moderated AE model | 5 | 10 322.71 | 1 947 | 10 332.71 | .003 | .97 |
| DunedinPACE | | | | | | |
| Moderated ADE model | 9 | -3 788.62 | 1 943 | -3 770.62 |  | .44 |
| Moderated AE model | 7 | -3 784.33 | 1 945 | -3 770.33 | .117 | .47 |
| Non-moderated *a* effect in ADE model | 8 | -3 788.57 | 1 944 | -3 772.57 | .825 | .14 |
| Non-moderated *d* effect in ADE model | 8 | -3 787.64 | 1 944 | -3 771.64 | .324 | .29 |
| Non-moderated *e* effect in ADE model | 8 | -3 784.71 | 1 944 | -3 768.71 | .048 | .65 |
| *Non-moderated a and d effects in ADE model* | *7* | *-3 786.81* | *1 945* | *-3 772.81* | *.406* |  |
| **Non-moderated *a* and *e* effects in ADE model** | **7** | **-3 784.68** | **1 945** | **-3 770.68** | **.140** | **.43** |
| Non-moderated *d* and *e* effects in ADE model | 7 | -3 781.75 | 1 945 | -3 767.75 | .032 | .73 |
| Non-moderated ADE model | 6 | -3 773.34 | 1 946 | -3 761.34 | .002 | .96 |

*Note.* *A/a,* additive genetic factor/effect; *D/d,* non-additive genetic factor/effect; *C/c,* shared environmental factor/effect; *E/e*, individually-unique environmental factor/effect including measurement error; -2LL, log-likelihood statistics; *df*, degrees of freedom; AIC, Akaike information criterion; *p*, results of the comparison of a parsimonious model with a more complex model based on the chi-square difference test; 1 *–* β, achieved statistical power*.* The parsimonious models that did not fit the data worse compared to less restricted models in the sensitivity analysis are shown in italics and those in the main analysis are shown in bold.

**Table S29**

*Standardized variance components in the best-fitting univariate models of age moderation four epigenetic measures* ***unadjusted for cell type composition***

| Epigenetic aging measure | Age group | *A* (%) | *C/D* (%) | *E* (%) |
| --- | --- | --- | --- | --- |
| Horvath Accel. | 9.5 | **19** | 32 | **49** |
|  | 12 | **23** | 28 | **49** |
|  | 15.5 | **28** | 23 | **49** |
|  | 18 | **31** | 20 | **49** |
|  | 21.5 | **35** | 17 | **49** |
|  | 24 | **37** | 15 | **48** |
|  | 27.5 | **40** | 13 | **48** |
|  | 30 | **41** | 11 | **47** |
| PedBE Accel. | 9.5 | **14** | 45 | **40** |
|  | 12 | **16** | 39 | **45** |
|  | 15.5 | **18** | 31 | **51** |
|  | 18 | **19** | 27 | **54** |
|  | 21.5 | **20** | 22 | **58** |
|  | 24 | **21** | 20 | **60** |
|  | 27.5 | **21** | 17 | **62** |
|  | 30 | **22** | 15 | **63** |
| GrimAge Accel. | 9.5 | - | 19 | **81** |
|  | 12 | - | 18 | **82** |
|  | 15.5 | - | 17 | **83** |
|  | 18 | - | 16 | **84** |
|  | 21.5 | - | 15 | **85** |
|  | 24 | - | 15 | **85** |
|  | 27.5 | - | 14 | **86** |
|  | 30 | - | 13 | **87** |
| DunedinPACE | 9.5 | 3 | 38 | **58** |
|  | 12 | 3 | 37 | **60** |
|  | 15.5 | 3 | 35 | **63** |
|  | 18 | 3 | 33 | **64** |
|  | 21.5 | 3 | 31 | **66** |
|  | 24 | 2 | 30 | **68** |
|  | 27.5 | 2 | 28 | **69** |
|  | 30 | 2 | 27 | **71** |

*Note.* The estimates are derived from the best-fitting univariate models of age moderation of genetic and environmental components in the four epigenetic aging measures. *A*, relative contribution of the additive genetic factors; *D*, relative contribution of the non-additive genetic factors; *C*, relative contribution of the shared environmental factors; *E*, relative contribution of the unique environmental factors including measurement error; Accel., Acceleration. Components that are moderated by age are shown in bold.

**Table S30**

*Goodness-of-fit statistics for univariate twin models with age as a moderator for epigenetic measures of the first measurement occasion only (488 twin pairs)*

| Comparison | ep | -2LL | *df* | AIC | *p* | 1 − β |
| --- | --- | --- | --- | --- | --- | --- |
| Horvath Acceleration | | | | | | |
| Moderated ACE model | 9 | 5 053.52 | 967 | 5 071.52 |  | .35 |
| Non-moderated *a* effect in ACE model | 8 | 5 056.18 | 968 | 5 072.18 | .103 | .44 |
| ***Non-moderated c effect in ACE model*** | ***8*** | ***5 053.95*** | ***968*** | ***5 069.95*** | ***.510*** |  |
| Non-moderated *e* effect in ACE model | 8 | 5 071.29 | 968 | 5 087.29 | < .001 | .99 |
| Moderated AE model | 7 | 5 058.74 | 969 | 5 072.74 | .073 | .51 |
| Non-moderated ACE model | 7 | 5 055.98 | 969 | 5 069.98 | .292 | .11 |
| PedBE Acceleration | | | | | | |
| Moderated ACE model | 9 | 2 637.61 | 967 | 2 655.61 |  | .50 |
| Non-moderated *a* effect in ACE model | 8 | 2 639.05 | 968 | 2 655.05 | .229 | .43 |
| Non-moderated *c* effect in ACE model | 8 | 2 638.55 | 968 | 2 654.55 | .332 | .36 |
| Non-moderated *e* effect in ACE model | 8 | 2 676.58 | 968 | 2 692.58 | < .001 | 1 |
| **Non-moderated *a* and *c* effects in ACE model** | **7** | **2 639.06** | **969** | **2 653.06** | **.483** | **.12** |
| Moderated AE model | 7 | 2 651.58 | 969 | 2 665.58 | .001 | .97 |
| Moderated CE model | 7 | 2 640.90 | 969 | 2 654.90 | .192 | .41 |
| *Non-moderated c effect in CE model* | *6* | *2 640.93* | *970* | *2 652.93* | *.345* |  |
| Non-moderated ACE model | 6 | 2 689.97 | 970 | 2 701.97 | < .001 | 1 |
| GrimAge Acceleration | | | | | | |
| Moderated ACE model | 9 | 4 304.65 | 967 | 4 322.65 |  | .71 |
| Non-moderated *a* effect in ACE model | 8 | 4 305.97 | 968 | 4 321.97 | .251 | .66 |
| Non-moderated *c* effect in ACE model | 8 | 4 304.65 | 968 | 4 320.65 | 1.00 | .52 |
| Non-moderated *e* effect in ACE model | 8 | 4 305.77 | 968 | 4 321.77 | .291 | .64 |
| Moderated CE model | 7 | 4 309.96 | 969 | 4 323.96 | .070 | .80 |
| Moderated AE model | 7 | 4 304.65 | 969 | 4 318.65 | 1.00 | .24 |
| Non-moderated *a* effect in AE model | 6 | 4 307.68 | 970 | 4 319.68 | .387 | .40 |
| ***Non-moderated e*** ***effect in AE model*** | ***6*** | ***4 305.77*** | ***970*** | ***4 317.77*** | ***.774*** |  |
| Non-moderated ACE model | 6 | 4 313.75 | 970 | 4 325.75 | .028 | .88 |
| Non-moderated AE model | 5 | 4 313.75 | 971 | 4 323.75 | .059 | .79 |
| DunedinPACE | | | | | | |
| Moderated ADE model | 9.00 | -2 343.16 | 967 | -2 325.16 |  | .63 |
| Moderated AE model | 7 | -2 341.88 | 969 | -2 327.88 | .527 | .29 |
| Non-moderated *a* effect in ADE model | 8 | -2 343.04 | 968 | -2 327.04 | .727 | .41 |
| Non-moderated *d* effect in ADE model | 8 | -2 342.18 | 968 | -2 326.18 | .321 | .52 |
| Non-moderated *e* effect in ADE model | 8 | -2 343.15 | 968 | -2 327.15 | .919 | .39 |
| Non-moderated *a* and *d* effects in ADE model | 7 | -2 339.54 | 969 | -2 325.54 | .163 | .59 |
| ***Non-moderated a and e effects in ADE model*** | ***7*** | ***-2 343.04*** | ***969*** | ***-2 329.04*** | ***.938*** |  |
| Non-moderated *d* and *e* effects in ADE model | 7 | -2 342.12 | 969 | -2 328.12 | .593 | .25 |
| Non-moderated ADE model | 6 | -2 337.92 | 970 | -2 325.92 | .155 | .55 |

*Note.* Goodness-of-fit statistics are presented for univariate twin models with age-moderated effects, quadratic, and linear main effects of age. *A/a,* additive genetic factor/effect; *D/d,* non-additive genetic factor/effect; *C/c,* shared environmental factor/effect; *E/e*, individually-unique environmental factor/effect including measurement error; ep, the number of estimated parameters; -2LL, log-likelihood statistics; *df*, degrees of freedom; AIC, Akaike information criterion; *p*, results of the comparison of a parsimonious model with a more complex model based on the chi-square difference test; 1 *–* β, achieved statistical power*.* The parsimonious models that did not fit the data worse compared to less restricted models in the sensitivity analysis are shown in italics and those in the main analysis are shown in bold.
