## Supplementary material for "Genetic and Environmental Contributions to Epigenetic Aging Across Adolescence and Young Adulthood": Tables within the manuscript

**Table 1**

*Descriptive statistics for epigenetic aging measures*

|  | Early adolescence | | Late adolescence | | Emerging adulthood | | Young adulthood | |
| --- | --- | --- | --- | --- | --- | --- | --- | --- |
| Variable | Time 1:  age 9.5 | Time 2:  age 12 | Time 1:  age 15.5 | Time 2:  age 18 | Time 1:  age 21.5 | Time 2:  age 24 | Time 1:  age 27.5 | Time 2:  age 30 |
| *N* (MZ pairs/DZ pairs) | 326 (82/81) | | 372 (87/99) | | 152 (42/34) | | 126 (52/11) | |
| Chronological Age *M* (*SD*) | 9.51 (0.33) | 11.83 (0.58) | 15.48 (0.31) | 17.75 (0.58) | 21.58 (0.37) | 23.89 (0.62) | 27.60 (0.70) | 29.92 (1.04) |
| Horvath Accel. *M* (*SD*) | -0.02 (2.64) | -0.25 (3.35) | -0.07 (3.56) | 0.28 (3.99) | 0.47 (4.30) | 0.29 (4.38) | -0.30 (4.17) | -0.52 (4.20) |
| PedBE Accel. *M* (*SD*) | -0.10 (0.91) | -0.21 (0.98) | 0.10 (0.95) | 0.24 (1.14) | 0.16 (1.21) | 0.17 (1.36) | -0.22 (1.24) | -0.39 (1.30) |
| GrimAge Accel. *M* (*SD*) | 0.31 (2.03) | -0.08 (2.41) | -0.44 (2.32) | 0.11 (2.40) | 0.09 (2.23) | 0.01 (2.35) | 0.37 (2.59) | -0.14 (2.61) |
| DunedinPACE *M* (*SD*) | 0.00 (0.07) | -0.01 (0.08) | -0.01 (0.08) | 0.00 (0.08) | 0.00 (0.08) | 0.00 (0.08) | 0.02 (0.08) | 0.02 (0.09) |

*Note.* Four cohorts (early adolescents, late adolescents, emerging adults, and young adults) were measured twice, approximately 2.5 years apart. MZ, monozygotic; DZ, dizygotic; *M*, mean; *SD*, standard deviation; Accel., Acceleration.

**Table 2**

*Spearman correlations for monozygotic and dizygotic twins for epigenetic aging measures*

|  |  |  | Early adolescence | | Late adolescence | | Emerging adulthood | | Young adulthood | |
| --- | --- | --- | --- | --- | --- | --- | --- | --- | --- | --- |
| Variable | Zygosity | Cross-cohort correlation | Time 1: age 9.5 | Time 2: age 12 | Time 1: age 15.5 | Time 2:  age 18 | Time 1: age 21.5 | Time 2: age 24 | Time 1: age 27.5 | Time 2:  age 30 |
| Horvath Accel. | MZ | .51*** | .44*** | .53*** | .46*** | .49*** | .56*** | .69*** | .59*** | .43** |
|  | DZ | .37*** | .37*** | .59*** | .34*** | .40*** | .25 | .08 | .52 | .36 |
| PedBE Accel. | MZ | .45*** | .51*** | .61*** | .61*** | .48*** | .76*** | .59*** | .11 | .06 |
|  | DZ | .49*** | .60*** | .45*** | .55*** | .61*** | .01 | .26 | .23 | .33 |
| GrimAge Accel. | MZ | .32*** | .27* | .30** | .35*** | .16 | .43** | .34* | .45*** | .39** |
|  | DZ | .17*** | .17 | .10 | .11 | .31** | .23 | .04 | .12 | .36 |
| DunedinPACE | MZ | .48*** | .44*** | .50*** | .40*** | .36*** | .65*** | .62*** | .53*** | .48*** |
|  | DZ | .19*** | .32** | .23* | .16 | .13 | .22 | .23 | .26 | -.13 |

*Note.* MZ, monozygotic; DZ, dizygotic; Accel., Acceleration; *, *p* < .05; **, *p* < .01; ***, *p* < .001.

**Table 3**

*Variance components derived from age-moderated univariate models*

| Epigenetic aging measure | Age group | *A* (%) | *C/D* (%) | *E* (%) | *V* |
| --- | --- | --- | --- | --- | --- |
| Horvath Accel.  (ACE model with moderated *a* and *e* effects) | 9.5 | **1.59 (18)** | 3.04 (35) | **4.18 (47)** | 8.82 |
|  | 12 | **2.20 (22)** | 3.04 (30) | **4.90 (48)** | 10.14 |
|  | 15.5 | **3.20 (26)** | 3.04 (25) | **6.00 (49)** | 12.24 |
|  | 18 | **4.03 (29)** | 3.04 (22) | **6.85 (49)** | 13.92 |
|  | 21.5 | **5.36 (32)** | 3.04 (19) | **8.14 (49)** | 16.54 |
|  | 24 | **6.42 (35)** | 3.04 (16) | **9.13 (49)** | 18.59 |
|  | 27.5 | **8.08 (37)** | 3.04 (14) | **10.60 (49)** | 21.72 |
|  | 30 | **9.37 (39)** | 3.04 (12) | **11.73 (49)** | 24.14 |
| PedBE Accel.  (ACE model with moderated *e* effect) | 9.5 | 0.16 (18) | 0.43 (49) | **0.29 (33)** | 0.88 |
|  | 12 | 0.16 (16) | 0.43 (45) | **0.37 (39)** | 0.96 |
|  | 15.5 | 0.16 (14) | 0.43 (40) | **0.50 (46)** | 1.09 |
|  | 18 | 0.16 (13) | 0.43 (36) | **0.60 (51)** | 1.19 |
|  | 21.5 | 0.16 (11) | 0.43 (32) | **0.76 (57)** | 1.35 |
|  | 24 | 0.16 (11) | 0.43 (29) | **0.89 (60)** | 1.48 |
|  | 27.5 | 0.16 (9) | 0.43 (26) | **1.08 (65)** | 1.67 |
|  | 30 | 0.16 (8) | 0.43 (24) | **1.23 (68)** | 1.82 |
| GrimAge Accel.  (AE model with moderated *a* effect) | 9.5 | **1.17 (24)** | - | 3.78 (76) | 4.95 |
|  | 12 | **1.33 (26)** | - | 3.78 (74) | 5.11 |
|  | 15.5 | **1.57 (29)** | - | 3.78 (71) | 5.35 |
|  | 18 | **1.76 (32)** | - | 3.78 (68) | 5.54 |
|  | 21.5 | **2.04 (35)** | - | 3.78 (65) | 5.82 |
|  | 24 | **2.25 (37)** | - | 3.78 (63) | 6.03 |
|  | 27.5 | **2.57 (40)** | - | 3.78 (60) | 6.35 |
|  | 30 | **2.81 (43)** | - | 3.78 (57) | 6.59 |
| DunedinPACE  (ADE model with moderated *d* effect) | 9.5 | 0.002 (32) | **0.001 (10)** | 0.003 (58) | 0.005 |
|  | 12 | 0.002 (30) | **0.001 (13)** | 0.003 (57) | 0.006 |
|  | 15.5 | 0.002 (29) | **0.001 (17)** | 0.003 (54) | 0.006 |
|  | 18 | 0.002 (28) | **0.001 (20)** | 0.003 (52) | 0.006 |
|  | 21.5 | 0.002 (27) | **0.001 (24)** | 0.003 (49) | 0.006 |
|  | 24 | 0.002 (26) | **0.002 (27)** | 0.003 (47) | 0.007 |
|  | 27.5 | 0.002 (24) | **0.002 (31)** | 0.003 (45) | 0.007 |
|  | 30 | 0.002 (23) | **0.002 (34)** | 0.003 (43) | 0.007 |

*Note.* The estimates are derived from the best-fitting univariate models with age moderation of genetic and environmental components of the four epigenetic aging measures, which are depicted in parentheses. *A* (%), component due to additive genetic factors; *D* (%), component due to non-additive genetic factors; *C* (%), component due to shared environmental factors; *E* (%), component due to individually-unique environmental factors including measurement error; *V*, total variance; Accel., Acceleration. Components that are significantly moderated by age are shown in bold.
